# Generative 3D Spatial Reconstruction of Cellular Organization from Single-Cell Expression

**DOI:** 10.64898/2026.11.05.764656

**Authors:** Ehtesamul Azim, Muhtasim Noor Alif, Tae Hyun Hwang, Hao Zheng, Wei Zhang

## Abstract

Single-cell RNA sequencing profiles cellular states at scale but removes their three-dimensional (3D) spatial organization. Reconstructing this organization requires recovering not only individual cell positions but also the physical size, geometry, and neighborhoods of an entire tissue. Existing methods either map cells onto measured spatial references or reconstruct two-dimensional sections independently, leaving the joint reconstruction of a new 3D specimen unresolved. Moreover, predicting the average plausible position of each cell can reduce per-cell error while compressing tissue geometry. We introduce **ORBIS**, a generative framework that jointly reconstructs whole 3D specimens from single-cell gene expression without requiring a spatial reference at inference. ORBIS learns spatial organization through flow matching in a shared physical coordinate frame, using training views sampled from sparse serial sections, tissue blocks, and whole specimens. At inference, a shared spatial state is iteratively updated through dynamically regrouped windows, enabling the generation of millions of cell positions without independently reconstructing and stitching tissue regions. We evaluate physical size, cell-population shape, per-cell placement, and neighborhood organization separately to distinguish accurate tissue geometry from low per-cell error. Across eight held-out specimens spanning whole mouse brains, thick brain blocks, and *Drosophila* embryos, ORBIS achieves lower cell-population shape error than all five evaluated baselines on every test and reconstructs specimens containing up to 3.7 million cells. On a single GPU, ORBIS samples million-cell brains in 0.9–2.8 minutes, approximately 38 times faster than the sampler of LUNA-3D, our 3D extension of the state-of-the-art generative spatial reconstruction method LUNA. On two held-out brain tests, sampling reduces size error by 3.3-fold and shape error by 1.8-fold compared with a single-call variant. On an independent dissociated single-cell brain atlas, ORBIS returns on average 37.9% of each region’s cells to their dissection region across 23 regions, compared with 28.5% for LUNA-3D.

Checkpoints: https://doi.org/10.5281/zenodo.23925991

## 1 Introduction

Cells function within three-dimensional (3D) tissues, where their states and functions are shaped by neighboring cells, anatomical regions, tissue boundaries, and molecular gradients [1–8]. Single-cell RNA sequencing (scRNA-seq) profiles cellular states at large scale [9, 10], but tissue dissociation removes the spatial positions of the measured cells. Spatial transcriptomics preserves spatial information [11–14], but most technologies measure thin two-dimensional (2D) sections, sparse serial sections, or tissue blocks with limited depth [8, 15–19]. As a result, many single-cell datasets lack the 3D organization needed to study cell–cell interactions, tissue patterning, and molecular gradients [20–23]. Computationally reconstructing dissociated cells in 3D can therefore connect large-scale single-cell profiling with the spatial organization in which cells function.

From a computational perspective, this problem is much harder than predicting one additional coordinate. Given the expression profiles of an unordered set of cells, the goal is to recover their **joint 3D organization**. Gene expression does not uniquely determine position, because cells with similar molecular states can appear in multiple possible locations [24, 25]. At the same time, a useful reconstruction must preserve both local neighborhoods and global tissue properties, including physical size, anatomical shape, distances between cell populations, and organization across tissue depth. These requirements must also be met for specimens containing millions of cells, far more than a neural network can process at once. Thus, 3D reconstruction requires modeling the organization of the whole tissue rather than predicting each cell independently.

Existing computational approaches address only parts of this problem. Most methods are reference-based. Deconvolution methods estimate cell-type composition in measured spatial locations [26–28], while cell-to-space methods map dissociated cells to locations or coordinate systems provided by a spatial reference [25, 29–38]. These methods can place cells accurately when a closely matched spatial reference is available. However, the candidate locations and coordinate system are defined by the reference, so the resulting tissue geometry is largely inherited from that reference rather than reconstructed from the expression profiles of the new specimen. Methods for 3D spatial transcriptomics address a different problem by aligning measured sections into volumes or inferring expression between sections [39–43]. These methods start from cells whose spatial positions are already observed. Neither setting directly addresses reconstructing the 3D organization of a new dissociated specimen from expression alone.

Recent generative methods take a different approach by learning spatial organization from spatially resolved training data and reconstructing cells without a spatial reference at inference [44–46]. However, existing methods reconstruct one 2D section at a time. Extending them to a whole 3D specimen creates two major problems. First, independently reconstructed sections or cell subsets do not share the same physical frame. Their orientations, spacing, scale, and relative positions are not guaranteed to agree, so they cannot simply be combined into a coherent tissue. Second, existing methods often focus on per-cell accuracy. Because expression does not fully determine position, minimizing coordinate error tends to place a cell near the average of its possible locations [47]. This can reduce spatial variation, shrink the reconstructed tissue, compress cell populations, increase neighborhood density, and change distances between anatomical regions. These errors are especially important in 3D, where tissue size and spatial relationships are biologically meaningful.

We therefore formulate 3D spatial reconstruction as **generating an entire specimen jointly**. Instead of predicting one best position for each cell, we aim to sample one plausible configuration of all cells based on their expression. We introduce **ORBIS** (**<u>O</u>**rganism-scale spatial **<u>R</u>**econstruction **<u>B</u>**y **<u>I</u>**terative **<u>S</u>**ampling), a flow-matching framework [48, 49] that learns this distribution from spatially resolved reference specimens and reconstructs a new specimen, from whole organs to whole organisms, by iteratively sampling the positions of all its cells from their expression alone. Training specimens are represented in a shared physical coordinate frame, allowing the model to learn tissue scale, orientation, and relationships across depth. Randomly sampled whole-specimen, single-section, and multi-section block views allow the same framework to learn from sparse serial sections, continuous tissue blocks, and densely sectioned organisms.

To scale it to millions of cells, ORBIS maintains one shared spatial state while processing cells in windows that are regrouped across flow steps. This allows information to propagate across the specimen while preserving one consistent 3D configuration. ORBIS therefore reconstructs specimens with up to 3.7 million cells without generating and stitching separate pieces. A final neighborhood-consistent reseating step improves local organization while preserving tissue geometry. Our 50 steps sample a brain of millions of cells in under three minutes on one GPU, whereas the 1,000-step sampler of LUNA [45], extended to 3D (LUNA-3D), takes 36 to 108 minutes, about 38 times longer; with reseating, the full pipeline remains 1.2–3.2 times faster.

We evaluate ORBIS on eight held-out specimens, including whole mouse brains [15, 16], thick brain blocks [8], and *Drosophila* embryos [50]. ORBIS better recovers tissue and cell-population geometry than all evaluated baselines and preserves tissue size where LUNA-3D contracts several specimens by 15–66% along at least one axis. ORBIS also better preserves cellular neighborhoods, cell-type contacts, positional order along the *Drosophila* gut, and regional organization in a whole-brain single-cell atlas, where it returns on average 37.9% of a region’s cells to it, against 28.5% for LUNA-3D.

Our key contributions are as follows:

- *Whole-specimen generation:* We formulate 3D spatial reconstruction as generating one joint configuration of all cells rather than independently predicting their most likely positions, and show why point prediction can compress tissue geometry. To our knowledge, ORBIS is the first method to jointly generate whole-specimen 3D configurations of dissociated cells without a spatial reference at inference.
- *Scalable 3D reconstruction:* ORBIS combines flow matching in a shared physical frame, multiple training views, shared-state windowed sampling, and neighborhood reseating to reconstruct specimens containing millions of cells from expression alone.
- *Biologically grounded evaluation:* We propose an evaluation of 3D reconstruction that scores physical size, cell-population shape, per-cell placement, and neighborhood organization separately. Across three distinct 3D tissue settings, ORBIS improves tissue and cell-population geometry while better preserving neighborhoods, anatomical organization, and, in brains and tissue blocks, spatial gene-expression patterns.

## 2 Related Work

### Reference-based reconstruction

Most methods place dissociated cells by matching them to measured reference spots or cells through learned or optimal-transport correspondences [25, 31, 30, 33, 29, 32], or by learning coordinate or spatial-affinity predictors from a reference [36, 34, 35, 37]. REMAP uses reference-derived neighborhood gene–gene covariance, including for 3D mouse-brain references [38]. These methods inherit locations or coordinate frames from the reference rather than generating the geometry of a new specimen. Other 3D spatial-transcriptomics methods align measured sections or infer expression between them [39–43], while MIMYR generates cells in missing regions adjacent to measured sections [51]. Unlike ORBIS, these approaches rely on spatially observed cells at inference.

### Reference-free generative reconstruction

Recent methods learn spatial organization from training data and reconstruct cells from expression alone. GEARS stitches patchwise pairwise distances into a 2D layout through a landmark eigendecomposition [44]. LUNA trains an expression-conditioned diffusion model for 2D sections, with noisy positions entering attention through directions from the section centroid [45] (Supplementary Section S8). G2T builds on LUNA with flow matching, recovering coordinates from learned embedding distances through multidimensional scaling [46]. All three reconstruct 2D sections rather than a whole 3D specimen in a shared physical frame. The *N* × *N* matrix that G2T’s scaling decomposes needs about 6 TB for a whole brain of 1.2 million cells, far beyond our largest GPU (96 GB), and GEARS relies on a similar decomposition and on stitching, so we extend LUNA to 3D as the representative baseline.

### Generation versus averaging

Flow matching [48, 49] and diffusion models [52] generate samples through iterative updates, with diffusion transformers [53] as denoisers. MultiDiffusion maintains a shared generative state by combining predictions from overlapping windows [54]; our sampler instead regroups cells into new windows at each step. Sampling also addresses a fundamental trade-off: squared-error prediction averages plausible outcomes and reduces their variation [55], whereas an exact conditional sample preserves the output distribution but incurs twice the expected squared error of the conditional mean [47]. Thus, optimizing individual-cell accuracy can compress tissue geometry, motivating our objective of sampling a coherent whole specimen rather than averaging its possible configurations.

## 3 Problem Statement

We reconstruct the three-dimensional positions of dissociated cells from gene expression alone, using *R* spatially resolved reference specimens for supervision. Reference specimen *r* contains expression 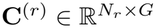 for *N*_*r*_ cells and *G* genes, and corresponding coordinates 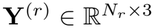, with rows **c**_*i*_ and **y**_*i*_ for cell *i*. References of the same tissue share a physical coordinate frame preserving section order, spacing, orientation, and scale, despite incomplete spatial coverage. At inference, given only the expression **C** ∈ ℝ^*N×G*^ of a new specimen, we jointly generate positions **Ŷ** ∈ ℝ^*N×*3^. Since expression does not determine absolute pose, evaluation permits rigid alignment but not rescaling.

We formulate reconstruction as conditional generation, seeking **Ŷ** ∼ *p*(**Y** | **C**). Since cells are unordered, the distribution must satisfy *p*(**PY** | **PC**) = *p*(**Y** | **C**) for any permutation matrix **P**. Under squared error, the optimal point estimate for a randomly selected cell is the conditional mean 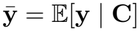. The law of total covariance gives

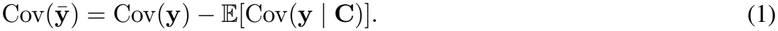

Thus, predicting conditional means reduces spatial variation by averaging uncertain positions. Exact conditional sampling instead preserves marginal covariance, Cov(**ŷ**) = Cov(**y**), at the cost of twice the expected squared error of the conditional mean [47]. Our goal is therefore to generate a plausible whole-specimen configuration rather than an averaged layout.

## 4 Method

### Framework overview

Figure 1 illustrates the training and inference pipelines of ORBIS. During training, we sample whole-specimen, section, or block views from spatial references and interpolate their 3D coordinates with Gaussian noise at a sampled flow time. A diffusion transformer (DiT) embeds each cell’s expression and noisy position, conditions its attention blocks on flow time and view type, and predicts clean coordinates under an MSE objective. During inference, we initialize the positions of all query cells from Gaussian noise and partition them into randomly regrouped windows at each sampling step. The DiT predicts clean coordinates within each window, which are assembled into the shared state and updated through 50 Euler steps. Finally, neighborhood reseating reassigns cell identities among fixed sampled positions to refine local composition.

**Figure 1:**
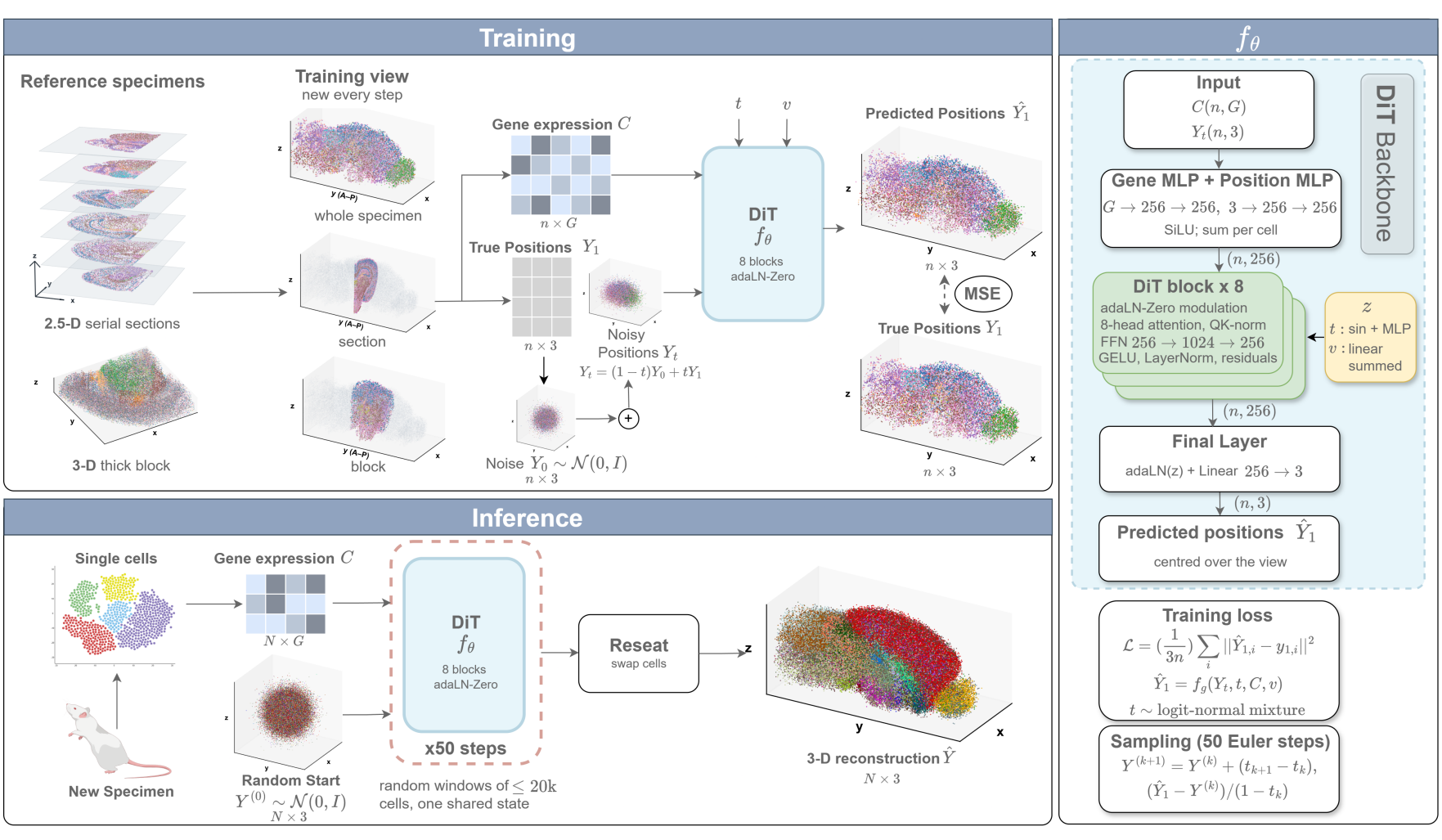
Overview of **ORBIS. Top left:** Training views sampled from spatially resolved reference specimens for flow-matching supervision. **Bottom left:** Whole-specimen reconstruction through shared-state iterative sampling and neighborhood reseating. **Right:** Diffusion transformer (DiT) architecture, training objective, and Euler sampling update.

### 4.1 Flow matching in a shared physical frame

We model the conditional distribution of three-dimensional cell positions given gene expression using flow matching [48, 49], which learns a time-dependent velocity field that transforms Gaussian noise into a spatial configuration. Reference specimens of each tissue share a physical coordinate frame preserving section order, spacing, and relative orientation. We divide coordinates by one length scale shared across specimens and axes, preserving relative size and proportions, and center each training view to remove translation without changing its orientation or scale.

Let **Y**_1_ denote the reference coordinates and **Y**_0_ independent standard normal vectors centered over the view. Along a linear path from Gaussian source (*t* = 0) to reference (*t* = 1), the network predicts the reference endpoint, conditioned on expression **C**, time *t*, and view label *v* (Section 4.2). The resulting path, endpoint prediction, and velocity are

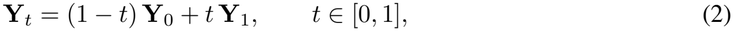

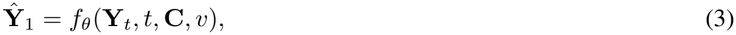

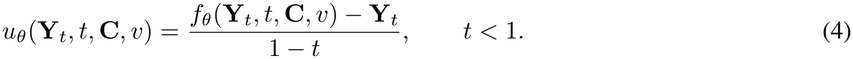

This endpoint parameterization permits direct coordinate supervision (Section 4.4). At inference, we integrate this field from Gaussian noise using the whole-specimen sampler in Section 4.5.

### 4.2 Training views

At every training step, we resample a subset of reference cells, retaining the specimen’s orientation and physical scale and centering the subset as in Section 4.1. This exposes the model to varying cell combinations and supports datasets with different spatial coverage and measurement density. We use three view types: a *whole-specimen* view samples cells uniformly throughout a specimen, providing global context and matching inference windows; a *section* view contains the cells of one measured section, capturing dense in-plane structure; and a *block* view samples consecutive sections, retaining spacing and orientation across depth. Sectioned datasets use all three types, whereas continuous tissue blocks use whole-specimen views only.

The label *v* identifies the view type and, for section and block views, the cutting plane, distinguishing planar from volumetric observations. At inference, every window uses the whole-specimen label. During view construction, we randomly thin transcript counts before log transformation to accommodate differences in sequencing depth. For bilaterally symmetric embryos, we also fold coordinates across the body midline to account for ambiguous left–right placement; the same fold is applied to the ground truth and all reconstructions during evaluation. Further details are in Supplementary Section S1.2.

### 4.3 Set denoiser

We parameterize *f*_*θ*_ with a diffusion transformer [53] operating on an unordered set of cell tokens. Full self-attention lets each prediction depend on the expression and current coordinates of other cells in the view. With no token-order positional encoding, the network is permutation-equivariant: permuting its input cells permutes its outputs accordingly.

For cell *i*, the input token combines expression and interpolated position, while time and view type provide a separate conditioning vector:

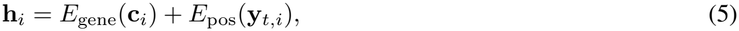

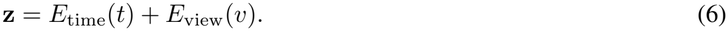

Because **h**_*i*_ includes **y**_*t,i*_, predictions adapt to the evolving spatial configuration rather than depending on expression alone. Following adaLN-Zero [53], **z** modulates normalization layers and residual gates within each transformer block. A final projection produces three coordinates per cell, and predictions are centered over the view. Architecture and embedding details appear in Supplementary Section S1.3.

### 4.4 Coordinate supervision

An isolated two-dimensional section may be reconstructed up to translation, rotation, and reflection, making pairwise-distance supervision suitable because it leaves pose unspecified. Our three-dimensional formulation instead requires a shared physical frame: views retain anatomical orientation and inter-section spacing, while flow steps update coordinates in that frame. We therefore supervise coordinates directly. For a view of *n* cells, the objective is

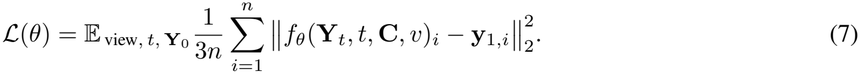

Predictions and targets are independently centered over the view, removing translation while preserving orientation and physical scale. For fixed interpolated coordinates, time, and conditioning, the squared-error minimizer is the conditional mean of the reference coordinates; inserting it into Eq. (4) gives the conditional expected velocity along the path.

We draw *t* from an equal mixture of two logit-normal distributions: one covers strongly perturbed coordinates and the broader path, following rectified-flow training [56]; the other emphasizes refinement near the reference endpoint, using the EDM noise-level distribution [57] mapped onto our interpolation. Parameters and optimization details are in Supplementary Section S1.4.

### 4.5 Whole-specimen sampling and reseating

#### Whole-specimen sampling

A query may contain millions of cells, exceeding the network’s window capacity. Generating chunks independently would give them separate spatial states that may disagree after stitching. Instead, following MultiDiffusion [54], we maintain one shared coordinate state for all *N* cells; unlike its fixed windows, we regroup the processing windows at every step. Starting from noise **Y**^(0)^, at step *k* we randomly partition cells into windows *W* of training-view size, sampled across the specimen rather than as spatial tiles. We center each window by its mean ***μ***_*W*_, predict its endpoint under the whole-specimen label, restore that mean, and advance the assembled state by Euler integration:

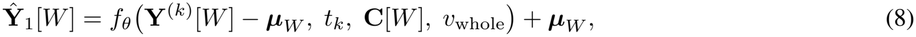

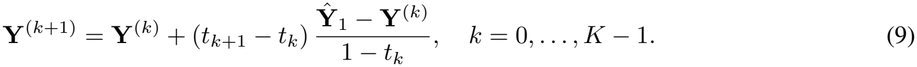

Redrawing windows changes which cells interact at each step while retaining one specimen-wide state. The final **Ŷ** = **Y**^(*K*)^ is a stochastic tissue realization, rather than the average layout of a single network call (Section 3). We use EDM-spaced steps [57] without guidance (Supplementary Section S1.5).

#### Neighborhood reseating

Sampling recovers overall geometry and population distributions, but cells with similar expression may remain ambiguous despite having different neighborhoods. We therefore keep sampled positions fixed and refine which cell occupies each site, motivated by transporting a point estimate onto a learned distribution [58]. A neighborhood reader trained on reference specimens predicts **q**_*c*_, the expected neighborhood composition of cell *c* over soft expression groups. We score an assignment *π* of cells to sampled sites **s**_*a*_ by

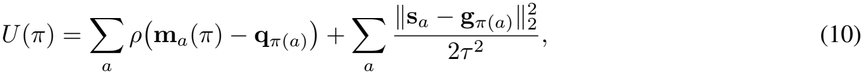

where **m**_*a*_(*π*) is the neighborhood composition at site *a, ρ* a robust penalty, **g**_*c*_ the network’s single-call position estimate, and *τ* the per-axis root-mean-square difference between these estimates and the sample. The terms favor neighborhood compatibility and proximity to predicted positions; an additional term penalizes changes to each group’s sampled center and spread. We lower *U* through Metropolis swaps between nearby sites [59], limiting each cell’s displacement from its original site to *τ* . Since sites remain fixed, the specimen retains exactly the sampled positions, and thus its overall shape and size (Supplementary Section S1.6).

## 5 Experiments

### 5.1 Experimental setup

#### Datasets

We evaluate on three tissue geometries, each probing a distinct generalization regime: sparsely sectioned whole mouse brains profiled with MERFISH (1.2–2.8 million cells per test brain) [15], tested on held-out animals and on a held-out cutting orientation, together with a whole brain from another laboratory and platform (3.7 million cells) [16]; continuous thick blocks of mouse brain profiled with Deep-STARmap, each reconstructed by a model trained on the other [8]; and densely sectioned whole *Drosophila* embryos from Flysta3D, with two embryos held out [50]. Dataset construction, splits, coordinate frames and preprocessing are in Supplementary Section S2.

#### Baselines

We compare against five baselines spanning generative reconstruction (LUNA [45]) and reference-based reconstruction (REMAP [38], CeLEry [36], Tangram [25] and novoSpaRc [31]). Each is run from its released code, with two-dimensional methods extended to three-dimensional coordinates (suffix -3D). All methods learn from the same training specimens and are scored by the same evaluation code; adaption details in Supplementary Section S3.

#### Evaluation

Because the orientation of a specimen cannot be read from expression, every reconstruction is rigidly aligned to the truth before scoring. Eight metrics in two groups then score the specimen and its cell types, namely physical size (*size error*), the shape of each cell-type population independently of size (*shape error, W*_2_), the overlap of each population with its true counterpart (*F-score*) and where each population lies (*type-centre error*), and the cells and their neighborhoods, namely the position of each cell relative to a random position of its own cell type (*placement*), neighborhood organization (*niche NMI*), which cell types are in contact (*type contacts*) and the spatial patterns of genes (*Moran’s I*). The per-cell distance correlation and RSSD used for two-dimensional reconstructions [45, 46] favor predicting each cell’s average position, and the first does not change when a reconstruction shrinks; we report them, with this analysis, in Supplementary Section S4. For every method we report the mean and standard deviation over three runs with different random seeds. Formal definitions and the full protocol are in Supplementary Section S4.

### 5.2 Results

#### Main results

Our model recovers the shape of the cell-type populations more closely than every baseline in all three geometries (Table 1), and its shape error is lower than that of each of LUNA-3D’s three trainings on all eight tests (Supplementary Tables S4–S6). On the whole brains it averages 198 *μ*m against 431 *μ*m for LUNA-3D and 673–883 *μ*m for the reference-based methods; the gain is largest on mouse 1 (169 versus 523 *μ*m). Each population also lies where it should, with the best type-centre error and F-score of all methods in every geometry (type-centre error 182/213/10 *μ*m versus 415/263/17 *μ*m for LUNA-3D). They also keep the tissue’s physical size: size error averages 0.041 on the brains and 0.077 on the blocks, against 0.105 and 0.264 for LUNA-3D, whose mice 1 and 2 and blocks are 15–66% too small along at least one axis in each of its trainings. On the brains, LUNA-3D also squeezes 8–23% of the cell types below 0.8 of their true spread even after size matching, against 1–3% for our model. Tangram-3D has a small size error only because it copies the positions of training cells; its placement is near or below zero on every test.

**Table 1:** Mean over the tests of each geometry (per test: Supplementary Tables S4–S6). Every method: mean _±SD_ over 3 runs with different seeds. **Bold**/<u>underline</u>/*italic*: 1st/2nd/3rd.

| Data | Method | Specimen and cell types |  |  |  | Cells and neighborhoods |  |  |  |
|---|---|---|---|---|---|---|---|---|---|
|  |  | Size error ↓ | Shape *W*<sub>2</sub> ↓<br>(*µ*m) | F-score ↑ | Type<br>centre ↓<br>(*µ*m) | Placement ↑ | Niche<br>NMI ↑ | Type<br>contacts ↑ | Moran's *I* ↑ |
| **Whole brains** | LUNA-3D | *.105*<sub>±.015</sub> | <u>431</u><sub>±185</sub> | <u>.496</u><sub>±.089</sub> | <u>415</u><sub>±182</sub> | <u>.450</u><sub>±.128</sub> | <u>.429</u><sub>±.041</sub> | <u>.913</u><sub>±.017</sub> | <u>.778</u><sub>±.086</sub> |
|  | REMAP | .289<sub>±.014</sub> | 878<sub>±4</sub> | .387<sub>±.002</sub> | 888<sub>±9</sub> | .312<sub>±.003</sub> | *.407*<sub>±.003</sub> | *.857*<sub>±.002</sub> | .624<sub>±.013</sub> |
|  | CeLEry-3D | .256<sub>±.0002</sub> | *673*<sub>±10</sub> | .398<sub>±.003</sub> | *628*<sub>±19</sub> | *.385*<sub>±.002</sub> | .384<sub>±.005</sub> | .816<sub>±.010</sub> | *.635*<sub>±.015</sub> |
|  | Tangram-3D | **.038**<sub>±.001</sub> | 883<sub>±2</sub> | .166<sub>±.001</sub> | 746<sub>±4</sub> | −.093<sub>±.002</sub> | .230<sub>±.001</sub> | .581<sub>±.002</sub> | .380<sub>±.003</sub> |
|  | novoSpaRc-3D | .282<sub>±.004</sub> | 816<sub>±6</sub> | *.422*<sub>±.001</sub> | 752<sub>±11</sub> | .199<sub>±.001</sub> | .280<sub>±.003</sub> | .816<sub>±.003</sub> | .431<sub>±.011</sub> |
|  | **Ours** | <u>.041</u><sub>±.002</sub> | **198**<sub>±2</sub> | **.605**<sub>±.004</sub> | **182**<sub>±4</sub> | **.467**<sub>±.001</sub> | **.479**<sub>±.002</sub> | **.954**<sub>±.001</sub> | **.892**<sub>±.004</sub> |
| **Thick blocks** | LUNA-3D | *.264*<sub>±.026</sub> | *313*<sub>±78</sub> | *.765*<sub>±.083</sub> | <u>263</u><sub>±38</sub> | *.251*<sub>±.136</sub> | .224<sub>±.077</sub> | *.895*<sub>±.036</sub> | .711<sub>±.196</sub> |
|  | REMAP | .801<sub>±.372</sub> | 691<sub>±48</sub> | .431<sub>±.035</sub> | 617<sub>±40</sub> | −.236<sub>±.233</sub> | .111<sub>±.023</sub> | .815<sub>±.003</sub> | .620<sub>±.064</sub> |
|  | CeLEry-3D | .345<sub>±.012</sub> | <u>302</u><sub>±3</sub> | <u>.768</u><sub>±.002</sub> | *299*<sub>±13</sub> | <u>.395</u><sub>±.004</sub> | <u>.294</u><sub>±.002</sub> | <u>.912</u><sub>±.006</sub> | <u>.737</u><sub>±.059</sub> |
|  | Tangram-3D | <u>.084</u><sub>±.0002</sub> | 450<sub>±2</sub> | .682<sub>±.005</sub> | 445<sub>±3</sub> | −.196<sub>±.004</sub> | .079<sub>±.001</sub> | .805<sub>±.002</sub> | *.722*<sub>±.013</sub> |
|  | novoSpaRc-3D | .468<sub>±.004</sub> | 386<sub>±4</sub> | .711<sub>±.001</sub> | 394<sub>±9</sub> | .242<sub>±.002</sub> | *.232*<sub>±.005</sub> | .883<sub>±.002</sub> | .613<sub>±.038</sub> |
|  | **Ours** | **.077**<sub>±.020</sub> | **211**<sub>±0.9</sub> | **.862**<sub>±.004</sub> | **213**<sub>±5</sub> | **.445**<sub>±.005</sub> | **.373**<sub>±.002</sub> | **.973**<sub>±.001</sub> | **.926**<sub>±.0002</sub> |
| **Embryos** | LUNA-3D | **.102**<sub>±.025</sub> | <u>18.5</u><sub>±3.2</sub> | <u>.981</u><sub>±.004</sub> | <u>16.7</u><sub>±4.4</sub> | <u>.253</u><sub>±.045</sub> | <u>.226</u><sub>±.018</sub> | <u>.921</u><sub>±.011</sub> | **.857**<sub>±.048</sub> |
|  | REMAP | .516<sub>±.094</sub> | 56.0<sub>±2.7</sub> | .825<sub>±.005</sub> | 55.8<sub>±1.2</sub> | −.177<sub>±.039</sub> | .030<sub>±.004</sub> | .678<sub>±.014</sub> | .203<sub>±.108</sub> |
|  | CeLEry-3D | .587<sub>±.014</sub> | *32.6*<sub>±0.5</sub> | *.898*<sub>±.003</sub> | *27.6*<sub>±0.3</sub> | *.168*<sub>±.002</sub> | *.198*<sub>±.005</sub> | *.817*<sub>±.005</sub> | .640<sub>±.018</sub> |
|  | Tangram-3D | *.124*<sub>±.010</sub> | 42.8<sub>±0.3</sub> | .897<sub>±.003</sub> | 41.5<sub>±0.5</sub> | −.137<sub>±.009</sub> | .032<sub>±.001</sub> | .694<sub>±.009</sub> | .525<sub>±.012</sub> |
|  | novoSpaRc-3D | .563<sub>±.005</sub> | 32.8<sub>±0.4</sub> | .893<sub>±.002</sub> | 29.4<sub>±0.6</sub> | .099<sub>±.004</sub> | .136<sub>±.004</sub> | .789<sub>±.003</sub> | *.673*<sub>±.015</sub> |
|  | **Ours** | <u>.107</u><sub>±.003</sub> | **11.5**<sub>±0.2</sub> | **.990**<sub>±.0001</sub> | **10.3**<sub>±0.3</sub> | **.400**<sub>±.003</sub> | **.253**<sub>±.001</sub> | **.931**<sub>±.001</sub> | <u>.747</u><sub>±.014</sub> |

Our model also places individual cells more accurately than every baseline in all three geometries (placement 0.467/0.445/0.400 versus at most 0.450/0.395/0.253). Per-cell accuracy alone rewards predicting each cell’s average plausible position, which also narrows the populations (Section 2). A single call of our network approximates this average: its per-cell error is lower than LUNA-3D’s on all eight tests, but its size and shape errors are higher than our sample’s on all eight (Supplementary Section S7, Supplementary Table S7). Sampling also tells which placements to trust: the spread of four samples predicts the error of each cell (Spearman 0.51–0.56 on the held-out mice), and the quarter of cells with the lowest spread have a median error of 0.84–0.89 mm, against 1.46–1.56 mm for all cells (Supplementary Figure S1). After reseating, niche organization agrees with the truth more closely for our model than for any baseline in all three geometries and on seven of the eight tests. So do the contacts between cell types (correlation 0.954/0.973/0.931 versus at most 0.913/0.912/0.921) and, on the brains and blocks, the spatial patterns of genes (Moran’s I agreement 0.892/0.926 versus at most 0.778/0.737).

#### Ablation study

Table 2 removes one design choice at a time from our final model (mean over mice 2 and 3; every variant on every test in Supplementary Section S6). Our sampler draws all cells of a specimen as one shared state over 50 steps, splitting them at every step into a fresh random partition of 20,000-cell windows. *Single call* replaces the 50 steps by one network call from pure noise, which approximates each cell’s average position: it places single cells more accurately (placement 0.560 versus 0.454), but its size error is 3.3 times and its shape error 1.8 times that of the sample (363 versus 205 *μ*m) and its niche agreement is 13% lower, the trade-off described in Section 2. *Independent chunks* samples separate chunks of about 19,000 cells, each with its own state, and aligns them afterwards on shared cells; it is worse on all four measures, with 7.3 times the size error, 2.1 times the shape error, and 30% and 17% lower placement and niche agreement. A new partition at every step matters where the tissue is continuous: with fixed windows, the size error of the two blocks doubles and triples, while the mice are unaffected. Two training choices matter as much. We train the network to predict cell positions in micrometres. *Distance loss* trains it instead on pairwise distances, which leave the frame free: each window comes out in its own pose, so the model is sampled in independent chunks, and its shape error is 2.5 times ours and still 16% above our own model sampled the same way. *Unit-box coordinates* rescales each training view to a unit box and so discards physical scale: the shape is kept but not the size (6.2 times the size error), and placement drops by 38%. Randomly thinning the transcript counts of training cells prepares the model for shallow profiles: given a quarter of the counts, a model trained without thinning has a shape error of 536 instead of 381 *μ*m (mouse 2). Finally, *w/o reseating* omits the last step of our method, which exchanges the positions of pairs of sampled cells when this makes their neighborhoods agree better with those predicted from their expression, without moving any cell further than the model’s uncertainty; reseating raises niche agreement from 0.416 to 0.491 and leaves size and shape unchanged.

**Table 2:**
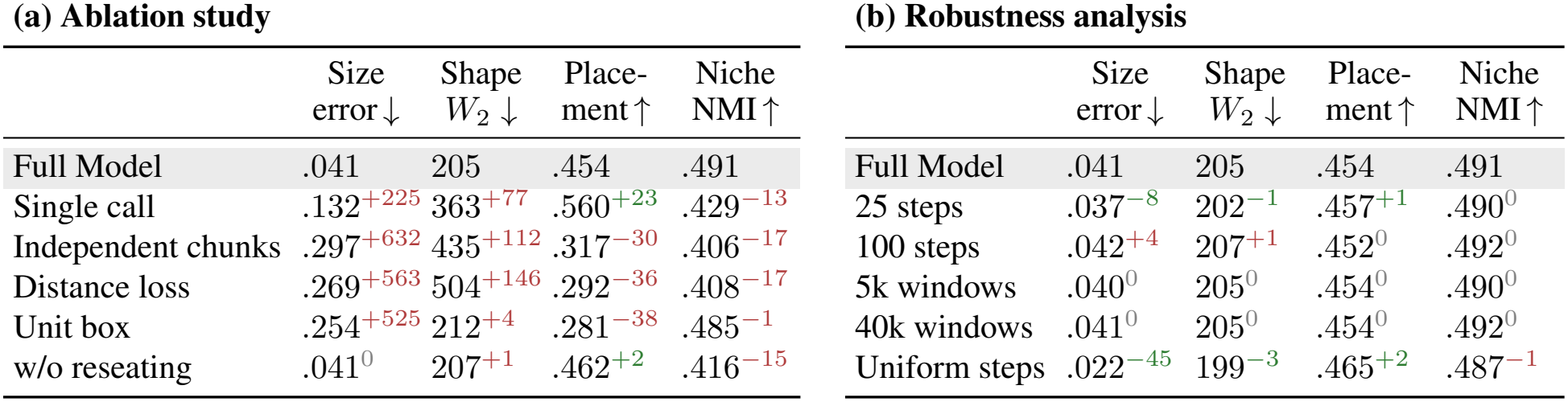
Ablation study (a) and robustness analysis (b), mice 2 and 3, training seed 0. Superscripts: % change from the Full Model, green better, red worse.

| (a) Ablation study |  |  |  |  | (b) Robustness analysis |  |  |  |  |
|---|---|---|---|---|---|---|---|---|---|
|  | Size<br>error ↓ | Shape<br>*W*<sub>2</sub> ↓ | Place-<br>ment ↑ | Niche<br>NMI ↑ |  | Size<br>error ↓ | Shape<br>*W*<sub>2</sub> ↓ | Place-<br>ment ↑ | Niche<br>NMI ↑ |
| Full Model | .041 | 205 | .454 | .491 | Full Model | .041 | 205 | .454 | .491 |
| Single call | .132<sup>+225</sup> | 363<sup>+77</sup> | .560<sup>+23</sup> | .429<sup>−13</sup> | 25 steps | .037<sup>−8</sup> | 202<sup>−1</sup> | .457<sup>+1</sup> | .490<sup>0</sup> |
| Independent chunks | .297<sup>+632</sup> | 435<sup>+112</sup> | .317<sup>−30</sup> | .406<sup>−17</sup> | 100 steps | .042<sup>+4</sup> | 207<sup>+1</sup> | .452<sup>0</sup> | .492<sup>0</sup> |
| Distance loss | .269<sup>+563</sup> | 504<sup>+146</sup> | .292<sup>−36</sup> | .408<sup>−17</sup> | 5k windows | .040<sup>0</sup> | 205<sup>0</sup> | .454<sup>0</sup> | .490<sup>0</sup> |
| Unit box | .254<sup>+525</sup> | 212<sup>+4</sup> | .281<sup>−38</sup> | .485<sup>−1</sup> | 40k windows | .041<sup>0</sup> | 205<sup>0</sup> | .454<sup>0</sup> | .492<sup>0</sup> |
| w/o reseating | .041<sup>0</sup> | 207<sup>+1</sup> | .462<sup>+2</sup> | .416<sup>−15</sup> | Uniform steps | .022<sup>−45</sup> | 199<sup>−3</sup> | .465<sup>+2</sup> | .487<sup>−1</sup> |

#### Robustness analysis

The sampler is robust to its settings (Table 2): half or twice the steps, windows of 5,000 or 40,000 cells instead of 20,000, or uniformly spaced steps worsen no measure by more than 4%; the spacing mainly moves the size error, which uniform steps nearly halve on mice 2 and 3 but raise on the other six tests. Nor does the model need every cell: a random 10% of the cells, placed alone, is reconstructed as well as among all the others, with no measure worse by more than 4% on any of the eight tests (both in Supplementary Table S8). Training is equally stable: a network with 3.3 times the parameters changes the shape error of mouse 2 by 2%, and the standard deviation across our three trainings is at most 2% of the shape error (Table 1).

#### Training and sampling cost

Table 3 gives the cost of our model and LUNA-3D on one GPU. We sample a whole mouse brain of 1.2 to 3.7 million cells in 0.9 to 2.8 minutes against 36 to 108 for LUNA-3D, about 38 times faster on all eight tests. Flow matching reaches the layout in 50 steps where LUNA-3D’s diffusion sampler runs 1,000, and each of our steps costs about half as much per cell. Both work on sets of fixed size (our windows of 20,000 cells, its graphs of about 5,000), so sampling time grows linearly with the number of cells (log-log slope 1.00 for both; Supplementary Section S13). Reseating adds 2 to 77 minutes on a smaller GPU, where sampling and reseating together remain 1.2 to 3.2 times faster than LUNA-3D’s sampling alone. Training scales differently: an epoch of ours holds 246 views at any data size, so its time does not grow with the training cells (12 to 18 seconds from 0.2 to 5.8 million cells, slope −0.03), while a LUNA-3D epoch is one pass over every cell (1 to 22 seconds, slope 0.86). A full training on a whole brain takes 1.2 to 1.8 hours with ours and 1.1 to 1.9 with LUNA-3D. On the blocks and embryos we take far more updates, 123 per epoch against 5 to 8, and so more time (1.3 to 1.7 against 0.1 to 0.2 hours); per update ours is 1.2 to 1.9 times faster on every test, and training LUNA-3D for our number of updates does not improve it (Supplementary Section S13).

**Table 3:** Training and sampling cost on one GPU (NVIDIA RTX PRO 6000). Training: ours 360 epochs (embryos 300), LUNA-3D 300 (blocks 400). Sampling: every test cell, 50 steps for ours and 1,000 for LUNA-3D.

|  |  | Mouse 3 | Mouse 2 | Mouse 1 | Allen | Block A | Block B | Embryo c | Embryo d |
|---|---|---|---|---|---|---|---|---|---|
| **Training** | Cells (M) | 4.07 | 4.60 | 2.98 | 5.82 | 0.20 | 0.18 | 0.29 | 0.29 |
|  | **Ours** (h) | 1.71 | 1.76 | 1.77 | 1.19 | 1.71 | 1.70 | 1.33 | 1.33 |
|  | LUNA-3D (h) | 1.43 | 1.63 | 1.05 | 1.86 | 0.13 | 0.12 | 0.17 | 0.17 |
| **Sampling** | Cells (M) | 1.59 | 1.23 | 2.85 | 3.74 | 0.18 | 0.20 | 0.026 | 0.040 |
|  | **Ours** (min) | 1.2 | 0.93 | 2.2 | 2.8 | 0.14 | 0.15 | 0.02 | 0.03 |
|  | LUNA-3D (min) | 46.3 | 35.6 | 82.9 | 107.5 | 5.4 | 5.8 | 0.74 | 1.2 |

#### ORBIS keeps a continuous tissue close to its true size and density

Which cells can interact, how densely a region is packed and how far apart two populations lie all depend on distances, and a continuous block can check them in three dimensions, because every measured cell has its true neighbors at their true distances (block B [8]; positions as produced, after a rigid alignment only). Our reconstruction keeps 92/99/93% of the true extent along the three axes; LUNA-3D keeps 71/79/77%, packing the same cells into 43% of the true volume (Figure 2a–c). Its cells therefore have 2.0–2.5 times as many neighbors as in the tissue at every radius from 10 to 200 *μ*m, against about 1.2 times for ours (median within 30 *μ*m: 29 cells, 12 in the tissue, 14 in ours), and the distances between cell-type centers shrink together (slope 0.68, mean distortion 0.42, pairs more than 10% too close/too far 86/4%, against 0.93, 0.22 and 43/23% for ours; Figure 2d–g). Block A repeats both patterns (crowding about 2 versus at most 1.5 times, distortion 0.35 versus 0.24). The contraction reaches the biology: LUNA-3D mixes cell types that live apart, presses the cortical layers together and leaves sections near the edges of the block empty (Supplementary Section S9). It is the signature of returning a single best guess per cell: a layout of best guesses behaves like an average and is smaller than the tissue by the uncertainty of each cell (Supplementary Section S7). LUNA-3D’s network does this: given the true layout with almost no noise, it predicts nearly the same layout as from pure noise, so its prediction is set almost entirely by expression and its sampler ends at about one guess per cell. Our network instead reads the layout it is given, and its prediction improves as that layout approaches the truth (Supplementary Section S8), so its sampler draws one configuration of the whole block, which keeps the spread of the tissue and, with it, nearly its real size and density.

**Figure 2:**
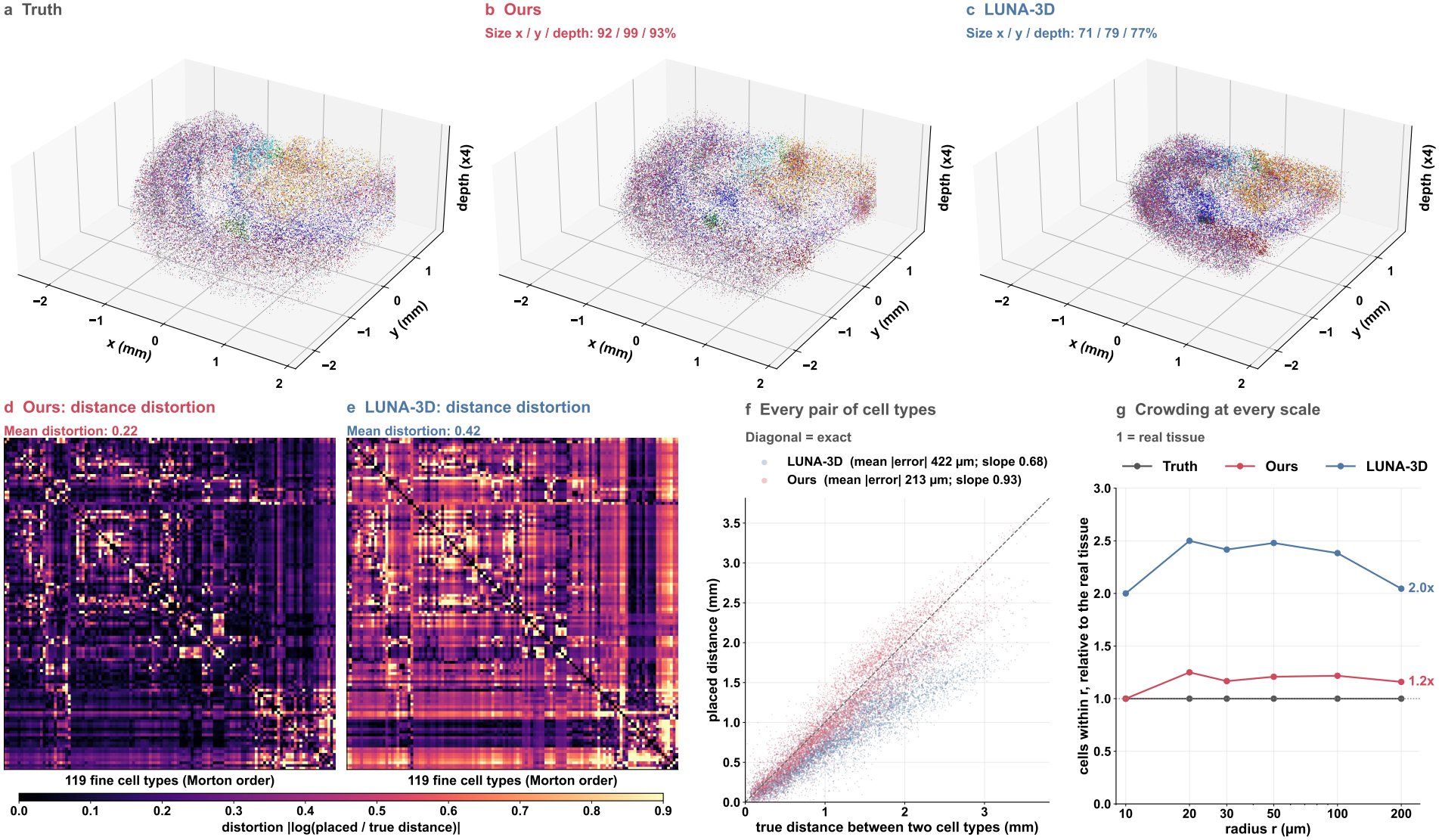
Size and density of a continuous tissue (Deep-STARmap block B, 198,675 cells). (a–c) Truth, ours and LUNA-3D at true scale, colored by cell class (depth stretched 4 ); titles: extent relative to the truth. (d, e) Distortion 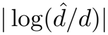 of the distances between the centers of 119 fine cell types, ours (d) and LUNA-3D (e), no rescaling; types in Morton order of their true centers. (f) Placed versus true distance per type pair. (g) Median number of cells within *r*, relative to the truth (1 = real density).

#### ORBIS preserves the positional order of cells within an organ

A whole-body reconstruction serves developmental biology only if cells keep their position *within* an organ, not merely inside the right organ. The digestive tract is a clear case: in the embryo, homeotic genes acting across germ layers assign the gut its regional identity along the anterior–posterior axis [60, 61], and the adult midgut is divided along its length into regions with distinct functions [62]. We therefore ask whether each gut cell of a held-out embryo is placed at its true anterior–posterior position (embryo c; 4,784 gut cells in six annotated parts from pharynx to hindgut; Figure 3), scored by the rank correlation between true and placed positions. To separate order from part identity, a reference places every cell at a random true position of its own gut part. Our reconstruction keeps the order of the gut (Spearman 0.92 against 0.78 for LUNA-3D) and exceeds the reference (0.81): it recovers where each cell lies within its part, not only which part it belongs to. Inside the largest part, the midgut chamber, the difference widens (0.57 versus 0.19), and both results repeat on the second held-out embryo (whole gut 0.87 versus 0.69, reference 0.63; midgut chamber 0.64 versus 0.21). The same holds beyond the gut: our model keeps the anterior–posterior order of cells inside their organ better than LUNA-3D in all 21 organs of embryo c and in 19 of 22 organs of embryo d (mean 0.52 versus 0.35 and 0.47 versus 0.36). This is consistent with sampling: a single best guess per cell pulls cells of similar expression toward the middle of their part (Supplementary Section S7), whereas a sample keeps the spread of the part and, with it, the order of its cells.

**Figure 3:**
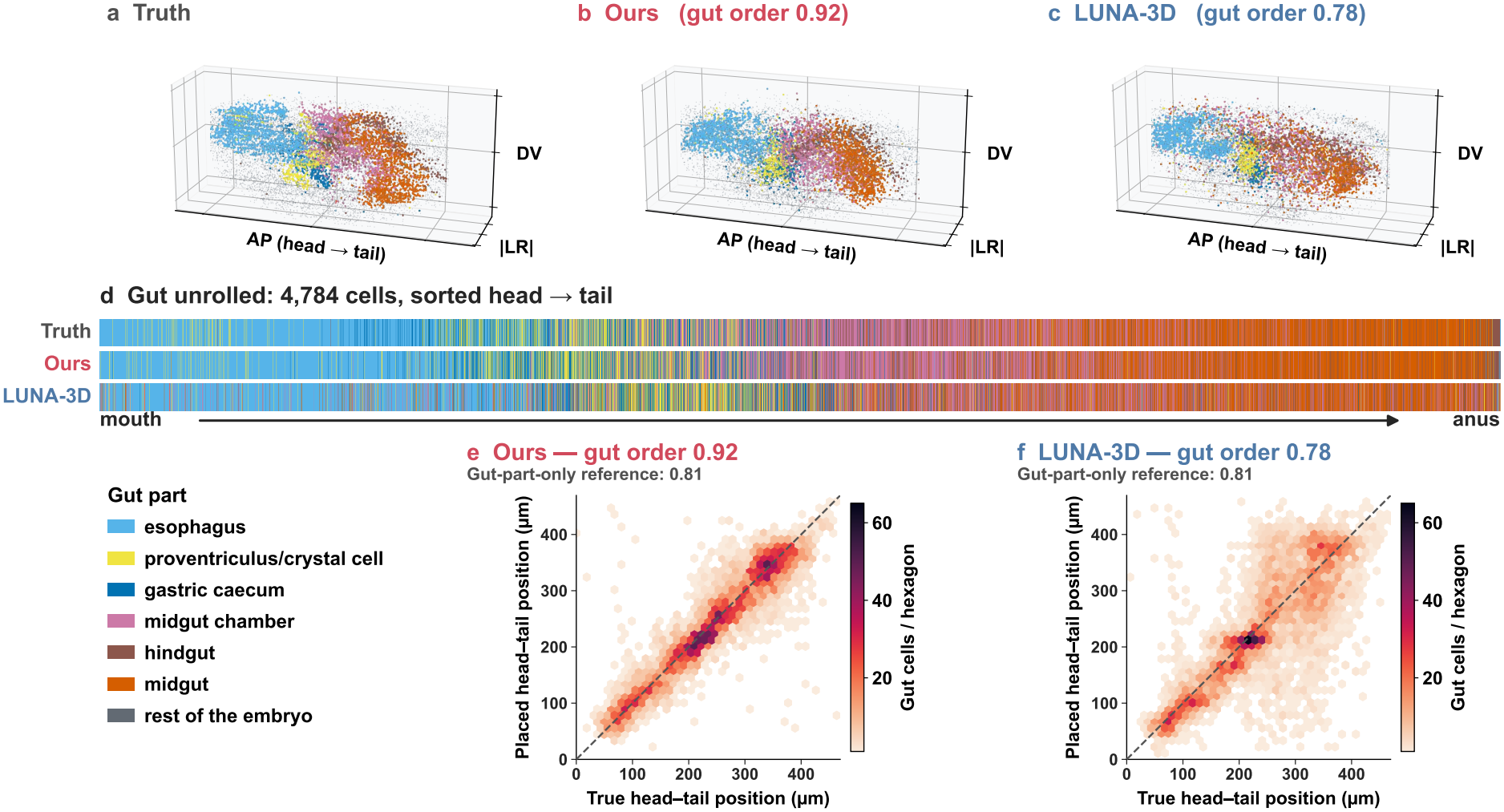
Positional order in the gut of held-out *Drosophila* embryo c (4,784 cells, six parts). (a–c) Truth, ours and LUNA-3D. (d) One line per gut cell, sorted head to tail by placed position; the truth shows the parts as ordered blocks. (e, f) True versus placed head-to-tail position, ours (e) and LUNA-3D (f).

#### ORBIS returns cells to their region of origin on a dissociated single-cell atlas

Every test so far withholds coordinates that were measured. The cells a reconstruction is built for have none: they are dissociated, profiled by another technology with another gene panel, and carry at most the region they were dissected from. We place cells of the Allen whole-brain single-cell atlas [16], five cohorts of 100,000 10x v3 cells drawn without regard to region, with our model and LUNA-3D, both trained on all five MERFISH brains and given only each cell’s expression mapped into the expression space of the spatial training data [63]; the dissection region, one of 23, is used only to score, after the same label-free rigid alignment of each layout to the atlas (Supplementary Section S12). Averaged over the 23 regions, our model returns 37.9% of a region’s cells to it, more than ten times random placement (3.6%), against 28.5% for LUNA-3D (95% interval of the difference over donors, 8.5–10.3 points), and it is ahead in 19 of the 23 regions (Figure 4). The advantage holds under the official Allen anatomy and in cohorts balanced across regions (8.5–9.9 points; Supplementary Figure S13). It does not come from recognizing cell types alone: placing each cell at a random atlas position of its own type reaches 27.4%, which LUNA-3D barely exceeds, whereas our model gains another 10.5 points from the differences in expression within a type, and shuffling its positions among cells of the same type removes this gain (25.2%). LUNA-3D also fails outright in parts of the cortex, returning almost none of the cells of the anterior cingulate and prefrontal cortex (0.1% and 1%), of which our model returns more than a quarter. Each cell type is also placed with the right geometry: the placed cloud of a subclass lies 402 *μ*m from its MERFISH counterpart, against 694 *μ*m for LUNA-3D, and ours is closer for 161 of 185 subclasses. This difference is consistent with how the two models are trained. ORBIS learns cell positions in a physical coordinate frame shared across reference brains, allowing it to capture regional differences in expression even within the same cell type. In contrast, LUNA-3D learns pairwise distances within sampled cell subsets, without a shared anatomical frame. This may limit its ability to transfer spatial organization to independently profiled cells from a different technology.

**Figure 4:**
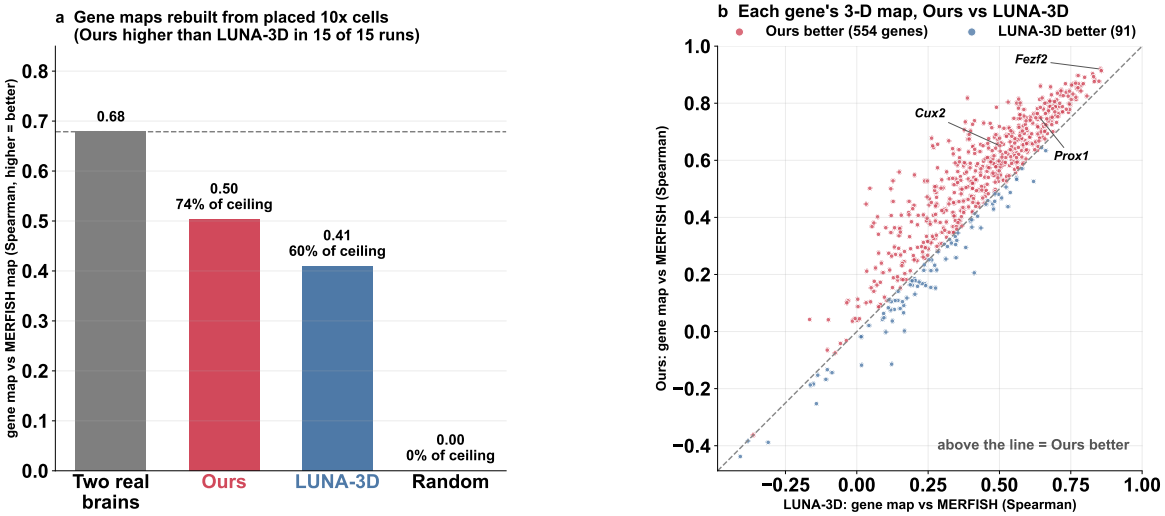
Placing dissociated Allen 10x v3 cells in a whole mouse brain (both models trained on five MERFISH brains, given expression only). (a–h) Side view: grey, MERFISH cells; orange, one dissection region; dots, the 10x cells from that region placed by ours (a–d) and LUNA-3D (e–h). Corner: hit rate, mean of 5 cohorts × 3 seeds (one shown). (i) Share of each region’s cells placed inside it (most of their 15 nearest atlas cells in the region, the nearest within 500 *μ*m); dashed, random. (j) Sliced Wasserstein-2 distance between placed and MERFISH cells per subclass (185); below the diagonal, ours is closer.

### ORBIS placements generalize to unseen genes

Each placed cell carries its full transcriptome, allowing reconstructed positions to recover spatial expression patterns beyond the input gene panel. We test this on 645 genes of the MERFISH panel [15] that lie outside the 477 input genes, so their maps can be right only if the right cells were placed in the right places for reasons other than these genes. The genes are chosen on the MER-FISH training brains alone: each is detected in at least 200 cells in at least two brains and has a non-constant map. For each gene we build a map of its mean expression per cell from the placed 10x cells and compare it by Spearman correlation with the map from the measured cells of the four MERFISH mice of that panel, on the volume both methods cover; the same score between two real brains (mouse 1 against mice 2–4) gives the ceiling of 0.68 (Supplementary Section S12). The map is the Gaussian-weighted sum of a gene’s expression over nearby cells divided by the Gaussian-weighted number of cells, so it reports the pattern of the gene rather than the density of cells, with *σ* = 200 *μ*m on a 200 *μ*m grid, the finest scale at which the serially sectioned reference brains give a continuous map. The Spearman correlation compares where a gene is high and where it is low, so it is unaffected by the different detection efficiencies of the two platforms, and a map value is used only where at least five placed cells contribute. The maps rebuilt from our placement reach 0.50 against 0.41 for LUNA-3D, 74% and 60% of the ceiling (Figure 5a), higher in all 15 runs (difference 0.094 ± 0.003) and for 554 of the 645 genes (Figure 5b). This includes markers [64, 16] of the upper cortical layers (*Cux2*, 0.65 against 0.50, ceiling 0.76), of the dentate gyrus (*Prox1*, 0.75 against 0.64, ceiling 0.83) and of the deep cortical layers (*Fezf2*, 0.92 against 0.85, ceiling 0.93). The ranking does not depend on how the volume a layout leaves empty is scored: scoring each layout on its own volume, or on the whole brain with empty cubes set to the brain-wide mean of the gene, widens the margin to +0.107 and +0.116, in all 15 runs and for 563 and 571 genes (Supplementary Table S11), so the shared-volume score is the conservative one. The same 100,000 cells cover 58% of the brain volume under our placement and 41% under LUNA-3D, consistent with the contraction of LUNA-3D in the continuous block (Figure 2). Shuffling our positions among cells of the same type lowers the score in all 15 runs and for 582 of the 645 genes: where a cell is placed within its type adds to the map.

**Figure 5:**
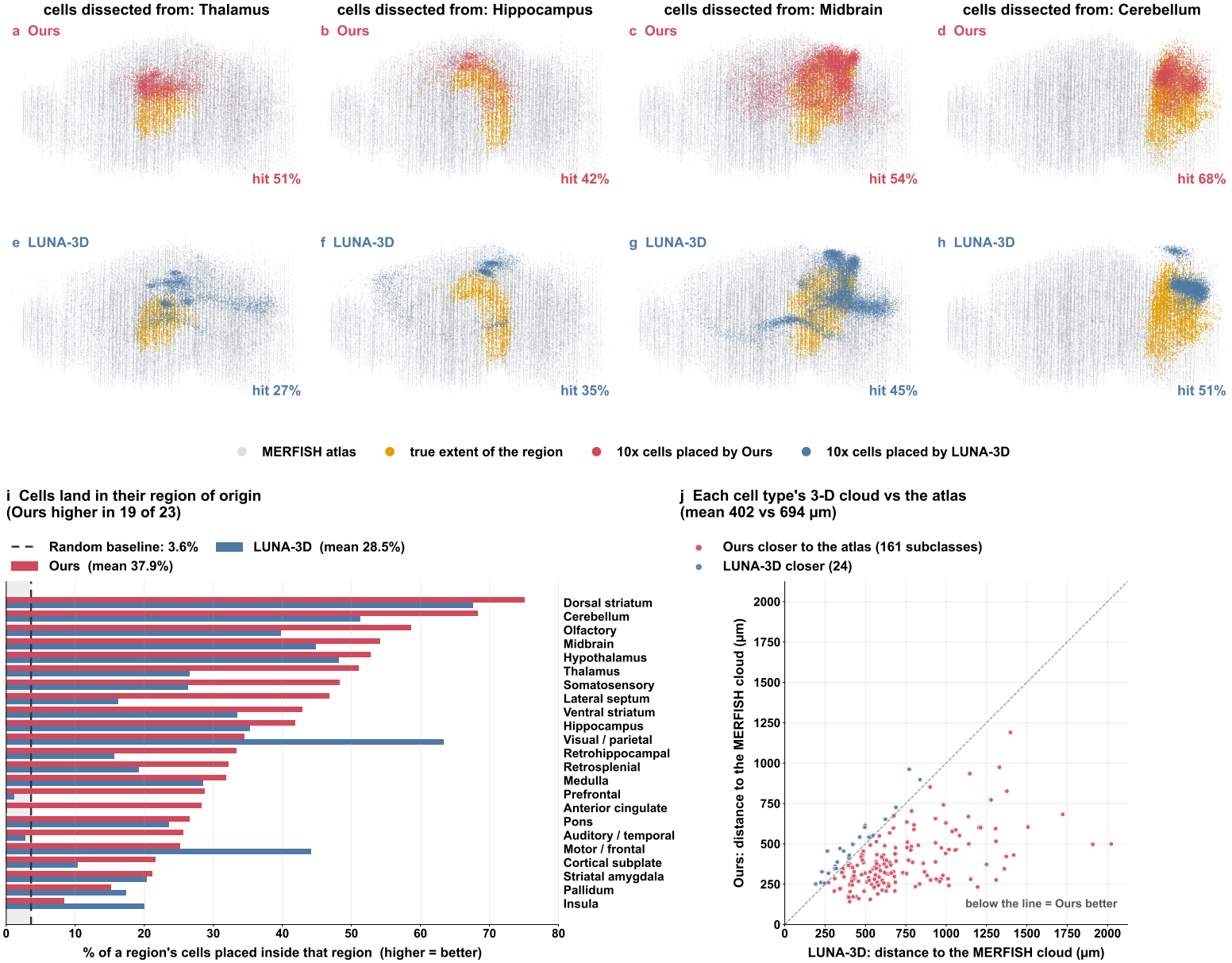
Maps of 645 unseen genes, rebuilt from placed 10x cells (15 runs). (a) Spearman correlation with the MERFISH reference map, mean over genes; percentages: share of the ceiling, same score between two real brains (dashed). (b) One dot per gene, ours against LUNA-3D; above the diagonal, ours is closer.

## 6 Concluding Remarks

ORBIS demonstrates that reconstructing a dissociated specimen in three dimensions is a problem of generating a coherent tissue, not of assigning positions to individual cells. By learning in a shared physical frame and updating a single specimen-wide state, it generates configurations of up to 3.7 million cells without stitching separately reconstructed regions. Across eight held-out specimens spanning mouse brains, tissue blocks and *Drosophila* embryos, ORBIS achieves lower cell-population shape error than all five baselines on every test, and on an independently profiled dissociated atlas it returns more cells to their region of origin than LUNA-3D. These results highlight the importance of preserving tissue size, geometry and neighborhood organization alongside per-cell accuracy. However, important limitations remain. Expression does not uniquely determine the position or the exact contacts of an individual cell, and serial-section references leave the detail between sections unresolved. Despite the abundance of measured cells, independent reference specimens remain few, and generalization to diseased or perturbed tissue has yet to be established. Larger and more diverse spatial atlases, especially volumetric whole-organ references, will be essential to improve generalization and assess uncertainty, and ORBIS provides a scalable foundation for using them.

## Supplementary Material

Generative 3D Spatial Reconstruction of Cellular Organization from Single-Cell Expression

### S1 Method details

This section gives the implementation details of Section 4, in the same order.

#### S1.1 Flow matching in a shared physical frame

##### The frame of each tissue

The reference specimens of a tissue are expressed in one frame with physical units. The brains are rotated to the axes of the Allen Common Coordinate Framework (medial–lateral, dorsal–ventral and anterior–posterior) and brought to one scale (Supplementary Section S2). Each Deep-STARmap block keeps its imaging frame, with two axes in the plane of the block and one through its depth, in micrometers. The embryos are posed on their body axes: the anterior–posterior axis is the first principal axis of the cell cloud, oriented by the expression centroids of anterior and posterior marker genes; the dorsal–ventral axis is the difference between the dorsal and ventral marker centroids, orthogonalized; and the left–right axis completes the frame. Each embryo is posed from its own expression and positions only.

##### Units

Positions are divided by one length *U* per tissue, shared by all its specimens and by all three axes, so the model sees the true size and the true proportions of every specimen. For the Deep-STARmap blocks and the embryos, *U* is set on the training specimens alone so that their root-mean-square coordinate over cells and axes is *d* = 0.163 (*U* = 5.64–5.66 mm for the blocks, 0.452 mm for the embryos); for the brains, *U* = 13.46 mm, which puts the complete training brains at 0.159–0.163. Every tissue therefore has the same spread against noise of unit variance, so a time *t* means the same ratio of signal to noise in every tissue, and one time distribution (Supplementary Section S1.4) and one set of step times (Supplementary Section S1.5) serve all tests. The same length converts the generated positions of a test specimen back to micrometers, followed for the brains by the per-axis slopes of Supplementary Section S2.

##### Path and source

The rows of **Y**_0_ are drawn independently from (**0, I**_3_), and both **Y**_0_ and **Y**_1_ are centered over the set of cells, so Eq. (2) carries no translation: a set keeps the orientation and the scale of its specimen, but not its location in space, which is arbitrary for a whole specimen. Noise and data are coupled independently, without reflow [49]. In Eq. (4) the sampler divides by 1 − *t* + *δ* with *δ* = 10^*−*3^, which keeps the slope finite as *t* ∥ 1; the network itself is trained on the endpoint of Eq. (3).

#### S1.2 Training views

##### Drawing a view

On the brains and embryos the kind of each view is whole-specimen, section or block with probabilities 0.5, 0.25 and 0.25; the Deep-STARmap blocks use whole-specimen views only. The reference specimen is chosen in proportion to its number of cells. A whole-specimen view holds *n* ∼ *U* [10^4^, 2 × 10^4^] cells drawn uniformly from the specimen. A section view holds every cell of one section, or 2 × 10^4^ of them drawn at random when the section holds more. A block view holds *n* cells drawn uniformly from *k* ∼ *U {*2, …, 64} consecutive sections, with *k* capped by the number of sections of the specimen. Every view is centered and keeps the orientation and scale of its specimen.

##### View label

The label takes five values: whole specimen, and section or block, each with its cutting plane (coronal or sagittal for the brains; the plane of the section stack for the embryos).

##### Count thinning

Before the transform log_2_(1 + count), the counts of every view are thinned binomially, each count kept with a probability *p* ∼ *U* [0.55, 1] drawn once per view. Thinning changes no position and needs no label. The model for the Allen brain, whose input is integrated expression components rather than counts (Supplementary Section S2), is trained without thinning.

##### Left–right fold

For the embryos, every view is folded across the midline plane of the body frame by replacing the left–right coordinate by its absolute value. The embryos are evaluated folded in the same way for every method (Supplementary Section S4).

#### S1.3 Set denoiser

##### Embeddings

*E*_gene_ and *E*_pos_ in Eq. (5) are two-layer perceptrons with SiLU activation into the model width of 256. The expression input is log_2_(1 + count) over the measured genes of each dataset (1,122 for the brain atlas, 1,017 for the tissue blocks and 6,936 for the embryos), or the 200 integrated components for the Allen brain (Supplementary Section S2). *E*_time_ in Eq. (6) is a 256-dimensional sinusoidal encoding followed by a two-layer perceptron, and *E*_view_ a linear map of the one-hot label, initialized to zero so that the label starts with no effect.

##### Transformer blocks

The network has eight blocks of width 256 with eight attention heads. Each block applies layer normalization without learned affine parameters, modulated by a shift, scale and gate computed from **z**, then multi-head self-attention over all tokens of the view, then a feed-forward network of width 1,024 with GELU activation, each with a residual connection. The projections that compute the gates are initialized to zero, so every block starts as the identity (adaLN-Zero [53]). Queries and keys are layer-normalized within each head before attention (QK normalization), which bounds the attention logits and keeps training stable on sets of tens of thousands of cells. Attention uses PyTorch’s scaled dot-product attention, and the blocks run in bfloat16. The model for the brain atlas has 10.2 million parameters.

##### Output

A final layer, modulated by **z** in the same way, maps each token to three coordinates, which are centered over the view.

#### S1.4 Coordinate supervision

##### Time distribution

We draw 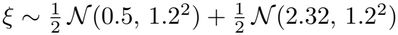 and set *t* = sigmoid(*ξ*), clipped to [10^*−*3^, 1 − 10^*−*3^]. The second component is EDM’s distribution of noise levels, ln *σ* ∼ *N* (− 1.2, 1.2^2^) at a data scale of 0.5 [57], carried over to our path at our data scale *d* = 0.163: on the path of Eq. (2) the noise relative to the data is (1 − *t*)*/*(*t d*), and equating it with EDM’s *σ/*0.5 gives logit *t* = ln(0.5*/d*) − ln *σ*, whose mean is ln(0.5*/*0.163) + 1.2 = 2.32. Each view receives one draw of *t* and one draw of the source **Y**_0_.

##### Optimization

We use AdamW with AMSGrad (*β*_1_ = 0.9, *β*_2_ = 0.999) at a learning rate of 5 × 10^*−*4^, constant and then decayed linearly to zero over the last 60 epochs (50 for the embryos), with gradients clipped to a norm of 0.05. Each optimizer step uses two views, and an epoch holds 246 views. The brain and block models are trained for 360 epochs and the embryo models for 300; training for 300 gives the same results. We keep an exponential moving average of the weights with decay 0.999 per step and sample with the averaged weights.

#### S1.5 Whole-specimen sampling

##### Start and step times

The rows of the initial state are drawn independently from *N* (**0, I**_3_) and centered. We take *K* = 50 steps at the times of EDM [57] (*σ*_max_ = 80, *σ*_min_ = 0.002, *σ*_data_ = 0.5, *ρ* = 7), carried over to our path by *t*_*k*_ = 1*/*(1 + *r*_*k*_ *d*), where 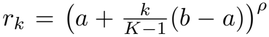 with *a* = (*σ*_max_*/σ*_data_)^1*/ρ*^ and *b* = (*σ*_min_*/σ*_data_)^1*/ρ*^, and *d* = 0.163 is the data scale (Supplementary Section S1.1); a final step goes to *t*_*K*_ = 1. The first time is *t*_0_ = 0.037, and the initial state is scaled by 1 − *t*_0_ so that it matches the noise of the path at *t*_0_. In Eq. (9) the sampler divides by 1 − *t*_*k*_ + *δ* with *δ* = 10^*−*3^. Steps are first-order Euler steps without guidance or stochastic correction.

##### Windows

At every step a uniform random permutation of the *N* cells is split into ⌈*N/*(2 × 10^4^) windows of nearly equal size, at most 2 10^4^ cells, so that every cell is denoised exactly once per step and a window has the size and spread of a whole-specimen training view. The partition is drawn afresh at every step. Every window is given the whole-specimen label.

##### Cost and output

A sample costs *K* ⌈*N/*(2 × 10^4^)⌉ network evaluations, for example 50 × 80 = 4,000 for a brain of 1.6 million cells. Each draw of the initial state and of the partitions gives a different sample. The final state is multiplied by the length *U* of the tissue (Supplementary Section S1.1) to return to micrometers.

#### S1.6 Reseating

##### Neighborhood reader

For each test, the reader is fitted on its training specimens only. Expression (genes selected by Moran’s I; the 200 integrated components for the Allen brain) is standardized, reduced to 30 principal components and modeled by a Gaussian mixture with 32 diagonal components; the posterior memberships **p**_*c*_ ∈ Δ^31^ are the soft expression groups. The targets are the mean memberships of each cell’s 32 and of its 128 nearest other cells (within its section for the sectioned brains, in three dimensions for the blocks and embryos). For each of the two scales, a ridge regression and a multilayer perceptron are fitted from expression to the target, with the regularization and early stopping set on held-out sections, tiles or embryos, and the one with the higher held-out *R*^2^ gives the prediction 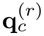 projected onto the probability simplex. The per-group scales 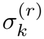 are the held-out errors of a reader fitted without the held-out rows. The test specimen contributes only its expression.

##### Energy

In full, Eq. (10) reads

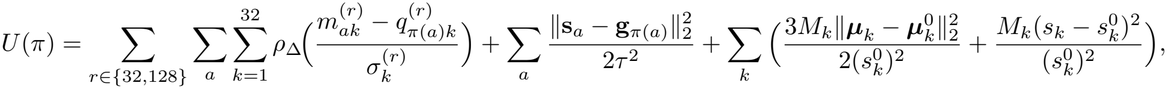

where 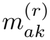 is the mean membership *p*_*π*(*b*)*k*_ over the *r* nearest other sites *b* of site *a, ρ*_Δ_ is the Huber loss with Δ = 1.345, and *M*_*k*_, ***μ***_*k*_ and *s*_*k*_ are the mass, membership-weighted centroid and spread of group *k* over the sites, with 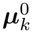 and 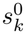 their values in the sample. The single-call estimate **g**_*c*_ is the output of one network call from noise with the same sampler, rigidly fitted onto the sample (rotation and translation, no scale, no truth), and *τ* ^2^ is the mean over cells of 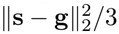.

##### Moves

We run 128 sweeps. In each sweep every site proposes one swap with a site chosen uniformly among its 30 nearest sites, and the swap is accepted with the Metropolis rule at unit temperature on the change of the total energy. A swap is refused if either cell would end farther than *τ* from its sampled site. Swaps whose neighborhoods do not overlap change the energy independently, so the sweeps run on a GPU as parallel exact swaps on a checkerboard of spatial blocks, with the energy recomputed from scratch after every sweep. Reseating changes only which cell occupies which site; the set of positions is unchanged.

### S2 Data and tests

Table S1 lists the eight tests. Each probes a distinct generalization regime: a held-out animal (mice 1 and 2, cut coronally, like one of their training brains); a held-out cutting orientation (mouse 3, cut sagittally and reconstructed by a model trained only on the two coronal mice 1 and 2); a held-out laboratory and platform (the Allen brain, reconstructed by a model trained on all four mice); a held-out tissue block (each Deep-STARmap block, reconstructed by a model trained on the other); and held-out embryos. A test specimen never enters training. Of a test specimen, every method receives only the expression of its cells: no positions, section identities, cell-type labels or tissue size. The measured positions are used only for scoring.

**Table S1:**
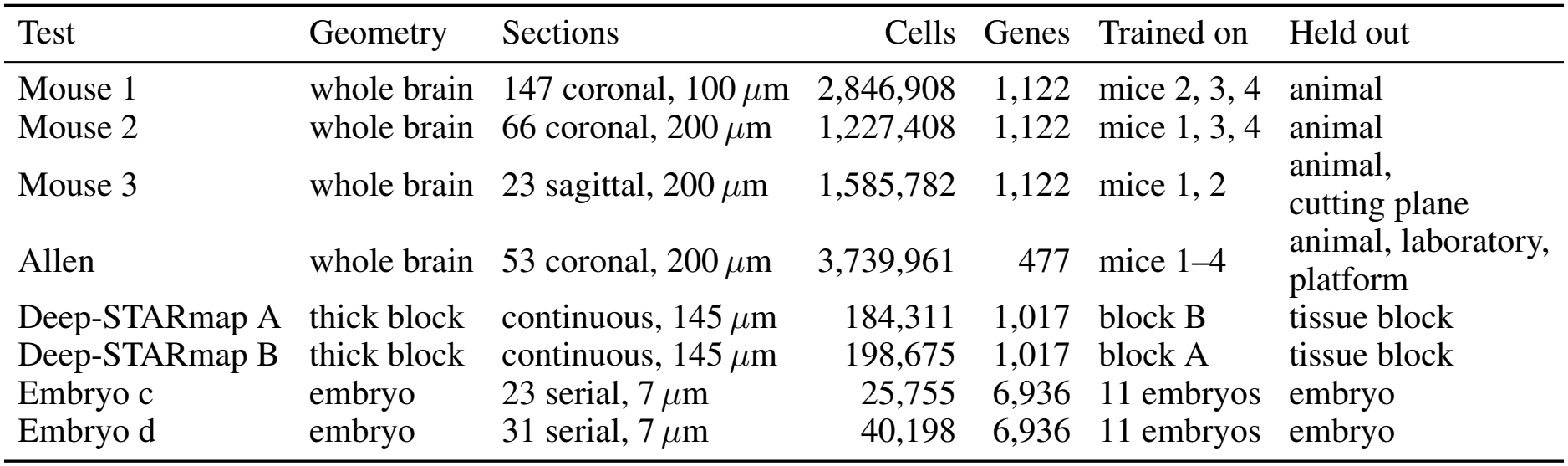
Test specimens and the training specimens of each test. Sections: number of serial sections, their plane and spacing (blocks: depth).

| Test | Geometry | Sections | Cells | Genes | Trained on | Held out |
|---|---|---|---|---|---|---|
| Mouse 1 | whole brain | 147 coronal, 100 *μ*m | 2,846,908 | 1,122 | mice 2, 3, 4 | animal |
| Mouse 2 | whole brain | 66 coronal, 200 *μ*m | 1,227,408 | 1,122 | mice 1, 3, 4 | animal |
| Mouse 3 | whole brain | 23 sagittal, 200 *μ*m | 1,585,782 | 1,122 | mice 1, 2 | animal, cutting plane |
| Allen | whole brain | 53 coronal, 200 *μ*m | 3,739,961 | 477 | mice 1–4 | animal, laboratory, platform |
| Deep-STARmap A | thick block | continuous, 145 *μ*m | 184,311 | 1,017 | block B | tissue block |
| Deep-STARmap B | thick block | continuous, 145 *μ*m | 198,675 | 1,017 | block A | tissue block |
| Embryo c | embryo | 23 serial, 7 *μ*m | 25,755 | 6,936 | 11 embryos | embryo |
| Embryo d | embryo | 31 serial, 7 *μ*m | 40,198 | 6,936 | 11 embryos | embryo |

#### Whole brains

Mice 1–4 are single hemispheres of adult C57BL/6J mice profiled with MERFISH by one laboratory [15]; mouse 4 has only three sagittal sections and is used for training only. The Allen brain (animal 638850) was profiled by another laboratory on the MERSCOPE platform and covers both hemispheres [16]. Each training brain’s published section-stack coordinates are rotated rigidly to its own registration to the Allen Common Coordinate Framework [65], so that the axes are medial–lateral, dorsal–ventral and anterior–posterior in every brain. The published coordinates mix two rulers (in-plane axes measured in tissue, the section axis estimated in atlas units), so each training brain is brought to the scale of one reference training brain by one linear factor per axis, the ratio of its least-squares slope of atlas on own coordinates to that of the reference brain; the test brain plays no part. A linear factor per axis keeps straight lines and the order of cells along every axis unchanged. Expression is log_2_(1 + count). For the Allen brain, which differs from the training mice in platform and gene panel, the 477 shared genes are integrated with Harmony [66] (200 principal components), and every method receives this input. Brains are scored in atlas-calibrated units: the measured positions of a test brain are multiplied, axis by axis, by its own slope, and the generated positions of every method by the slope of the reference training brain, fixed before any test cell is seen.

#### Thick blocks

The two Deep-STARmap blocks are coronal slabs of mouse brain of about 4.2 × 4.2 mm and 145 *μ*m [8]; each is reconstructed by a model trained on the other.

#### Embryos

The Flysta3D-v2 *Drosophila* embryos are reconstructed from serial 7-*μ*m sections [50]. The two test embryos (16.9 and 17.3 h) lie within the age range of the 11 training embryos (13.8–19.8 h). Positions are divided by one factor shared by all embryos and set on the training embryos.

### S3 Baselines

#### Choice of baselines

LUNA [45] represents generative reconstruction: it trains a diffusion model over the cells of a section, conditioned on their expression, and needs no spatial reference at inference. G2T [46] builds on the design of LUNA and recovers coordinates by multidimensional scaling, an eigendecomposition of an *N* × *N* matrix. This is extremely costly to scale to whole 3D specimens: in single precision the matrix alone needs about 6 TB at 1.2 million cells and 56 TB at 3.7 million, far beyond our largest GPU (96 GB), and in two dimensions one sampling step already takes about 80 seconds on a section of 33,000 cells. GEARS [44] has a related limit: it recovers 2D coordinates through an eigendecomposition on landmarks of distances stitched from patches, whereas a whole specimen calls for one shared state rather than stitched pieces. LUNA therefore represents this line (Section 2 of the main text). REMAP [38], CeLEry [36], Tangram [25] and novoSpaRc [31] represent reference-based reconstruction: REMAP and CeLEry learn a mapping from expression to coordinates on the reference, Tangram assigns each cell to reference cells, and novoSpaRc transports cells onto reference locations. Each draws its spatial reference from the same training specimens as our model, so no baseline sees more spatial data than our model.

#### Implementation

Every baseline was run from its official repository, with the changes and settings in Table S2. REMAP handles three-dimensional references as released; the other four were extended from two to three coordinates at the places where they fix the coordinate dimension, with no other change to the method. We also made memory-only changes where million-cell inputs did not fit, each verified to reproduce the released computation. We did not tune or correct the released methods, and every method was scored by the same code as our model. LUNA learns from graphs the size of a tissue section; because a whole specimen holds up to millions of cells, LUNA-3D is trained on random subsets of about 5,000 cells of each training specimen, the median section size of LUNA’s own training data. LUNA’s output is in the normalized units of its training graphs; we convert it with the scale of the training specimens, without using any true size, and align it with the same single rigid motion as every method.

### S4 Evaluation

#### Principles

The evaluation follows four rules. (i) Forgive only what expression cannot determine: the orientation, handedness and position of a specimen are removed by one rigid alignment, and a direction that the training data cannot teach is folded for every method alike. (ii) Do not penalize an exchange of positions between two cells of the same fine type, which expression does not distinguish in our annotation; the population metrics (shape, squeezing) are exactly invariant to such exchanges, while per-cell metrics (placement, per-cell error, niche NMI) are not and are reported separately. (iii) Score size and shape separately: shape is measured after matching each axis to the true size, and size is always reported next to it, so that a shrunken or collapsed reconstruction cannot pass as a correct one. (iv) Report physical units (*μ*m) wherever possible. Every metric is computed by one scoring program, with the same steps, for every method.

**Table S2:** Baselines: changes and training settings.

| Method | Changes and settings |
|---|---|
| LUNA-3D | Coordinate dimension 2 → 3 at every place where LUNA fixes it (11 lines; model otherwise unchanged); training graphs = random subsets of about 5,000 cells of each training specimen (a whole specimen far exceeds LUNA's section-sized graphs; 5,000 = the median section size of LUNA's own training data); memory-mapped data loading; global batch 8; 300 epochs (400 on the blocks). Output converted from the normalized units of its training graphs with the scale of the training specimens (no true sizes used); one rigid fit per specimen. |
| REMAP | Memory-only changes (chunked decoding; same computation). Official settings otherwise; all training cells, except the mouse 2 run (400,000-cell reference subsample, Harmony on). |
| CeLEry-3D | Three outputs instead of two (last layer, location loader, prediction columns); default hyperparam-eters. At least the number of updates of the authors' tutorial: 15–16 epochs on the blocks, 11 on the embryos, 1 on the brains. |
| Tangram-3D | Unmodified; cell mode, uniform density prior; each cell placed at the three-dimensional position of its most probable reference cell. Reference: 55,000 (blocks) and 48,000 (embryos) training cells, 1,000 epochs; brains: 15,000 cells, 500 epochs. |
| novoSpaRc-3D | Unmodified; atlas-assisted reconstruction as in the authors' *Drosophila* tutorial; position = transport-plan-weighted mean of the target locations. Targets: 10,000 training cells (blocks, embryos) and 5,000 (brains). |

#### Notation

A specimen has *N* cells with true positions **y**_*i*_ ∈ ℝ^3^ and reconstructed positions **ŷ**_*i*_, *i* = 1, …, *N* . Each cell has a fine cell type *c*(*i*) and a cell class *k*(*i*); the types partition the cells into sets *C*_*c*_ with *n*_*c*_ = |*C*_*c*_|. Fine types are the 1,188–1,201 supertypes of the atlas taxonomy on the brains, 137 annotated types on the blocks and 23 on the embryos; classes are the coarsest annotation level (34 on the brains). Types and classes are used only for scoring and never enter any model. *σ*_*a*_(·) is the standard deviation of coordinate *a* ∈ {1, 2, 3} over the cells of a layout.

#### Alignment

With type centroids 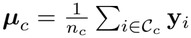 and 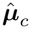 defined likewise on the reconstruction, we solve

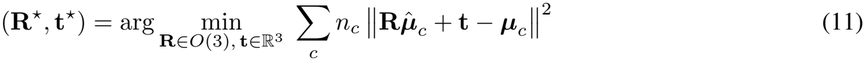

in closed form (Kabsch), with reflections allowed (det **R** = ±1) and no scaling, and replace **ŷ**_*i*_ by **R**^⋆^**ŷ**_*i*_ + **t**^⋆^. Centroids do not change when cells of one type exchange positions, so the alignment respects rule (ii), and it costs *O*(*N* ). In the embryos, the side of a cell in the bilateral body cannot be read from its expression, so the left–right coordinate of the truth and of every reconstruction is replaced by its distance from the layout’s own midline (the median of that coordinate), and the rigid motion minimizes the same weighted error between these folded type centroids, with each axis of the reconstruction matched to the true spread during the fit. Every training mouse is a single hemisphere, so no model can learn which hemisphere a cell of the Allen brain, which covers both, belongs to; the Allen brain is folded at its midline.

#### Size error

After alignment,

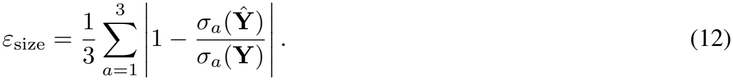

A reconstruction that is a uniformly scaled copy of the truth by a factor *s* has *ε*_size_ = |1 − *s*|.

#### Size matching

For the shape metrics, each axis of the aligned reconstruction is rescaled about its mean 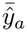 to the true spread,

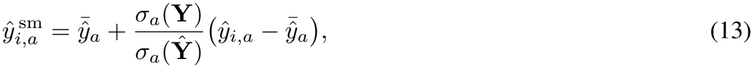

and the result is aligned again as above (on the embryos, whose fit already matches the spreads, it is not aligned again). Size matching removes any per-axis scale error, which size error reports, so the shape metrics measure only what remains.

#### Shape error (*W*_2_)

Let ***θ***_1_, …, ***θ***_64_ be fixed unit directions (isotropic Gaussian draws with a fixed seed, normalized), identical for every method. For a fine type *c* with *n*_*c*_ ≥ 20 and direction ***θ***_*m*_, let 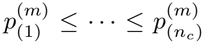 be the sorted projections 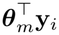 of its true cells and 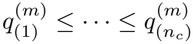 those of its size-matched reconstructed cells. Then

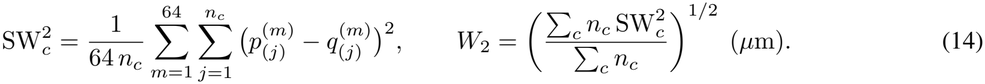

Each one-dimensional term is the exact Wasserstein-2 distance between the projected populations, so *W*_2_ compares where each population lies and how it is spread, and it is unchanged by any permutation of cells within a type (rule ii). It grows with misplacement and with blur of a population, and, since it is computed after size matching, it does not reward or penalize overall scale. Sliced distances are bounded above by the exact Wasserstein distance; *W*_2_ is used only to compare methods under identical settings.

#### F-score

For a fine type *c* with *n*_*c*_ ≥ 20 and a distance *τ*, the precision *P*_*c*_ is the share of its size-matched reconstructed cells that lie within *τ* of a true cell of type *c*, and the recall *R*_*c*_ the share of its true cells that lie within *τ* of a reconstructed cell of type *c*. With *F*_*c*_ = 2*P*_*c*_*R*_*c*_*/*(*P*_*c*_ + *R*_*c*_),

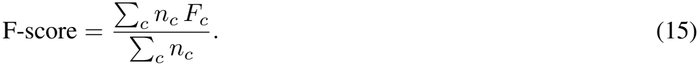

We set *τ* = 8*ℓ*, with *ℓ* the median nearest-neighbor distance between the true cells of the specimen (11.6–14.4 *μ*m on the brains, 14.1–14.2 *μ*m on the blocks, 4.5–4.6 *μ*m on the embryos), so that *τ* spans the same number of cell spacings in every tissue. Precision falls when a population spills into the wrong place and recall when part of it is missing; like *W*_2_, the score is unchanged by exchanges of cells within a type.

#### Type-centre error

For each fine type with *n*_*c*_ ≥ 200, the distance between the centroid of its size-matched reconstructed cells and its true centroid, averaged with weights *n*_*c*_ (*μ*m). It reports where each population lies, independently of its shape and spread.

#### Placement and per-cell error

With *d*_*i*_ = ∥**ŷ**_*i*_ − **y**_*i*_∥ after alignment (without size matching), the per-cell error is *e* = median_*i*_ *d*_*i*_ (*μ*m). As a reference, let *π*_*r*_ (*r* = 1, …, 10) be random permutations of the cells within each fine type, and 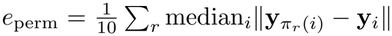 the error of placing every cell at the true position of a random cell of its own type. Then

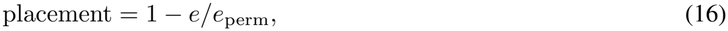

which is 1 for an exact reconstruction, 0 for one no better than a random position of the right type, and negative for worse. Expressing the error relative to *e*_perm_ makes it comparable across specimens of different size.

#### Niche NMI

For cell *i* of a layout, let *N*_100_(*i*) be its 100 nearest other cells and **f**_*i*_ their composition over the *K* cell classes with at least 20 cells, *f*_*ik*_ = | {*j* ∈ *N*_100_(*i*) : *k*(*j*) = *k*}|*/*100. *k*-means with 15 clusters (10 initializations, fixed seed) is fitted once on the compositions of the truth, giving centroids and true niche labels *z*_*i*_. For a reconstruction, the compositions 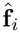 are computed from its own neighbors and every cell is assigned to the nearest true centroid, 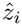. The score is the normalized mutual information

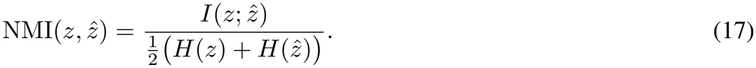

Fitting the niches once on the truth gives the truth a score of exactly 1; clustering truth and reconstruction separately would score the truth against itself below 1 because of *k*-means instability.

#### Type contacts

For a layout, let *M*_*ab*_ be the share of the 15 nearest other cells of the cells of type *a* that are of type *b*, at the contact level of each dataset (the subclasses of the atlas taxonomy on the brains, the 19 main types on the blocks and the 23 types on the embryos; types with at least 20 cells). The score is the Pearson correlation between the off-diagonal entries of *M* for the truth and for the reconstruction. It reports which cell types border which, the arrangement of layers and boundaries.

#### Moran’s I agreement

We select the 50 genes with the highest Moran’s I in the truth (on eight evenly spaced sections for serially sectioned specimens, on 100,000 random cells otherwise) and compute each gene’s Moran’s I on the row-normalized graph of the 15 nearest other cells, once with the true and once with the reconstructed positions. The score is the Pearson correlation of the two values across the genes. It reports whether genes keep their spatial patterns.

#### Squeezed cell types

For each fine type with *n*_*c*_ ≥ 200, the spread ratio

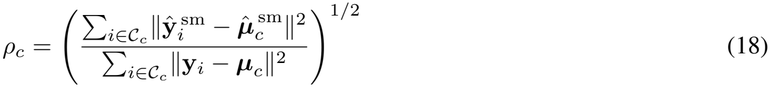

compares the spread of the type about its centroid after size matching with its true spread; a type is squeezed when *ρ*_*c*_ *<* 0.8, and we report the share of squeezed types. Because it is computed after size matching, it detects populations narrowed relative to the rest of the tissue, the signature of a reconstruction that averages over plausible positions.

#### 2-D metrics

Two-dimensional reconstructions have been evaluated by the per-cell distance correlation (SRC), contact precision and the root-sum-square deviation (RSSD) [45, 46]; we compute SRC and RSSD with the formulas of LUNA’s released code. SRC: on a class-stratified subsample of 10,000 cells, each cell’s distances to all subsample cells in the truth and in the reconstruction are compared by Spearman correlation, and the median over cells is reported. RSSD: for each cell class with at least 20 cells, the best proper rotation maps the class’s reconstructed positions onto its true positions (each layout centered at its own mean), and the root of the summed squared residuals is recorded; Sum RSSD adds these over classes (mm). Both are reported for our model and LUNA-3D on every test in Table S7.

#### Control layouts

Table S3 scores the true layout of mouse 2 after controlled corruptions. Three observations motivate our choice of headline metrics. First, a reconstruction shrunk uniformly to half size keeps a perfect SRC and niche NMI (1.000): scale-free scores suit a two-dimensional section, whose size and pose carry no meaning, but not a specimen reconstructed in physical units; size error detects the shrinkage. Second, exchanging cells within a fine type, which changes nothing that expression can determine, costs more under SRC (0.338) and RSSD (14,342) than collapsing every cell onto its type centroid (0.389 and 10,142), a reconstruction with no internal structure; *W*_2_ is exactly 0 for the exchange and 1,763 *μ*m for the collapse. Per-cell scores, including placement, share this preference, because the position that minimizes the expected error of a cell is the average of its plausible positions. Our reconstruction has an RSSD close to that of the collapsed layout (10,146), although its shape error is 200 rather than 1,763 *μ*m. Third, a single call of our network, which approximates this average, exceeds LUNA-3D on SRC and RSSD on all eight tests (Table S7). Size error and squeezing are one-sided guards: a method that copies the positions of reference cells keeps both small without placing cells correctly, so they are always read together with *W*_2_.

**Table S3:** Control layouts: the true layout of mouse 2 after controlled corruptions, scored by the same program as every method, with our reconstruction and LUNA-3D for reference. Exchanging cells within a fine type changes nothing that expression can determine; shrinking changes only size.

| **Layout** | **Size and shape** |  | **Per cell** | **Neighb.** | **2-D metrics** |  |
|---|---|---|---|---|---|---|
|  | Size err.↓ | *W*<sub>2</sub> (*µ*m)↓ | Placement↑ | Niche NMI↑ | SRC↑ | RSSD↓ |
| True layout | .000 | 0 | 1.000 | 1.000 | 1.000 | 0 |
| Shrunk uniformly to half size | .500 | 0 | .409 | 1.000 | 1.000 | 9,065 |
| One axis shrunk to 0.75 | .082 | 11 | .914 | .885 | .997 | 1,286 |
| Gaussian noise, 200 *µ*m per axis | .007 | 38 | .893 | .483 | .992 | 1,665 |
| Cells exchanged within each fine type | .000 | 0 | .001 | .291 | .338 | 14,342 |
| Every cell at its type centroid | .382 | 1,763 | .220 | .379 | .389 | 10,142 |
| LUNA-3D | .079 | 262 | .550 | .438 | .740 | 8,804 |
| **Ours** | .020 | 200 | .453 | .464 | .683 | 10,146 |

#### Protocol

Each test is reconstructed by three independent trainings of our model with different random seeds. Each training is sampled four times with different sampler seeds, and its output is the 50-step sample after neighborhood-consistent reseating. A training’s score is the mean over its four samples; we report the mean over trainings and the standard deviation across trainings. For the averages over a tissue geometry in the main table, each training’s scores are first averaged over the tests of the geometry and the mean and standard deviation are then taken across trainings. A difference from a baseline is called beyond noise only when it exceeds twice that standard deviation. Each baseline is run once and scored on its single output.

### S5 Full results

Tables S4–S6 give the per-test results behind the main-table averages, every method as the mean and standard deviation of three runs with different seeds. Every training of our model ran to its last epoch without loss spikes or rollbacks; one (Allen, seed 1) was interrupted by a hardware timeout and resumed exactly from its last checkpoint. In every one of its three trainings, LUNA-3D reconstructs mice 1 and 2 and both blocks 15–66% too small along at least one axis. Table S7 compares the one-call best guess with the sample.

**Table S4:** Per-test results: whole brains (Table 1 averages these rows). Every method: mean _±SD_ over 3 runs with different seeds; ours = the 50-step sample after reseating, each run the mean of four sampler draws. Columns as in Table 1; definitions in Section S4. **Bold**: best, <u>underline</u>: 2nd, *italic*: 3rd, within a test.

| **Test** | **Method** | **Specimen and cell types** |  |  |  | **Cells and neighborhoods** |  |  |  |
|---|---|---|---|---|---|---|---|---|---|
|  |  | Size error↓ | Shape *W*<sub>2</sub> (*μ*m)↓ | F-score ↑ | Type centre (*μ*m)↓ | Place-ment↑ | Niche NMI↑ | Type contacts↑ | Moran's *I* ↑ |
| **Mouse 3** | LUNA-3D | **.035**<sub>±.019</sub> | <u>385</u><sub>±186</sub> | <u>.461</u><sub>±.057</sub> | <u>413</u><sub>±240</sub> | **.471**<sub>±.094</sub> | <u>.450</u><sub>±.028</sub> | <u>.908</u><sub>±.016</sub> | <u>.784</u><sub>±.110</sub> |
|  | REMAP | .226<sub>±.003</sub> | 907<sub>±6</sub> | .335<sub>±.002</sub> | 930<sub>±7</sub> | .320<sub>±.009</sub> | *.434*<sub>±.004</sub> | *.844*<sub>±.005</sub> | *.598*<sub>±.066</sub> |
|  | CeLEry-3D | .217<sub>±.008</sub> | *641*<sub>±4</sub> | *.359*<sub>±.008</sub> | *608*<sub>±38</sub> | *.388*<sub>±.008</sub> | .399<sub>±.004</sub> | .805<sub>±.006</sub> | .568<sub>±.026</sub> |
|  | Tangram-3D | <u>.045</u><sub>±.003</sub> | 1,073<sub>±3</sub> | .116<sub>±.002</sub> | 952<sub>±11</sub> | −.153<sub>±.005</sub> | .217<sub>±.002</sub> | .465<sub>±.010</sub> | .053<sub>±.010</sub> |
|  | novoSpaRc-3D | .283<sub>±.003</sub> | 827<sub>±19</sub> | .347<sub>±.002</sub> | 943<sub>±35</sub> | .175<sub>±.007</sub> | .286<sub>±.001</sub> | .772<sub>±.013</sub> | .127<sub>±.026</sub> |
|  | **Ours** | *.054*<sub>±.004</sub> | **212**<sub>±6</sub> | **.511**<sub>±.004</sub> | **198**<sub>±18</sub> | <u>.457</u><sub>±.009</sub> | **.515**<sub>±.006</sub> | **.955**<sub>±.001</sub> | **.909**<sub>±.007</sub> |
| **Mouse 2** | LUNA-3D | *.181*<sub>±.141</sub> | <u>409</u><sub>±207</sub> | <u>.488</u><sub>±.101</sub> | <u>364</u><sub>±178</sub> | <u>.432</u><sub>±.177</sub> | <u>.398</u><sub>±.059</sub> | <u>.905</u><sub>±.030</sub> | <u>.701</u><sub>±.159</sub> |
|  | REMAP | .378<sub>±.053</sub> | 1,077<sub>±18</sub> | .355<sub>±.011</sub> | 1,092<sub>±13</sub> | .192<sub>±.004</sub> | .309<sub>±.015</sub> | *.833*<sub>±.006</sub> | .435<sub>±.042</sub> |
|  | CeLEry-3D | .274<sub>±.009</sub> | *699*<sub>±12</sub> | .388<sub>±.009</sub> | *624*<sub>±39</sub> | *.386*<sub>±.002</sub> | *.362*<sub>±.004</sub> | .808<sub>±.012</sub> | *.540*<sub>±.040</sub> |
|  | Tangram-3D | **.019**<sub>±.0003</sub> | 840<sub>±10</sub> | .191<sub>±.004</sub> | 685<sub>±12</sub> | −.049<sub>±.002</sub> | .210<sub>±.001</sub> | .650<sub>±.003</sub> | .503<sub>±.004</sub> |
|  | novoSpaRc-3D | .291<sub>±.006</sub> | 769<sub>±9</sub> | *.425*<sub>±.002</sub> | 625<sub>±19</sub> | .250<sub>±.004</sub> | .303<sub>±.006</sub> | .832<sub>±.005</sub> | .420<sub>±.013</sub> |
|  | **Ours** | <u>.020</u><sub>±.010</sub> | **200**<sub>±4</sub> | **.586**<sub>±.002</sub> | **199**<sub>±10</sub> | **.453**<sub>±.005</sub> | **.464**<sub>±.005</sub> | **.957**<sub>±.002</sub> | **.884**<sub>±.009</sub> |
| **Mouse 1** | LUNA-3D | *.108*<sub>±.022</sub> | <u>523</u><sub>±439</sub> | <u>.538</u><sub>±.241</sub> | <u>472</u><sub>±416</sub> | *.385*<sub>±.270</sub> | *.369*<sub>±.087</sub> | <u>.922</u><sub>±.030</sub> | <u>.802</u><sub>±.085</sub> |
|  | REMAP | .353<sub>±.008</sub> | 801<sub>±46</sub> | .467<sub>±.016</sub> | 763<sub>±44</sub> | .367<sub>±.016</sub> | <u>.397</u><sub>±.004</sub> | *.878*<sub>±.005</sub> | .613<sub>±.022</sub> |
|  | CeLEry-3D | .280<sub>±.012</sub> | *667*<sub>±31</sub> | .486<sub>±.014</sub> | 621<sub>±23</sub> | <u>.413</u><sub>±.004</sub> | .352<sub>±.009</sub> | .809<sub>±.017</sub> | *.666*<sub>±.035</sub> |
|  | Tangram-3D | **.028**<sub>±.002</sub> | 764<sub>±7</sub> | .224<sub>±.001</sub> | *584*<sub>±8</sub> | −.000<sub>±.001</sub> | .225<sub>±.001</sub> | .684<sub>±.011</sub> | .492<sub>±.012</sub> |
|  | novoSpaRc-3D | .323<sub>±.004</sub> | 878<sub>±9</sub> | *.514*<sub>±.003</sub> | 716<sub>±12</sub> | .234<sub>±.002</sub> | .267<sub>±.003</sub> | .853<sub>±.004</sub> | .543<sub>±.005</sub> |
|  | **Ours** | <u>.045</u><sub>±.007</sub> | **169**<sub>±5</sub> | **.720**<sub>±.009</sub> | **134**<sub>±7</sub> | **.494**<sub>±.011</sub> | **.457**<sub>±.008</sub> | **.962**<sub>±.002</sub> | **.901**<sub>±.011</sub> |
| **Allen** | LUNA-3D | *.095*<sub>±.057</sub> | <u>407</u><sub>±95</sub> | <u>.497</u><sub>±.039</sub> | <u>412</u><sub>±112</sub> | **.511**<sub>±.037</sub> | **.500**<sub>±.011</sub> | <u>.916</u><sub>±.009</sub> | *.826*<sub>±.010</sub> |
|  | REMAP | .198<sub>±.022</sub> | 727<sub>±16</sub> | .392<sub>±.007</sub> | 767<sub>±7</sub> | *.369*<sub>±.008</sub> | <u>.486</u><sub>±.004</sub> | *.873*<sub>±.005</sub> | <u>.850</u><sub>±.013</sub> |
|  | CeLEry-3D | .252<sub>±.015</sub> | *685*<sub>±18</sub> | .357<sub>±.008</sub> | *658*<sub>±19</sub> | .354<sub>±.007</sub> | .425<sub>±.012</sub> | .841<sub>±.009</sub> | .767<sub>±.015</sub> |
|  | Tangram-3D | <u>.059</u><sub>±.001</sub> | 855<sub>±3</sub> | .133<sub>±.002</sub> | 760<sub>±11</sub> | −.168<sub>±.005</sub> | .267<sub>±.0003</sub> | .523<sub>±.014</sub> | .473<sub>±.005</sub> |
|  | novoSpaRc-3D | .231<sub>±.003</sub> | 791<sub>±5</sub> | *.404*<sub>±.0005</sub> | 725<sub>±15</sub> | .138<sub>±.004</sub> | .265<sub>±.001</sub> | .808<sub>±.002</sub> | .635<sub>±.021</sub> |
|  | **Ours** | **.043**<sub>±.007</sub> | **212**<sub>±6</sub> | **.604**<sub>±.006</sub> | **199**<sub>±7</sub> | <u>.463</u><sub>±.004</sub> | *.479*<sub>±.003</sub> | **.941**<sub>±.0005</sub> | **.874**<sub>±.001</sub> |

**Table S5:**
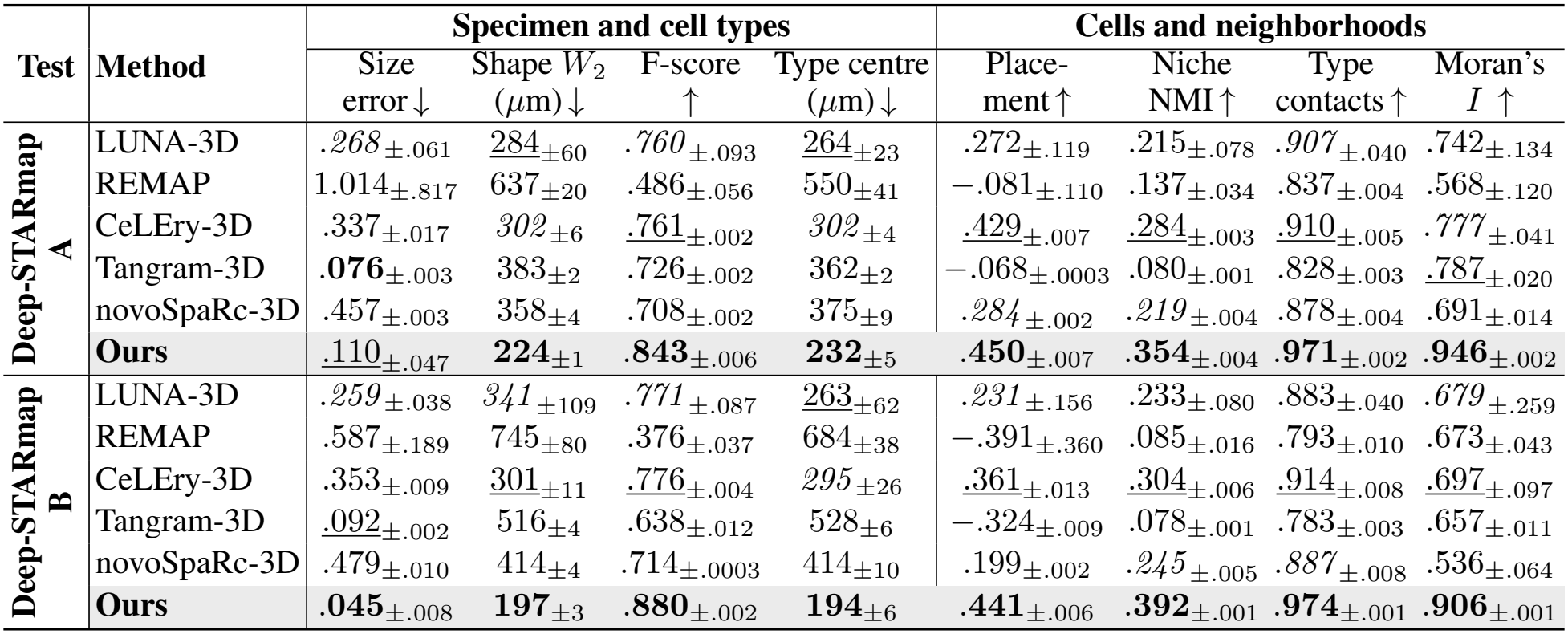
Per-test results: thick blocks (columns and marks as in Table S4).

|  |  | Specimen and cell types |  |  |  | Cells and neighborhoods |  |  |  |
|---|---|---|---|---|---|---|---|---|---|
| **Test** | **Method** | Size error↓ | Shape *W*<sub>2</sub> (µm)↓ | F-score ↑ | Type centre (µm)↓ | Place-ment↑ | Niche NMI↑ | Type contacts↑ | Moran's *I* ↑ |
| **Deep-STARmap A** | LUNA-3D | *.268*<sub>±.061</sub> | 284<sub>±60</sub> | *.760*<sub>±.093</sub> | 264<sub>±23</sub> | .272<sub>±.119</sub> | .215<sub>±.078</sub> | *.907*<sub>±.040</sub> | .742<sub>±.134</sub> |
|  | REMAP | 1.014<sub>±.817</sub> | 637<sub>±20</sub> | .486<sub>±.056</sub> | 550<sub>±41</sub> | −.081<sub>±.110</sub> | .137<sub>±.034</sub> | .837<sub>±.004</sub> | .568<sub>±.120</sub> |
|  | CeLEry-3D | .337<sub>±.017</sub> | *302*<sub>±6</sub> | .761<sub>±.002</sub> | *302*<sub>±4</sub> | .429<sub>±.007</sub> | .284<sub>±.003</sub> | .910<sub>±.005</sub> | *.777*<sub>±.041</sub> |
|  | Tangram-3D | **.076**<sub>±.003</sub> | 383<sub>±2</sub> | .726<sub>±.002</sub> | 362<sub>±2</sub> | −.068<sub>±.0003</sub> | .080<sub>±.001</sub> | .828<sub>±.003</sub> | .787<sub>±.020</sub> |
|  | novoSpaRc-3D | .457<sub>±.003</sub> | 358<sub>±4</sub> | .708<sub>±.002</sub> | 375<sub>±9</sub> | *.284*<sub>±.002</sub> | *.219*<sub>±.004</sub> | .878<sub>±.004</sub> | .691<sub>±.014</sub> |
|  | **Ours** | .110<sub>±.047</sub> | **224**<sub>±1</sub> | **.843**<sub>±.006</sub> | **232**<sub>±5</sub> | **.450**<sub>±.007</sub> | **.354**<sub>±.004</sub> | **.971**<sub>±.002</sub> | **.946**<sub>±.002</sub> |
| **Deep-STARmap B** | LUNA-3D | *.259*<sub>±.038</sub> | *341*<sub>±109</sub> | *.771*<sub>±.087</sub> | 263<sub>±62</sub> | *.231*<sub>±.156</sub> | .233<sub>±.080</sub> | .883<sub>±.040</sub> | *.679*<sub>±.259</sub> |
|  | REMAP | .587<sub>±.189</sub> | 745<sub>±80</sub> | .376<sub>±.037</sub> | 684<sub>±38</sub> | −.391<sub>±.360</sub> | .085<sub>±.016</sub> | .793<sub>±.010</sub> | .673<sub>±.043</sub> |
|  | CeLEry-3D | .353<sub>±.009</sub> | 301<sub>±11</sub> | .776<sub>±.004</sub> | *295*<sub>±26</sub> | .361<sub>±.013</sub> | .304<sub>±.006</sub> | .914<sub>±.008</sub> | .697<sub>±.097</sub> |
|  | Tangram-3D | .092<sub>±.002</sub> | 516<sub>±4</sub> | .638<sub>±.012</sub> | 528<sub>±6</sub> | −.324<sub>±.009</sub> | .078<sub>±.001</sub> | .783<sub>±.003</sub> | .657<sub>±.011</sub> |
|  | novoSpaRc-3D | .479<sub>±.010</sub> | 414<sub>±4</sub> | .714<sub>±.0003</sub> | 414<sub>±10</sub> | .199<sub>±.002</sub> | *.245*<sub>±.005</sub> | *.887*<sub>±.008</sub> | .536<sub>±.064</sub> |
|  | **Ours** | **.045**<sub>±.008</sub> | **197**<sub>±3</sub> | **.880**<sub>±.002</sub> | **194**<sub>±6</sub> | **.441**<sub>±.006</sub> | **.392**<sub>±.001</sub> | **.974**<sub>±.001</sub> | **.906**<sub>±.001</sub> |

**Table S6:**
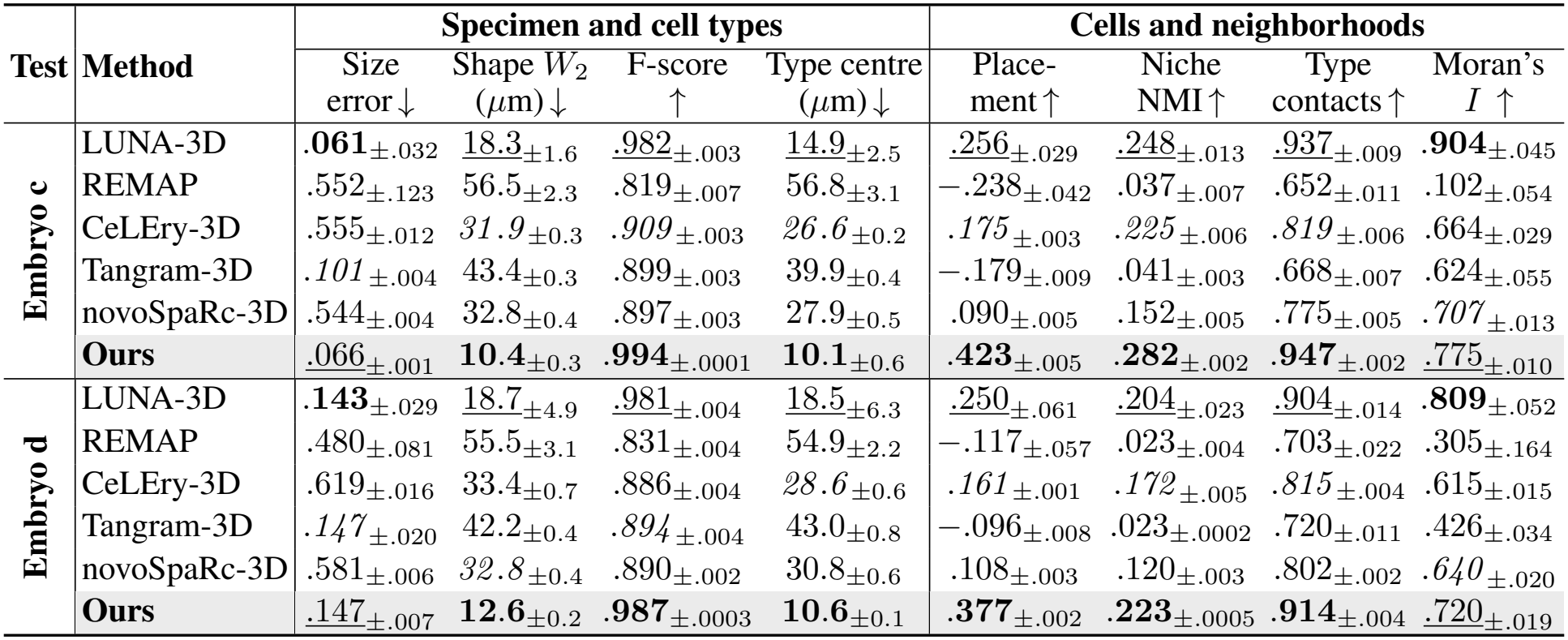
Per-test results: *Drosophila* embryos (columns and marks as in Table S4).

| **Test** | **Method** | **Specimen and cell types** |  |  |  | **Cells and neighborhoods** |  |  |  |
|---|---|---|---|---|---|---|---|---|---|
|  |  | Size error↓ | Shape *W*<sub>2</sub> (*µ*m)↓ | F-score ↑ | Type centre (*µ*m)↓ | Place-ment↑ | Niche NMI↑ | Type contacts↑ | Moran's *I* ↑ |
| **Embryo c** | LUNA-3D | **.061**<sub>±.032</sub> | 18.3<sub>±1.6</sub> | .982<sub>±.003</sub> | 14.9<sub>±2.5</sub> | .256<sub>±.029</sub> | .248<sub>±.013</sub> | .937<sub>±.009</sub> | **.904**<sub>±.045</sub> |
|  | REMAP | .552<sub>±.123</sub> | 56.5<sub>±2.3</sub> | .819<sub>±.007</sub> | 56.8<sub>±3.1</sub> | −.238<sub>±.042</sub> | .037<sub>±.007</sub> | .652<sub>±.011</sub> | .102<sub>±.054</sub> |
|  | CeLEry-3D | .555<sub>±.012</sub> | *31.9*<sub>±0.3</sub> | *.909*<sub>±.003</sub> | *26.6*<sub>±0.2</sub> | *.175*<sub>±.003</sub> | *.225*<sub>±.006</sub> | *.819*<sub>±.006</sub> | .664<sub>±.029</sub> |
|  | Tangram-3D | *.101*<sub>±.004</sub> | 43.4<sub>±0.3</sub> | .899<sub>±.003</sub> | 39.9<sub>±0.4</sub> | −.179<sub>±.009</sub> | .041<sub>±.003</sub> | .668<sub>±.007</sub> | .624<sub>±.055</sub> |
|  | novoSpaRc-3D | .544<sub>±.004</sub> | 32.8<sub>±0.4</sub> | .897<sub>±.003</sub> | 27.9<sub>±0.5</sub> | .090<sub>±.005</sub> | .152<sub>±.005</sub> | .775<sub>±.005</sub> | *.707*<sub>±.013</sub> |
|  | **Ours** | .066<sub>±.001</sub> | **10.4**<sub>±0.3</sub> | **.994**<sub>±.0001</sub> | **10.1**<sub>±0.6</sub> | **.423**<sub>±.005</sub> | **.282**<sub>±.002</sub> | **.947**<sub>±.002</sub> | .775<sub>±.010</sub> |
| **Embryo d** | LUNA-3D | **.143**<sub>±.029</sub> | 18.7<sub>±4.9</sub> | .981<sub>±.004</sub> | 18.5<sub>±6.3</sub> | .250<sub>±.061</sub> | .204<sub>±.023</sub> | .904<sub>±.014</sub> | **.809**<sub>±.052</sub> |
|  | REMAP | .480<sub>±.081</sub> | 55.5<sub>±3.1</sub> | .831<sub>±.004</sub> | 54.9<sub>±2.2</sub> | −.117<sub>±.057</sub> | .023<sub>±.004</sub> | .703<sub>±.022</sub> | .305<sub>±.164</sub> |
|  | CeLEry-3D | .619<sub>±.016</sub> | 33.4<sub>±0.7</sub> | .886<sub>±.004</sub> | *28.6*<sub>±0.6</sub> | *.161*<sub>±.001</sub> | *.172*<sub>±.005</sub> | *.815*<sub>±.004</sub> | .615<sub>±.015</sub> |
|  | Tangram-3D | *.147*<sub>±.020</sub> | 42.2<sub>±0.4</sub> | *.894*<sub>±.004</sub> | 43.0<sub>±0.8</sub> | −.096<sub>±.008</sub> | .023<sub>±.0002</sub> | .720<sub>±.011</sub> | .426<sub>±.034</sub> |
|  | novoSpaRc-3D | .581<sub>±.006</sub> | *32.8*<sub>±0.4</sub> | .890<sub>±.002</sub> | 30.8<sub>±0.6</sub> | .108<sub>±.003</sub> | .120<sub>±.003</sub> | .802<sub>±.002</sub> | *.640*<sub>±.020</sub> |
|  | **Ours** | .147<sub>±.007</sub> | **12.6**<sub>±0.2</sub> | **.987**<sub>±.0003</sub> | **10.6**<sub>±0.1</sub> | **.377**<sub>±.002</sub> | **.223**<sub>±.0005</sub> | **.914**<sub>±.004</sub> | .720<sub>±.019</sub> |

**Table S7:** One network call versus the full sampler. A single call of our network from pure noise returns its estimate of the average of the plausible configurations (the best guess); the 50-step sample is the Full Model’s sample before reseating. Error = median distance between predicted and true positions; matched = the same after size matching; squeezed = share of fine cell types (≥ 200 cells) whose spread after size matching is below 0.8 of the truth; SRC = LUNA’s per-cell Spearman correlation of distances; RSSD = LUNA’s sum over cell classes of the residual after rotation (it grows with the number of cells, so it compares methods within a test). Ours: final model of each test, mean of four draws; LUNA-3D: mean of 3 runs with different seeds. **Bold**: best of the three within a test.

| Test | Output | Per cell |  | Size and shape |  |  | Niche<br>NMI↑ | 2-D metrics |  |
|---|---|---|---|---|---|---|---|---|---|
|  |  | Error<br>(*µ*m)↓ | Matched<br>(*µ*m)↓ | *W*<sub>2</sub><br>(*µ*m)↓ | Size<br>err.↓ | Squeezed<br>(%)↓ |  | SRC<br>↑ | RSSD<br>(mm)↓ |
| Mouse 3 | LUNA-3D | 1,398 | 1,380 | 385 | **.035** | 22.6 | **.450** | .766 | 10,145 |
|  | One call | **1,173** | **1,233** | 334 | .106 | 18.9 | .444 | **.809** | **8,272** |
|  | Sample | 1,432 | 1,389 | **211** | .058 | **5.2** | .441 | .753 | 10,548 |
| Mouse 2 | LUNA-3D | 1,634 | 1,602 | 409 | .181 | 11.2 | .398 | .682 | 10,424 |
|  | One call | **1,254** | **1,352** | 391 | .157 | 15.5 | **.413** | **.758** | **7,777** |
|  | Sample | 1,537 | 1,530 | **202** | **.024** | **3.3** | .391 | .687 | 10,028 |
| Mouse 1 | LUNA-3D | 1,799 | 1,855 | 523 | .108 | 8.3 | .369 | .636 | 16,768 |
|  | One call | **1,227** | **1,300** | 297 | .150 | 13.5 | **.410** | **.772** | **12,080** |
|  | Sample | 1,453 | 1,478 | **163** | **.039** | **3.2** | .393 | .723 | 14,499 |
| Allen | LUNA-3D | 1,216 | 1,240 | 407 | .095 | 21.2 | **.500** | .782 | 15,049 |
|  | One call | **1,102** | **1,130** | 282 | .118 | 11.3 | .468 | **.795** | **13,866** |
|  | Sample | 1,317 | 1,339 | **211** | **.050** | **1.6** | .459 | .729 | 16,419 |
| Deep-STARmap A | LUNA-3D | 1,065 | 1,111 | 284 | .268 | **0.6** | .215 | .283 | 2,500 |
|  | One call | 797 | 832 | 232 | .311 | 9.5 | .313 | **.457** | **2,245** |
|  | Sample | **789** | **819** | **226** | **.097** | 7.8 | **.346** | .441 | 2,315 |
| Deep-STARmap B | LUNA-3D | 1,008 | 1,047 | 341 | .259 | 2.8 | .233 | .372 | 2,465 |
|  | One call | **725** | 753 | 199 | .138 | 0.8 | .346 | **.533** | **2,160** |
|  | Sample | 729 | **744** | **197** | **.043** | **0.0** | **.379** | .508 | 2,291 |
| Embryo c | LUNA-3D | 74.2 | 73.6 | 18.3 | **.061** | **0.0** | .248 | .452 | 79 |
|  | One call | **57.0** | 59.0 | 11.8 | .105 | **0.0** | .275 | **.620** | **60** |
|  | Sample | 57.3 | **58.5** | **10.7** | .064 | **0.0** | **.277** | .614 | 63 |
| Embryo d | LUNA-3D | 89.4 | 93.0 | 18.7 | **.143** | 6.1 | .204 | .382 | 107 |
|  | One call | **72.9** | 76.4 | 14.0 | .203 | **0.0** | .221 | **.550** | **85** |
|  | Sample | 73.2 | **76.0** | **12.9** | .154 | 6.8 | **.230** | .542 | 90 |

### S6 Ablations and robustness

Table S8 lists every ablation and robustness experiment on every test where it was run. Each training variant is a separate training of our model that differs from the final recipe in one choice and is otherwise identical, including the number of epochs and the global batch; it was run on the tests where that choice acts, the mice for choices about whole brains and the embryos for choices about the views and the left–right fold, and is compared with the training of our final model that has the same seed (seed 0), shown in the first block of the table as the w/o-reseating row (the Full Model before its reseating step). Each sampler variant changes one setting of the sampler on that same trained model and is compared with it under the default sampler: 50 steps, one shared state for the whole specimen, windows of 20,000 cells redrawn at every step, and the default step times. Each input variant degrades the input of the same model: zeroing a random 25% or 50% of the genes, thinning every cell’s transcript counts binomially to 50% or 25%, or keeping a random 50%, 25% or 10% of the cells. Every variant is sampled four times with different sampler seeds and scored before reseating with the same scorer as the main results, so that it is compared with the Full Model before reseating; the effect of reseating itself is the second row of the first block. The pairwise-distance loss and the independent chunks, which use the chunk sampler, rest on one draw. The variants of Table 2 are also reseated in the same way as the Full Model, one draw each on mice 2 and 3, and compared with the Full Model (second block); their means over the two mice are the values of Table 2. A model trained with the pairwise-distance loss predicts positions in a frame of its own at every step and so cannot update one shared state; it is sampled in chunks of about 19,000 cells that are aligned to each other on shared cells, and the same chunk sampler applied to our final model (independent chunks) separates the effect of the loss from that of the sampler. The count-thinning variant is evaluated under thinned counts against our final model under the same counts, since thinning in training is meant for shallow input. A subset of cells placed alone is compared with the same cells taken from the reconstruction of the whole specimen, so that both are scored on identical cells.

The experiments separate the choices that carry the result from the settings that do not. Sampling with one shared state is what keeps the size and shape of the specimen: one or two steps approximate the average layout and raise the size error 1.3–6.6-fold on every test, improving placement by 15–21% on the brains and changing it by at most 1% elsewhere, and independent chunks are worse than our sampler on all four measures on both mice. After reseating (second block), niche agreement under every sampler setting and with the unit box stays within 2% of the Full Model on both mice, while the single call, the independent chunks and the pairwise-distance loss stay 12–20% below it. Redrawing the windows at every step matters for the continuous blocks, whose size error rises 2.0- and 3.0-fold with fixed windows, and fixed windows lower niche agreement by 3–6% on every test. Among the training choices, the pairwise-distance loss and the unit box lose the size of the specimen on both mice, and the unit box also loses 77% of the placement on mouse 3. Without the left–right fold, the size error of the embryos rises by 25–43% and their shape error by 2–9%. A smaller network (four layers of width 128) raises the shape error of mouse 2 by 9% and lowers niche agreement by 6–9% on all three tests where it was run, and training on three or six of the eleven embryos raises their shape error by 26–61% and 4–10%. A larger network, one or two training brains instead of three, whole-specimen views only, the number of cells per view and the view label worsen the shape error by at most 2%, and the time distribution changes it by at most 12% in either direction. With ten or more steps, any window size and either step spacing, the shape error stays within 5% of the default on every test and placement and niche agreement worsen by at most 3%, while the size error, a small quantity, varies up to twofold in either direction. Degrading the input lowers the quality gradually: halving the counts raises the shape error by at most 19%, zeroing a quarter of the genes by up to 84% (mouse 1), and a model trained without count thinning loses more under thinned counts than our model does. Placing a subset of the cells alone gives the same reconstruction of those cells as placing all of them.

**Table S8:** Every ablation and robustness experiment. All rows are our final training (seed 0) or a training with the same seed. Rows are scored before reseating, as means over four sampler draws (pairwise-distance loss and independent chunks: one draw), and compared with the w/o-reseating row of the same test (the Full Model’s samples before its reseating step, first block); the exceptions are the Full Model itself and the second block, in which the first draw of each variant is reseated like the Full Model and compared with the Full Model. The default sampler takes 50 steps from one shared state, with windows of 20,000 cells redrawn at every step. Superscripts: % change, green better, red worse. ^*a*^Compared with the Full Model’s training under the same thinned counts (rows above). In the last block, a random subset of the cells is placed alone and compared with the same cells placed among all the others (row above). Shape *W*_2_ in *μ*m.

| Change | Test | Size error ↓ | Shape *W*<sub>2</sub> ↓ | Placement ↑ | Niche NMI ↑ |
|---|---|---|---|---|---|
| *Reference: the Full Model and its samples before reseating* |  |  |  |  |  |
| w/o reseating | Mouse 1 | .039 | 163 | .503 | .393 |
|  | Mouse 2 | .024 | 202 | .465 | .391 |
|  | Mouse 3 | .058 | 211 | .458 | .441 |
|  | Allen | .050 | 211 | .471 | .459 |
|  | Block A | .097 | 226 | .460 | .346 |
|  | Block B | .043 | 197 | .444 | .379 |
|  | Embryo c | .064 | 10.7 | .425 | .277 |
|  | Embryo d | .154 | 12.9 | .385 | .230 |
| **Full Model** | Mouse 1 | .039<sup>0</sup> | 163<sup>0</sup> | .499<sup>−1</sup> | .461<sup>+17</sup> |
|  | Mouse 2 | .024<sup>0</sup> | 202<sup>0</sup> | .458<sup>−2</sup> | .470<sup>+20</sup> |
|  | Mouse 3 | .058<sup>0</sup> | 208<sup>−2</sup> | .450<sup>−2</sup> | .513<sup>+16</sup> |
|  | Allen | .050<sup>0</sup> | 212<sup>+1</sup> | .465<sup>−1</sup> | .480<sup>+4</sup> |
|  | Block A | .097<sup>0</sup> | 224<sup>−1</sup> | .456<sup>−1</sup> | .359<sup>+4</sup> |
|  | Block B | .043<sup>+1</sup> | 197<sup>0</sup> | .438<sup>−1</sup> | .391<sup>+3</sup> |
|  | Embryo c | .065<sup>+2</sup> | 10.8<sup>0</sup> | .421<sup>−1</sup> | .281<sup>+1</sup> |
|  | Embryo d | .153<sup>0</sup> | 12.8<sup>0</sup> | .379<sup>−2</sup> | .223<sup>−3</sup> |
| *After reseating (compared with the Full Model)* |  |  |  |  |  |
| 1 step (single call) | Mouse 2 | .158<sup>+567</sup> | 391<sup>+94</sup> | .564<sup>+23</sup> | .414<sup>−12</sup> |
|  | Mouse 3 | .106<sup>+85</sup> | 335<sup>+61</sup> | .557<sup>+24</sup> | .444<sup>−13</sup> |
| Independent chunks | Mouse 2 | .228<sup>+863</sup> | 383<sup>+89</sup> | .373<sup>−19</sup> | .391<sup>−17</sup> |
|  | Mouse 3 | .366<sup>+537</sup> | 487<sup>+135</sup> | .261<sup>−42</sup> | .420<sup>−18</sup> |
| Pairwise-distance loss (chunk-sampled) | Mouse 2 | .216<sup>+815</sup> | 431<sup>+113</sup> | .371<sup>−19</sup> | .407<sup>−13</sup> |
|  | Mouse 3 | .322<sup>+460</sup> | 578<sup>+178</sup> | .214<sup>−52</sup> | .409<sup>−20</sup> |
| Unit-box coordinates | Mouse 2 | .047<sup>+100</sup> | 202<sup>0</sup> | .464<sup>+1</sup> | .462<sup>−2</sup> |
|  | Mouse 3 | .460<sup>+699</sup> | 223<sup>+7</sup> | .098<sup>−78</sup> | .508<sup>−1</sup> |
| 25 steps | Mouse 2 | .022<sup>−7</sup> | 201<sup>−1</sup> | .462<sup>+1</sup> | .468<sup>0</sup> |
|  | Mouse 3 | .053<sup>−8</sup> | 204<sup>−2</sup> | .453<sup>+1</sup> | .512<sup>0</sup> |
| 100 steps | Mouse 2 | .024<sup>+3</sup> | 204<sup>+1</sup> | .457<sup>0</sup> | .471<sup>0</sup> |
|  | Mouse 3 | .060<sup>+4</sup> | 209<sup>+1</sup> | .448<sup>0</sup> | .513<sup>0</sup> |
| Windows of 5,000 cells | Mouse 2 | .024<sup>+1</sup> | 203<sup>0</sup> | .458<sup>0</sup> | .469<sup>0</sup> |
|  | Mouse 3 | .057<sup>−1</sup> | 207<sup>0</sup> | .449<sup>0</sup> | .512<sup>0</sup> |
| Windows of 40,000 cells | Mouse 2 | .023<sup>−1</sup> | 203<sup>0</sup> | .458<sup>0</sup> | .470<sup>0</sup> |
|  | Mouse 3 | .058<sup>+1</sup> | 207<sup>0</sup> | .449<sup>0</sup> | .513<sup>0</sup> |
| Uniform step times | Mouse 2 | .014<sup>−42</sup> | 197<sup>−3</sup> | .468<sup>+2</sup> | .464<sup>−1</sup> |
|  | Mouse 3 | .031<sup>−47</sup> | 200<sup>−4</sup> | .461<sup>+3</sup> | .509<sup>−1</sup> |
| *Training variants* |  |  |  |  |  |
| Pairwise-distance loss (chunk-sampled) | Mouse 2 | .216<sup>+813</sup> | 435<sup>+115</sup> | .376<sup>−19</sup> | .321<sup>−18</sup> |
|  | Mouse 3 | .322<sup>+460</sup> | 582<sup>+176</sup> | .218<sup>−52</sup> | .305<sup>−31</sup> |
| Unit-box coordinates | Mouse 2 | .047<sup>+100</sup> | 202<sup>0</sup> | .470<sup>+1</sup> | .382<sup>−2</sup> |
|  | Mouse 3 | .459<sup>+697</sup> | 224<sup>+6</sup> | .103<sup>−77</sup> | .435<sup>−1</sup> |
| Whole-specimen views only | Mouse 3 | .034<sup>−42</sup> | 206<sup>−3</sup> | .477<sup>+4</sup> | .441<sup>0</sup> |
| No count thinning | Mouse 2 | .022<sup>−6</sup> | 204<sup>+1</sup> | .452<sup>−3</sup> | .384<sup>−2</sup> |
| Single time distribution | Mouse 2 | .013<sup>−46</sup> | 217<sup>+7</sup> | .492<sup>+6</sup> | .398<sup>+2</sup> |
|  | Embryo c | .078<sup>+22</sup> | 9.5<sup>−12</sup> | .436<sup>+2</sup> | .277<sup>0</sup> |
|  | Embryo d | .158<sup>+3</sup> | 12.9<sup>+1</sup> | .370<sup>−4</sup> | .219<sup>−5</sup> |
| Smaller network (4 × 128) | Mouse 2 | .035<sup>+48</sup> | 221<sup>+9</sup> | .425<sup>−9</sup> | .366<sup>−6</sup> |
|  | Embryo c | .057<sup>−11</sup> | 10.4<sup>−3</sup> | .407<sup>−4</sup> | .257<sup>−7</sup> |
|  | Embryo d | .142<sup>−8</sup> | 12.1<sup>−6</sup> | .371<sup>−4</sup> | .209<sup>−9</sup> |
| Larger network (12 × 384) | Mouse 2 | .021<sup>−10</sup> | 198<sup>−2</sup> | .468<sup>+1</sup> | .391<sup>0</sup> |
| 1 training brain (mouse 1) | Mouse 2 | .037<sup>+55</sup> | 206<sup>+2</sup> | .454<sup>−2</sup> | .390<sup>0</sup> |
| 2 training brains (mice 1, 3) | Mouse 2 | .015<sup>−36</sup> | 199<sup>−2</sup> | .459<sup>−1</sup> | .383<sup>−2</sup> |
| 3 training embryos | Embryo c | .068<sup>+7</sup> | 17.3<sup>+61</sup> | .317<sup>−26</sup> | .285<sup>+3</sup> |
|  | Embryo d | .160<sup>+4</sup> | 16.3<sup>+26</sup> | .314<sup>−19</sup> | .227<sup>−1</sup> |
| 6 training embryos | Embryo c | .050<sup>−21</sup> | 11.1<sup>+4</sup> | .416<sup>−2</sup> | .288<sup>+4</sup> |
|  | Embryo d | .144<sup>−6</sup> | 14.2<sup>+10</sup> | .367<sup>−5</sup> | .240<sup>+4</sup> |
| No view label | Mouse 3 | .041<sup>−28</sup> | 213<sup>+1</sup> | .465<sup>+1</sup> | .440<sup>0</sup> |
| 5,000–10,000 cells per view | Embryo c | .077<sup>+21</sup> | 10.0<sup>−7</sup> | .441<sup>+4</sup> | .287<sup>+4</sup> |
|  | Embryo d | .163<sup>+6</sup> | 12.7<sup>−1</sup> | .386<sup>0</sup> | .237<sup>+3</sup> |
| 20,000–40,000 cells per view | Embryo c | .053<sup>−17</sup> | 9.8<sup>−9</sup> | .432<sup>+2</sup> | .282<sup>+2</sup> |
|  | Embryo d | .136<sup>−12</sup> | 12.3<sup>−5</sup> | .388<sup>+1</sup> | .239<sup>+4</sup> |
| No left–right fold | Embryo c | .091<sup>+43</sup> | 10.9<sup>+2</sup> | .450<sup>+6</sup> | .293<sup>+6</sup> |
|  | Embryo d | .192<sup>+25</sup> | 14.0<sup>+9</sup> | .388<sup>+1</sup> | .233<sup>+1</sup> |
| *Sampler variants (the Full Model's training)* |  |  |  |  |  |
| 1 step (single call) | Mouse 1 | .150<sup>+282</sup> | 297<sup>+82</sup> | .581<sup>+15</sup> | .410<sup>+4</sup> |
|  | Mouse 2 | .157<sup>+564</sup> | 391<sup>+93</sup> | .564<sup>+21</sup> | .413<sup>+6</sup> |
|  | Mouse 3 | .106<sup>+84</sup> | 334<sup>+58</sup> | .556<sup>+21</sup> | .444<sup>+1</sup> |
|  | Allen | .118<sup>+138</sup> | 282<sup>+34</sup> | .557<sup>+18</sup> | .468<sup>+2</sup> |
|  | Block A | .311<sup>+222</sup> | 232<sup>+3</sup> | .455<sup>−1</sup> | .313<sup>−10</sup> |
|  | Block B | .138<sup>+219</sup> | 199<sup>+1</sup> | .447<sup>+1</sup> | .346<sup>−9</sup> |
|  | Embryo c | .105<sup>+65</sup> | 11.8<sup>+10</sup> | .428<sup>+1</sup> | .275<sup>−1</sup> |
|  | Embryo d | .203<sup>+32</sup> | 14.0<sup>+9</sup> | .388<sup>+1</sup> | .221<sup>−4</sup> |
| 2 steps | Mouse 1 | .124<sup>+216</sup> | 282<sup>+73</sup> | .582<sup>+16</sup> | .415<sup>+5</sup> |
|  | Mouse 2 | .139<sup>+486</sup> | 376<sup>+86</sup> | .565<sup>+21</sup> | .414<sup>+6</sup> |
|  | Mouse 3 | .097<sup>+68</sup> | 325<sup>+54</sup> | .556<sup>+21</sup> | .448<sup>+2</sup> |
|  | Allen | .120<sup>+142</sup> | 282<sup>+34</sup> | .560<sup>+19</sup> | .475<sup>+3</sup> |
|  | Block A | .255<sup>+164</sup> | 232<sup>+3</sup> | .457<sup>−1</sup> | .315<sup>−9</sup> |
|  | Block B | .111<sup>+156</sup> | 199<sup>+1</sup> | .448<sup>+1</sup> | .347<sup>−8</sup> |
|  | Embryo c | .107<sup>+67</sup> | 11.8<sup>+10</sup> | .428<sup>+1</sup> | .275<sup>−1</sup> |
|  | Embryo d | .205<sup>+33</sup> | 14.0<sup>+9</sup> | .388<sup>+1</sup> | .221<sup>−4</sup> |
| 5 steps | Mouse 1 | .074<sup>+89</sup> | 191<sup>+17</sup> | .542<sup>+8</sup> | .413<sup>+5</sup> |
|  | Mouse 2 | .054<sup>+126</sup> | 235<sup>+16</sup> | .505<sup>+9</sup> | .399<sup>+2</sup> |
|  | Mouse 3 | .022<sup>−62</sup> | 222<sup>+6</sup> | .511<sup>+11</sup> | .458<sup>+4</sup> |
|  | Allen | .123<sup>+147</sup> | 245<sup>+16</sup> | .551<sup>+17</sup> | .512<sup>+12</sup> |
|  | Block A | .092<sup>−5</sup> | 228<sup>+1</sup> | .465<sup>+1</sup> | .339<sup>−2</sup> |
|  | Block B | .099<sup>+130</sup> | 198<sup>0</sup> | .456<sup>+3</sup> | .378<sup>0</sup> |
|  | Embryo c | .095<sup>+49</sup> | 11.4<sup>+6</sup> | .432<sup>+1</sup> | .288<sup>+4</sup> |
|  | Embryo d | .192<sup>+25</sup> | 13.4<sup>+4</sup> | .390<sup>+1</sup> | .239<sup>+4</sup> |
| *Sampler variants (the Full Model's training) (continued)* |  |  |  |  |  |
| 10 steps | Mouse 1 | .051<sup>+30</sup> | 171<sup>+5</sup> | .521<sup>+3</sup> | .397<sup>+1</sup> |
|  | Mouse 2 | .019<sup>−19</sup> | 204<sup>+1</sup> | .481<sup>+3</sup> | .383<sup>−2</sup> |
|  | Mouse 3 | .030<sup>−48</sup> | 206<sup>−3</sup> | .477<sup>+4</sup> | .437<sup>−1</sup> |
|  | Allen | .097<sup>+95</sup> | 220<sup>+4</sup> | .522<sup>+11</sup> | .502<sup>+9</sup> |
|  | Block A | .127<sup>+32</sup> | 227<sup>0</sup> | .464<sup>+1</sup> | .347<sup>0</sup> |
|  | Block B | .070<sup>+63</sup> | 197<sup>0</sup> | .451<sup>+2</sup> | .379<sup>0</sup> |
|  | Embryo c | .082<sup>+28</sup> | 11.0<sup>+3</sup> | .430<sup>+1</sup> | .283<sup>+2</sup> |
|  | Embryo d | .176<sup>+15</sup> | 13.1<sup>+2</sup> | .389<sup>+1</sup> | .236<sup>+3</sup> |
| 25 steps | Mouse 1 | .041<sup>+5</sup> | 163<sup>0</sup> | .507<sup>+1</sup> | .394<sup>0</sup> |
|  | Mouse 2 | .022<sup>−6</sup> | 200<sup>−1</sup> | .468<sup>+1</sup> | .389<sup>−1</sup> |
|  | Mouse 3 | .053<sup>−8</sup> | 207<sup>−2</sup> | .461<sup>+1</sup> | .440<sup>0</sup> |
|  | Allen | .064<sup>+30</sup> | 210<sup>0</sup> | .486<sup>+3</sup> | .473<sup>+3</sup> |
|  | Block A | .108<sup>+12</sup> | 227<sup>0</sup> | .461<sup>0</sup> | .347<sup>0</sup> |
|  | Block B | .052<sup>+21</sup> | 197<sup>0</sup> | .446<sup>+1</sup> | .380<sup>0</sup> |
|  | Embryo c | .069<sup>+8</sup> | 10.8<sup>+1</sup> | .427<sup>0</sup> | .279<sup>+1</sup> |
|  | Embryo d | .160<sup>+4</sup> | 12.9<sup>0</sup> | .387<sup>0</sup> | .232<sup>+1</sup> |
| 100 steps | Mouse 1 | .039<sup>−1</sup> | 163<sup>0</sup> | .501<sup>0</sup> | .393<sup>0</sup> |
|  | Mouse 2 | .025<sup>+4</sup> | 204<sup>+1</sup> | .464<sup>0</sup> | .392<sup>0</sup> |
|  | Mouse 3 | .059<sup>+3</sup> | 213<sup>+1</sup> | .456<sup>0</sup> | .441<sup>0</sup> |
|  | Allen | .041<sup>−17</sup> | 212<sup>+1</sup> | .463<sup>−2</sup> | .452<sup>−2</sup> |
|  | Block A | .089<sup>−7</sup> | 226<sup>0</sup> | .460<sup>0</sup> | .345<sup>0</sup> |
|  | Block B | .038<sup>−12</sup> | 197<sup>0</sup> | .443<sup>0</sup> | .378<sup>0</sup> |
|  | Embryo c | .061<sup>−5</sup> | 10.7<sup>0</sup> | .425<sup>0</sup> | .276<sup>0</sup> |
|  | Embryo d | .150<sup>−3</sup> | 12.9<sup>0</sup> | .385<sup>0</sup> | .229<sup>0</sup> |
| Independent chunks | Mouse 2 | .228<sup>+860</sup> | 387<sup>+91</sup> | .377<sup>−19</sup> | .297<sup>−24</sup> |
|  | Mouse 3 | .366<sup>+537</sup> | 492<sup>+133</sup> | .265<sup>−42</sup> | .327<sup>−26</sup> |
| Fixed windows | Mouse 1 | .039<sup>0</sup> | 162<sup>0</sup> | .499<sup>−1</sup> | .383<sup>−3</sup> |
|  | Mouse 2 | .025<sup>+5</sup> | 201<sup>0</sup> | .461<sup>−1</sup> | .378<sup>−3</sup> |
|  | Mouse 3 | .059<sup>+3</sup> | 209<sup>−1</sup> | .454<sup>−1</sup> | .429<sup>−3</sup> |
|  | Allen | .048<sup>−4</sup> | 212<sup>0</sup> | .465<sup>−1</sup> | .434<sup>−6</sup> |
|  | Block A | .197<sup>+105</sup> | 226<sup>0</sup> | .458<sup>−1</sup> | .336<sup>−3</sup> |
|  | Block B | .130<sup>+201</sup> | 197<sup>0</sup> | .441<sup>−1</sup> | .369<sup>−3</sup> |
| Uniform step times | Mouse 1 | .055<sup>+41</sup> | 162<sup>−1</sup> | .510<sup>+1</sup> | .390<sup>−1</sup> |
|  | Mouse 2 | .014<sup>−40</sup> | 196<sup>−3</sup> | .474<sup>+2</sup> | .384<sup>−2</sup> |
|  | Mouse 3 | .031<sup>−47</sup> | 202<sup>−4</sup> | .469<sup>+2</sup> | .436<sup>−1</sup> |
|  | Allen | .057<sup>+16</sup> | 211<sup>0</sup> | .496<sup>+5</sup> | .506<sup>+10</sup> |
|  | Block A | .107<sup>+11</sup> | 227<sup>0</sup> | .462<sup>0</sup> | .349<sup>+1</sup> |
|  | Block B | .087<sup>+101</sup> | 198<sup>0</sup> | .447<sup>+1</sup> | .387<sup>+2</sup> |
|  | Embryo c | .065<sup>+1</sup> | 10.8<sup>0</sup> | .431<sup>+1</sup> | .286<sup>+3</sup> |
|  | Embryo d | .155<sup>+1</sup> | 12.9<sup>+1</sup> | .389<sup>+1</sup> | .239<sup>+4</sup> |
| Windows of 5,000 cells | Mouse 1 | .040<sup>+2</sup> | 163<sup>0</sup> | .503<sup>0</sup> | .393<sup>0</sup> |
|  | Mouse 2 | .024<sup>+2</sup> | 202<sup>0</sup> | .465<sup>0</sup> | .390<sup>0</sup> |
|  | Mouse 3 | .057<sup>−1</sup> | 211<sup>0</sup> | .458<sup>0</sup> | .440<sup>0</sup> |
|  | Allen | .047<sup>−5</sup> | 211<sup>0</sup> | .467<sup>−1</sup> | .447<sup>−3</sup> |
|  | Block A | .094<sup>−3</sup> | 226<sup>0</sup> | .460<sup>0</sup> | .346<sup>0</sup> |
|  | Block B | .040<sup>−8</sup> | 197<sup>0</sup> | .444<sup>0</sup> | .378<sup>0</sup> |
| Windows of 10,000 cells | Mouse 1 | .040<sup>+1</sup> | 163<sup>0</sup> | .503<sup>0</sup> | .393<sup>0</sup> |
|  | Mouse 2 | .024<sup>+1</sup> | 202<sup>0</sup> | .465<sup>0</sup> | .391<sup>0</sup> |
|  | Mouse 3 | .057<sup>−1</sup> | 211<sup>0</sup> | .458<sup>0</sup> | .441<sup>0</sup> |
|  | Allen | .049<sup>−2</sup> | 211<sup>0</sup> | .469<sup>0</sup> | .455<sup>−1</sup> |
|  | Block A | .096<sup>−1</sup> | 226<sup>0</sup> | .460<sup>0</sup> | .346<sup>0</sup> |
|  | Block B | .042<sup>−2</sup> | 197<sup>0</sup> | .444<sup>0</sup> | .379<sup>0</sup> |
| *Sampler variants (the Full Model's training) (continued)* |  |  |  |  |  |
| Windows of 40,000 cells | Mouse 1 | .039<sup>0</sup> | 163<sup>0</sup> | .503<sup>0</sup> | .393<sup>0</sup> |
|  | Mouse 2 | .024<sup>0</sup> | 202<sup>0</sup> | .465<sup>0</sup> | .391<sup>0</sup> |
|  | Mouse 3 | .058<sup>0</sup> | 211<sup>0</sup> | .458<sup>0</sup> | .441<sup>0</sup> |
|  | Allen | .050<sup>+1</sup> | 211<sup>0</sup> | .472<sup>0</sup> | .462<sup>0</sup> |
|  | Block A | .097<sup>+1</sup> | 226<sup>0</sup> | .460<sup>0</sup> | .346<sup>0</sup> |
|  | Block B | .044<sup>+1</sup> | 197<sup>0</sup> | .444<sup>0</sup> | .379<sup>0</sup> |
| *Degraded input (the Full Model's training)* |  |  |  |  |  |
| 25% of genes zeroed | Mouse 1 | .130<sup>+229</sup> | 300<sup>+84</sup> | .441<sup>−12</sup> | .360<sup>−9</sup> |
|  | Mouse 2 | .031<sup>+32</sup> | 296<sup>+47</sup> | .375<sup>−19</sup> | .339<sup>−13</sup> |
|  | Mouse 3 | .079<sup>+37</sup> | 315<sup>+49</sup> | .366<sup>−20</sup> | .395<sup>−10</sup> |
|  | Block A | .085<sup>−12</sup> | 223<sup>−1</sup> | .394<sup>−14</sup> | .292<sup>−16</sup> |
|  | Block B | .066<sup>+52</sup> | 244<sup>+24</sup> | .344<sup>−22</sup> | .320<sup>−16</sup> |
|  | Embryo c | .062<sup>−3</sup> | 11.2<sup>+4</sup> | .395<sup>−7</sup> | .250<sup>−10</sup> |
|  | Embryo d | .148<sup>−4</sup> | 14.2<sup>+10</sup> | .356<sup>−8</sup> | .202<sup>−12</sup> |
| 50% of genes zeroed | Mouse 1 | .111<sup>+181</sup> | 483<sup>+197</sup> | .285<sup>−43</sup> | .288<sup>−27</sup> |
|  | Mouse 2 | .076<sup>+219</sup> | 487<sup>+141</sup> | .237<sup>−49</sup> | .290<sup>−26</sup> |
|  | Mouse 3 | .054<sup>−6</sup> | 484<sup>+130</sup> | .258<sup>−44</sup> | .346<sup>−22</sup> |
|  | Block A | .071<sup>−26</sup> | 292<sup>+29</sup> | .280<sup>−39</sup> | .212<sup>−39</sup> |
|  | Block B | .058<sup>+35</sup> | 325<sup>+65</sup> | .216<sup>−51</sup> | .235<sup>−38</sup> |
|  | Embryo c | .065<sup>+2</sup> | 18.2<sup>+70</sup> | .298<sup>−30</sup> | .218<sup>−21</sup> |
|  | Embryo d | .158<sup>+3</sup> | 19.1<sup>+49</sup> | .272<sup>−30</sup> | .173<sup>−25</sup> |
| Counts thinned to 50% | Mouse 1 | .046<sup>+16</sup> | 189<sup>+16</sup> | .415<sup>−18</sup> | .360<sup>−9</sup> |
|  | Mouse 2 | .029<sup>+23</sup> | 240<sup>+19</sup> | .370<sup>−20</sup> | .359<sup>−8</sup> |
|  | Mouse 3 | .062<sup>+7</sup> | 225<sup>+7</sup> | .372<sup>−19</sup> | .410<sup>−7</sup> |
|  | Block A | .053<sup>−45</sup> | 225<sup>0</sup> | .377<sup>−18</sup> | .291<sup>−16</sup> |
|  | Block B | .064<sup>+49</sup> | 233<sup>+18</sup> | .325<sup>−27</sup> | .305<sup>−20</sup> |
|  | Embryo c | .047<sup>−26</sup> | 12.4<sup>+15</sup> | .358<sup>−16</sup> | .236<sup>−15</sup> |
|  | Embryo d | .129<sup>−16</sup> | 13.6<sup>+6</sup> | .320<sup>−17</sup> | .187<sup>−19</sup> |
| Counts thinned to 25% | Mouse 1 | .090<sup>+128</sup> | 310<sup>+90</sup> | .294<sup>−42</sup> | .301<sup>−23</sup> |
|  | Mouse 2 | .045<sup>+90</sup> | 381<sup>+89</sup> | .251<sup>−46</sup> | .305<sup>−22</sup> |
|  | Mouse 3 | .044<sup>−23</sup> | 305<sup>+45</sup> | .272<sup>−41</sup> | .357<sup>−19</sup> |
|  | Block A | .159<sup>+65</sup> | 257<sup>+13</sup> | .235<sup>−49</sup> | .190<sup>−45</sup> |
|  | Block B | .286<sup>+562</sup> | 306<sup>+55</sup> | .143<sup>−68</sup> | .190<sup>−50</sup> |
|  | Embryo c | .047<sup>−26</sup> | 18.0<sup>+68</sup> | .225<sup>−47</sup> | .172<sup>−38</sup> |
|  | Embryo d | .142<sup>−8</sup> | 19.7<sup>+53</sup> | .200<sup>−48</sup> | .125<sup>−46</sup> |
| No count thinning, counts at 50%<sup>*a*</sup> | Mouse 2 | .030<sup>+4</sup> | 299<sup>+25</sup> | .362<sup>−2</sup> | .345<sup>−4</sup> |
| No count thinning, counts at 25%<sup>*a*</sup> | Mouse 2 | .140<sup>+211</sup> | 536<sup>+41</sup> | .236<sup>−6</sup> | .273<sup>−11</sup> |
| *Fewer cells (a subset placed alone)* |  |  |  |  |  |
| 50% of the cells, among all others | Mouse 1 | .040 | 165 | .504 | .393 |
|  | Mouse 2 | .024 | 204 | .465 | .386 |
|  | Mouse 3 | .057 | 213 | .458 | .444 |
|  | Allen | .050 | 212 | .471 | .458 |
|  | Block A | .097 | 228 | .463 | .341 |
|  | Block B | .045 | 200 | .442 | .379 |
|  | Embryo c | .067 | 10.9 | .438 | .280 |
|  | Embryo d | .151 | 13.7 | .384 | .232 |
| 50% of the cells, placed alone | Mouse 1 | .040<sup>0</sup> | 165<sup>0</sup> | .504<sup>0</sup> | .393<sup>0</sup> |
|  | Mouse 2 | .024<sup>−1</sup> | 205<sup>0</sup> | .464<sup>0</sup> | .386<sup>0</sup> |
|  | Mouse 3 | .057<sup>−1</sup> | 214<sup>+1</sup> | .459<sup>0</sup> | .444<sup>0</sup> |
|  | Allen | .050<sup>0</sup> | 211<sup>0</sup> | .471<sup>0</sup> | .458<sup>0</sup> |
|  | Block A | .095<sup>−2</sup> | 230<sup>+1</sup> | .461<sup>0</sup> | .342<sup>0</sup> |
|  | Block B | .045<sup>+1</sup> | 200<sup>0</sup> | .441<sup>0</sup> | .380<sup>0</sup> |
|  | Embryo c | .062<sup>−6</sup> | 10.8<sup>−1</sup> | .435<sup>−1</sup> | .282<sup>+1</sup> |
|  | Embryo d | .152<sup>0</sup> | 13.8<sup>+1</sup> | .389<sup>+1</sup> | .232<sup>0</sup> |
| 25% of the cells, among all others | Mouse 1 | .040 | 168 | .504 | .392 |
|  | Mouse 2 | .024 | 208 | .464 | .385 |
|  | Mouse 3 | .057 | 214 | .457 | .448 |
|  | Allen | .050 | 213 | .471 | .464 |
|  | Block A | .097 | 231 | .465 | .341 |
|  | Block B | .045 | 204 | .441 | .377 |
|  | Embryo c | .067 | 11.6 | .444 | .278 |
|  | Embryo d | .148 | 14.2 | .381 | .239 |
| 25% of the cells, placed alone | Mouse 1 | .040<sup>+1</sup> | 168<sup>0</sup> | .503<sup>0</sup> | .392<sup>0</sup> |
|  | Mouse 2 | .024<sup>−3</sup> | 208<sup>0</sup> | .464<sup>0</sup> | .385<sup>0</sup> |
|  | Mouse 3 | .057<sup>0</sup> | 215<sup>0</sup> | .458<sup>0</sup> | .447<sup>0</sup> |
|  | Allen | .049<sup>−1</sup> | 213<sup>0</sup> | .470<sup>0</sup> | .463<sup>0</sup> |
|  | Block A | .089<sup>−8</sup> | 232<sup>0</sup> | .464<sup>0</sup> | .341<sup>0</sup> |
|  | Block B | .035<sup>−21</sup> | 205<sup>+1</sup> | .439<sup>0</sup> | .375<sup>0</sup> |
|  | Embryo c | .068<sup>+1</sup> | 11.4<sup>−1</sup> | .437<sup>−2</sup> | .275<sup>−1</sup> |
|  | Embryo d | .150<sup>+1</sup> | 14.2<sup>0</sup> | .386<sup>+1</sup> | .240<sup>0</sup> |
| 10% of the cells, among all others | Mouse 1 | .040 | 173 | .504 | .401 |
|  | Mouse 2 | .025 | 216 | .463 | .387 |
|  | Mouse 3 | .058 | 216 | .457 | .451 |
|  | Allen | .050 | 219 | .470 | .469 |
|  | Block A | .096 | 248 | .464 | .339 |
|  | Block B | .042 | 218 | .436 | .375 |
|  | Embryo c | .077 | 13.6 | .431 | .282 |
|  | Embryo d | .150 | 16.6 | .381 | .231 |
| 10% of the cells, placed alone | Mouse 1 | .040<sup>0</sup> | 173<sup>0</sup> | .503<sup>0</sup> | .402<sup>0</sup> |
|  | Mouse 2 | .023<sup>−7</sup> | 216<sup>0</sup> | .465<sup>+1</sup> | .389<sup>+1</sup> |
|  | Mouse 3 | .059<sup>+2</sup> | 217<sup>0</sup> | .458<sup>0</sup> | .452<sup>0</sup> |
|  | Allen | .050<sup>0</sup> | 219<sup>0</sup> | .469<sup>0</sup> | .467<sup>0</sup> |
|  | Block A | .093<sup>−3</sup> | 250<sup>+1</sup> | .463<sup>0</sup> | .339<sup>0</sup> |
|  | Block B | .040<sup>−4</sup> | 219<sup>0</sup> | .434<sup>0</sup> | .376<sup>0</sup> |
|  | Embryo c | .073<sup>−5</sup> | 13.4<sup>−2</sup> | .431<sup>0</sup> | .284<sup>+1</sup> |
|  | Embryo d | .149<sup>0</sup> | 17.1<sup>+3</sup> | .387<sup>+2</sup> | .230<sup>−1</sup> |

### S7 Sampling versus averaging

Expression leaves each cell a distribution of possible positions with variance *σ*^2^. The position that minimizes the expected error of the cell is the mean of this distribution, and a layout made of these means is smaller than the tissue: by the law of total variance, its variance is that of the tissue minus the average *σ*^2^. A sample instead draws one position from each distribution and keeps the variance of the tissue; two independent draws of a cell lie 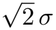 apart in expectation, and each lies 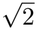 times as far from the truth as the mean does when the mean is exact. A single call of our network from pure noise estimates this mean, and the 50-step sampler draws a sample. Table S9 tests these relations on the final model of each test.

The one-call output is the center of the samples: the error of a sample equals 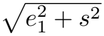, the error of the one-call output combined with the scatter of the samples around it, to within 2.1% on all eight tests, so sampling adds scatter around the one-call output without moving it; four seeds of the single call differ by 0.08–0.13 of its error. The one-call output is smaller than the tissue along every axis of the whole brains and embryos and along the two long axes of the blocks (0.76–0.98 of the true spread), and the sample restores size and shape: its size error and *W*_2_ are lower than those of the one-call output on all eight tests (Table S7). On the whole brains the samples scatter around the one-call output by 0.67–0.84 of its error, close to the ratio of 1 expected when that error is the spread of the possible positions, and the share of squeezed cell types falls from 11–19% to 2–5%.

**Table S9:** Sampling versus averaging. Size: standard deviation of the output over that of the truth along each axis. Errors are root-mean-square distances to the true position; scatter *s* is the root-mean-square distance between two samples of the same cell divided by 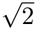. Last column: 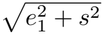, which equals *e*_50_ when the samples scatter around the one-call output. Final model of each test; four calls and four samples.

| **Test** | **Size (per axis)** |  | **Error (μm)** |  | **Scatter of the samples** |  |  |
|---|---|---|---|---|---|---|---|
|  | One call | Sample | One call *e*<sub>1</sub> | Sample *e*<sub>50</sub> | *s* (μm) | *s*/*e*<sub>1</sub> | $\sqrt{e_1^2 + s^2}$ |
| Mouse 3 | 0.86/0.86/0.97 | 1.10/1.05/1.02 | 1,841 | 2,372 | 1,472 | 0.80 | 2,357 |
| Mouse 2 | 0.79/0.83/0.91 | 1.00/1.05/0.98 | 2,018 | 2,638 | 1,699 | 0.84 | 2,638 |
| Mouse 1 | 0.77/0.86/0.92 | 0.92/1.00/0.97 | 2,032 | 2,456 | 1,394 | 0.69 | 2,465 |
| Allen | 0.86/0.86/0.93 | 0.96/0.93/0.96 | 1,864 | 2,195 | 1,242 | 0.67 | 2,240 |
| Deep-STARmap A | 0.88/0.89/1.70 | 0.89/0.90/0.93 | 1,368 | 1,411 | 404 | 0.30 | 1,427 |
| Deep-STARmap B | 0.91/0.98/1.30 | 0.94/1.01/0.94 | 1,284 | 1,364 | 487 | 0.38 | 1,373 |
| Embryo c | 0.91/0.87/0.91 | 0.95/0.91/0.95 | 92.9 | 98.2 | 32.3 | 0.35 | 98.4 |
| Embryo d | 0.83/0.76/0.80 | 0.88/0.81/0.85 | 107.7 | 114.3 | 38.6 | 0.36 | 114.4 |

#### The spread of the samples marks the cells to trust

Because our model is sampled rather than evaluated once, the disagreement between its samples measures its uncertainty without any ground truth, so it is available for dissociated cells, whose placement cannot be checked. We test whether it predicts the actual error on the three held-out mice. For every cell we take the spread of its positions in three samples, their mean distance from their average, and the error of its position in a fourth sample, the distance from its measured position, so that the two are measured on different samples. Spread predicts error on every mouse (Spearman correlation over cells 0.52, 0.56 and 0.51 on mice 1–3; Figure S1c; panels a and b show both for the cells of one section). Ranking the cells by spread turns one reconstruction into a sequence of increasingly reliable ones (Figure S1d): the 25% of cells with the lowest spread have a median error of 0.85, 0.84 and 0.89 mm, against 1.47, 1.56 and 1.46 mm for all cells and 2.82, 3.13 and 2.65 mm for the 20% with the highest spread, and keeping the 86, 76 and 74% most confident cells gives a lower median error than LUNA-3D over all its cells (1.32, 1.29 and 1.23 mm), which was run once and gives one position per cell.

**Figure S1:**
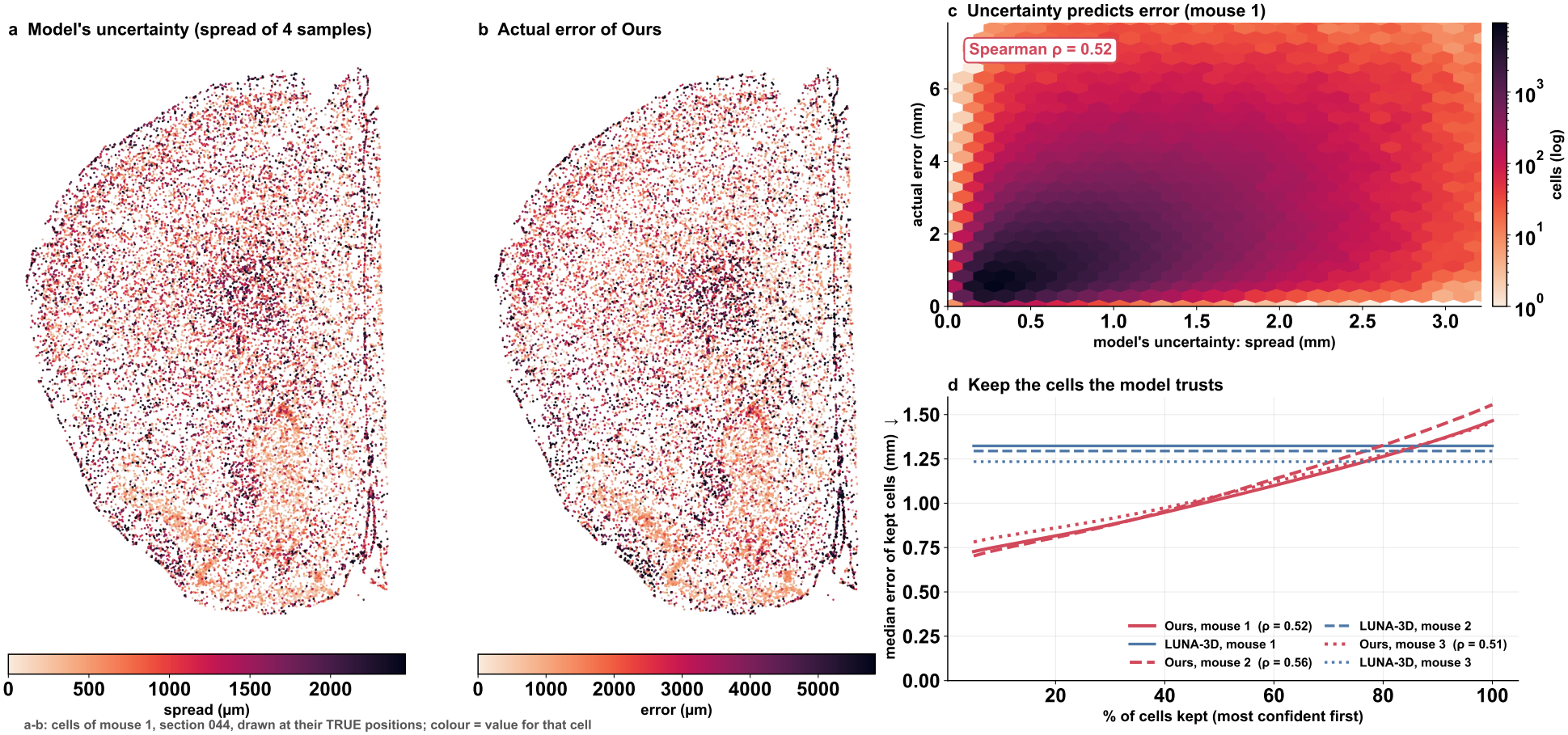
The spread of the samples predicts where the reconstruction is wrong. (a, b) Cells of one coronal section of mouse 1 (section 044, ±40 *μ*m) at their measured positions, colored by the spread of their positions over three samples (a) and by the error of their position in a fourth sample (b). (c) Spread against error for the cells of mouse 1 (300,000 random cells; color, number of cells on a log scale). (d) Median error of the cells kept, keeping cells in order of increasing spread, for mice 1–3; horizontal lines, the median error of LUNA-3D over all cells.

### S8 The noisy-layout test

Section S7 tells a sample from a best guess by the output of the sampler; the noisy-layout test asks the network itself. A sampler draws one configuration of a specimen only if its network uses the layout it is given, because every step must place each cell relative to the configuration drawn so far; a network that ignores the layout returns the same best guess per cell at every step, and the sampler ends at that guess. We take the true layout of the test cells, add Gaussian noise at a noise-to-signal ratio ranging from pure noise to almost none (the standard deviation of the noise along one axis over the root-mean-square spread of the true layout along one axis), and give it to the network together with the expression of the cells and the matching noise level. The network’s prediction of the clean layout is scored by its median per-cell error after a rigid fit (rotation and reflection, no scale), divided by the root-mean-square radius of the true layout. A network that reads the layout improves as the layout becomes clean; one that ignores it scores the same at every level. Both models are probed on the same cells, those of the test graphs of LUNA-3D’s own runs (59,603 cells of Deep-STARmap block B and about 60,000 cells of mouse 3), LUNA-3D through its released test pipeline and noise schedule, and our model with the first of its three trainings. LUNA-3D is scored graph by graph in its own normalized coordinates, so the level of its error is not comparable with the per-cell errors of the main results; what the test reads is whether the error changes with the noise. Because the samples of LUNA-3D come out mirrored and rotated by 95–101° relative to its training graphs (on every graph of block B), it receives the noisy layout in both orientations.

LUNA-3D’s network ignores the layout of block B. From pure noise to a noise-to-signal ratio of 0.003, at which the pairwise distances of the layout it receives correlate with the true ones at 1.00, its prediction changes by less than 10%, the correlation of its pairwise distances with the true ones stays at 0.28–0.29, and its error stays at 0.80, in both orientations; a network that used only the direction of each cell from the centroid could reach a distance correlation of 0.84 from the clean layout. Its prediction is therefore set almost entirely by the expression of each cell, one best guess per cell, as expected from an architecture in which the noised positions reach the attention layers only as the direction of each cell from the centroid (Section 2). On the other tests we probed, it ignores the layout only in the orientation of its own samples: there its error stays at 0.45 at every level on mouse 3 and falls by 12% from pure noise to a nearly clean layout on fly embryo c, whereas in the orientation of its training graphs a nearly clean layout lowers it from 0.45 to 0.25 on mouse 3 and from 0.79 to 0.53 on fly embryo c. Our network reads the layout. On mouse 3 its error falls from 0.29 at pure noise to 0.11 at a noise-to-signal ratio of 0.13 and to 0.05 for a nearly clean layout; when the layout it receives is shuffled across cells, so that each cell carries its own expression but the noisy position of another cell, its error rises instead, to 0.55. Each step of our sampler therefore places the cells relative to the configuration drawn so far, which turns a chain of predictions into a sample of the whole specimen.

### S9 Reconstructed tissue blocks keep their local organization

#### Cell types that live together stay together

A reconstruction is often read through its neighborhoods: which cell types share a microenvironment and which are kept apart. In serially sectioned brains this can be measured only within a section, but in a continuous Deep-STARmap block every cell has its true neighbors in all three dimensions [8], so the neighborhoods of a reconstruction can be compared with the real ones from the scale of a few cells to that of a brain region. For every pair of cell types (*A, B*) and radius *r* we take the share of type *B* among all cells within *r* of a type-*A* cell and divide it by the share of *B* in the whole block; this co-occurrence ratio, of the form used by Squidpy [67], is 1 when the two types mix at random, above 1 when they live together and below 1 when they avoid each other. We compute it for the truth and for both reconstructions, with positions as produced after the rigid alignment used throughout, for the fine cell types with at least 200 cells (116 in block A, 119 in block B) and for *r* = 10, 20, 50, 100 and 200 *μ*m, and we score a reconstruction by the mean absolute difference between its log_2_ ratios and the true ones over all pairs of types, with ratios clipped to [1*/*16, 16].

Both reconstructions recover the broad organization of the block, which groups of cell types live together and which keep apart (Figure S2a–c); they differ in how sharply they reproduce it. Over all pairs of fine types at 50 *μ*m our error is 0.46 against 0.56 for LUNA-3D on block A (0.50 against 0.61 on block B), and it is smaller at every radius in both blocks, by 11–23% on block A and 9–29% on block B (Figure S2g, h). The gap is largest where the true tissue is most finely structured. The 28 subtypes of telencephalic projecting excitatory neurons of block A each live mostly among their own kind, which draws a bright diagonal in the true map (Figure S2d); our reconstruction keeps part of this diagonal, whereas in LUNA-3D’s map the subtypes blend into an almost uniform block (Figure S2f), and the error over these pairs is 0.59 against 0.96 (0.43 against 0.60 on block B). Compact structures show the same at the scale of a cell’s immediate neighbors: within 20 *μ*m, on average 86% of the neighbors of a dentate gyrus granule neuron and 88% of those of a choroid plexus cell belong to the same class in the true block A, against 52% and 77% in our reconstruction and 30% and 56% in LUNA-3D (block B: 86% and 89% in the truth, 55% and 82% for ours, 25% and 59% for LUNA-3D). This is the contraction of the main text (Figure 2) seen through neighborhoods. LUNA-3D packs the block into less than half of its true volume, so within any radius each cell collects neighbors that in the tissue lie farther away, and cell types that should be separate are mixed; our reconstruction keeps the block close to its real size and density, so a neighborhood of a given radius covers about the same tissue as in the real block.

**Figure S2:**
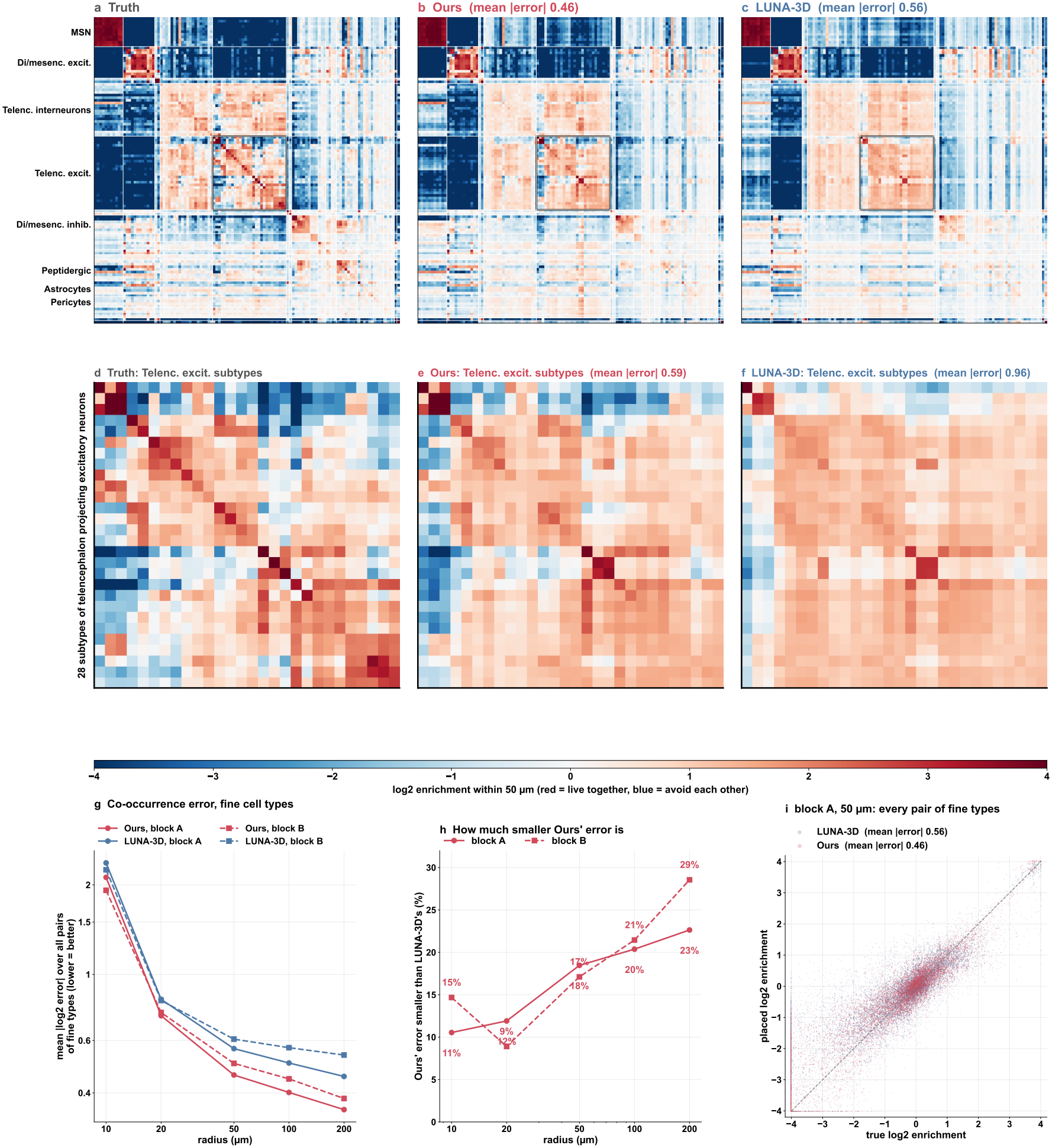
Which cell types live together in a continuous block (Deep-STARmap block A; 116 fine cell types with at least 200 cells). (a–c) Co-occurrence ratio within 50 *μ*m of every pair of fine types (log_2_; red, the types live together; blue, they avoid each other; white, random mixing) for the truth, our reconstruction and LUNA-3D. Types are grouped by main class (white lines; classes ordered by clustering the true class map, types within a class by clustering the truth), with the same order and color scale in all three panels; titles give the mean absolute error. (d–f) The 28 subtypes of telencephalic projecting excitatory neurons (black box in a–c); the bright diagonal marks subtypes that live among their own kind. Titles give the mean absolute error over these pairs. (g) Mean absolute log_2_ error over all pairs of fine types against radius, for both blocks. (h) How much smaller our error is than that of LUNA-3D, per radius and block. (i) Every pair of fine types at 50 *μ*m: placed against true log_2_ ratio; the diagonal is exact.

#### Marker genes stay in their cortical layers

The cerebral cortex is organized into layers of excitatory neurons, each marked by its own genes: *Cux2* in layers 2/3, *Rorb* in layer 4, *Fezf2* in layer 5 and *Foxp2* in layer 6 [64]. Every reconstructed cell keeps its own measured expression and only its position comes from the method, so the layers reappear only if the cells that carry each marker are placed in the right band, at the right depth below the cortical surface and with the right spacing; laminar markers are also the qualitative showcase of LUNA on two-dimensional sections of the motor cortex [45], and here we score them in three dimensions. For each of the 1,017 measured genes we build a smoothed map of its mean expression per cell on a 50 × 50 × 25 *μ*m grid (Gaussian kernel of one voxel), once from the cells at their true positions and once from the same cells at their reconstructed positions, taken as produced after the rigid alignment, and correlate the two maps (Spearman) over the voxels where the truth has at least 20 and each reconstruction at least five cells nearby (25,131 voxels in block B, 28,497 in block A). Placing the cells at random gives a correlation near zero (0.008 on block B, 0.015 on block A).

Our reconstruction rebuilds the layers (Figure S3a–c), and the map of every layer marker is closer to the truth than that of LUNA-3D in both blocks: on block B the correlation is 0.53, 0.60, 0.78 and 0.56 for *Cux2, Rorb, Fezf2* and *Foxp2*, against 0.29, 0.40, 0.51 and 0.32 for LUNA-3D, and on block A it is 0.55, 0.58, 0.82 and 0.59 against 0.39, 0.48, 0.72 and 0.46. The view of the cortex shows why (Figure S3d–f). In the measured cortex the four markers form bands parallel to the surface, with *Cux2* and *Rorb* cells outermost, *Fezf2* below and *Foxp2* deepest. Our reconstruction places the same cells in the same banded order at close to the real density, about 30,000 cells in a 1.4 mm window against 25,000 in the truth, whereas LUNA-3D packs about 49,000 cells into the same window: its bands are present but pressed together into a thin, crowded cortex, the contraction of the main text (Figure 2) at the scale of the layers. Beyond the four markers, our map is closer to the true one for 619 of the 1,017 genes in block B (mean correlation 0.30 against 0.26) and for 844 in block A (0.35 against 0.29; Figure S3g).

**Figure S3:**
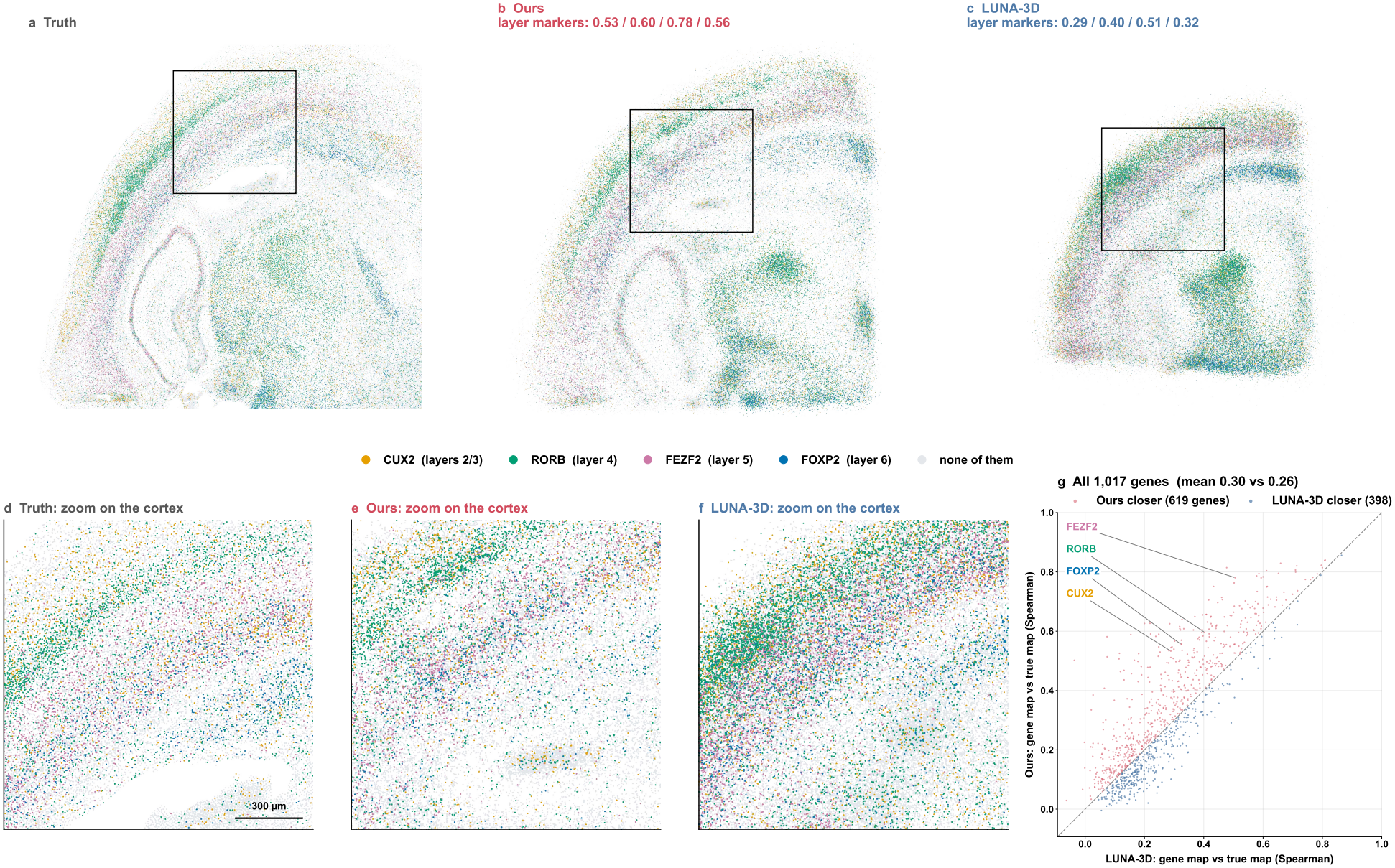
Cortical layers rebuilt from the marker genes of placed cells (Deep-STARmap block B). (a–c) Face view of the whole block, measured (a) and reconstructed by our model (b) and LUNA-3D (c). Every cell is colored by the layer marker it expresses most strongly (*Cux2*, layers 2/3; *Rorb*, layer 4; *Fezf2*, layer 5; *Foxp2*, layer 6; each gene scaled to its 99th percentile); cells expressing none of them are grey. A cell’s color comes from its own measured expression, so only its position differs between panels. Titles of b and c give the Spearman correlation of each marker’s map with its true map; black boxes mark the windows of d–f. (d–f) The cortex up close: a 1.4 mm window centered on where each layout placed the cells of the same true cortical region, at true scale. (g) All 1,017 measured genes: correlation of each gene’s map with its true map, ours against LUNA-3D; points above the diagonal favor our reconstruction, and the four layer markers are labeled.

#### Virtual sections through the reconstruction match real sections

Much of brain anatomy is done on sections: a thin slice is cut and the cells it contains are counted. A three-dimensional reconstruction can be cut the same way at any position, and such a virtual section is useful only if it contains the cells that a real section cut at the same place would contain; in a continuous block the true content of every section is known. We cut each block into parallel sections 100 *μ*m thick along each of its two long axes, at fixed true positions that are the same for the truth and both reconstructions (positions as produced, no rescaling), and keep the sections that hold at least 200 true cells (43 and 42 along the two axes of block B, 42 and 42 in block A). The error of a reconstructed section is the Jensen–Shannon divergence (base 2) between its composition over the 19 main cell classes and that of the real section at the same place, 0 for the same composition and 1 for disjoint ones; a reconstructed section with fewer than 20 cells counts as empty, with error 1.

Sections cut through our reconstruction contain the right cells across almost the whole block, while those of LUNA-3D go wrong progressively towards its edges (Figure S4). Averaged over the sections along both axes, the error is 0.05 against 0.25 in block B and 0.07 against 0.16 in block A. Within about 1 mm of the center of the block the two reconstructions are comparable (0.06 against 0.06 for the sections of block B cut along *x*), but farther out the error of LUNA-3D rises steadily with distance from the center (Figure S4d, e): at −1.8 mm in block B it is 0.22 against 0.01 for ours, and the sections near the true edges of the block are empty, 9 of 43 along *x* and 7 of 42 along *y* in block B (5 and 4 of 42 in block A). Our error stays low across most of the block. In block B it is highest in a short stretch near the center (0.22–0.25 around −0.3 mm along *x*) and in the last 200 *μ*m at one end, where our reconstruction places too few cells and the outermost section is empty; in block A it rises in the outermost 0.5 mm (0.35 and 0.72 beyond 2 mm from the center, where we place 7% and 2% of the true cells), but there the sections of LUNA-3D are already empty, and its error reaches 0.45 half a millimeter earlier, at 1.75–2 mm, where ours is 0.05. The pattern follows from the contraction of the main text (Figure 2). LUNA-3D shrinks the block towards its center to 71–79% of the true extent along each axis, and under such a contraction the displacement of a cell grows with its distance from the center: sections near the center keep their cells, sections farther out collect cells that belong farther out, and at the edges no cells remain. Our reconstruction keeps 89–99% of the true extent along each axis, so its sections stay correct until the last few hundred micrometers of the block.

**Figure S4:**
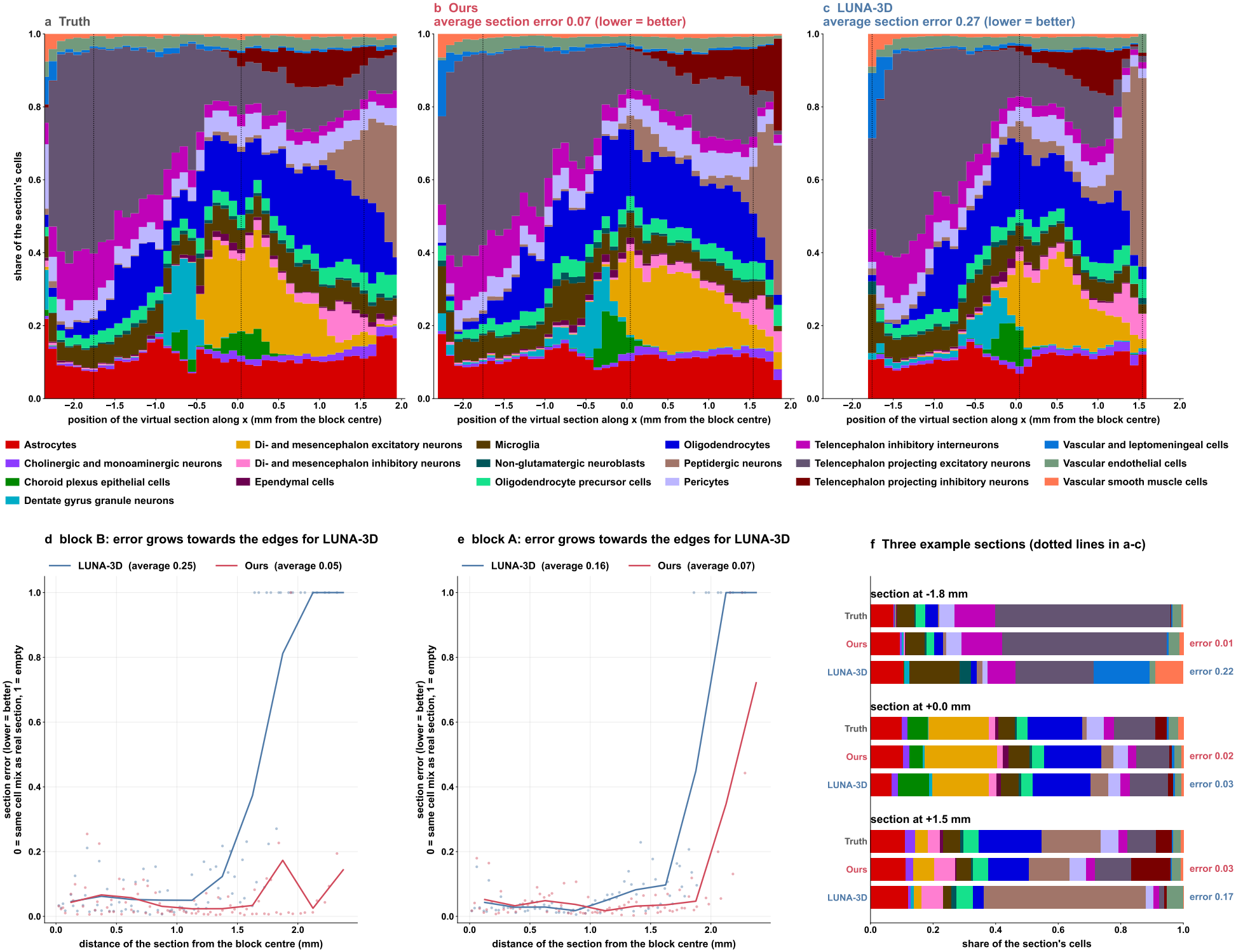
Virtual sections through Deep-STARmap block B (100 *μ*m thick, cut along *x*). (a–c) Cell-type composition of every section along the cut axis for the measured block (a), our reconstruction (b) and LUNA-3D (c); each color is one main cell class, and white means that no cells were placed at that position. Titles give the average section error (lower is better); dotted lines mark the sections shown in f. (d, e) Section error against distance from the center of the block, for block B (d) and block A (e), with sections cut along *x* and along *y*; dots are single sections and lines are averages in 0.25 mm bins. An error of 0 means the same cell mix as the real section, and 1 means an empty section. (f) Three example sections at −1.8, 0.0 and +1.5 mm: their cell mix in the measured block, in our reconstruction and in LUNA-3D, with the error of each reconstruction. At +1.5 mm LUNA-3D places only 116 cells where the real section holds 5,088.

### S10 Reconstructed embryos keep the internal organization of their organs

#### ORBIS reconstructs the expression gradients within organs

The cells of one organ are not alike: many genes rise or fall along an organ, and the fly midgut, for example, expresses different digestive proteases of the Jonah family in different stretches along its length [68, 62]. Recovering such spatial expression patterns from dissociated cells is a long-standing use of spatial reconstruction, first demonstrated on the early fly embryo [69, 31]. In a reconstruction every cell keeps its own measured expression and only its position comes from the method, so a gradient inside an organ survives only if each cell is placed at its right position *within* the organ; placing it in the right organ is not enough. We test this on held-out embryo c (Figure S5), scoring each gradient by the rank correlation *r* between a gene’s expression and the cells’ position along one body axis inside the organ. Two genes among the five strongest true midgut gradients, selected on the truth alone, illustrate the effect. The Jonah protease *Jon25Biii* falls from head to tail inside the true midgut (*r* = −0.40); our reconstruction rebuilds the same gradient (*r* = −0.40), whereas LUNA-3D keeps less than half of it (*r* = −0.17) and places part of the midgut in front of the true organ (Figure S5g). The acyl-CoA binding protein *Acbp5* gives *r* = −0.40 in the truth, −0.32 for our reconstruction and −0.19 for LUNA-3D. Over every gradient in every organ of the embryo (307 gradients, each a gene, an organ and one of the three body axes; organs with at least 300 cells, genes expressed in at least 10% of the organ’s cells, true |*r*| *>* 0.2), our reconstruction keeps at least half of the true strength, in the right direction, for 79% of the gradients against 64% for LUNA-3D, and its curve lies above that of LUNA-3D at every threshold (Figure S5i). The reason is the one behind the order of gut cells (Figure 3): a single best guess per cell pulls the cells of an organ toward its middle and flattens the gradients they carry, whereas sampling keeps the cells spread along the organ, and the gradients with them.

**Figure S5:**
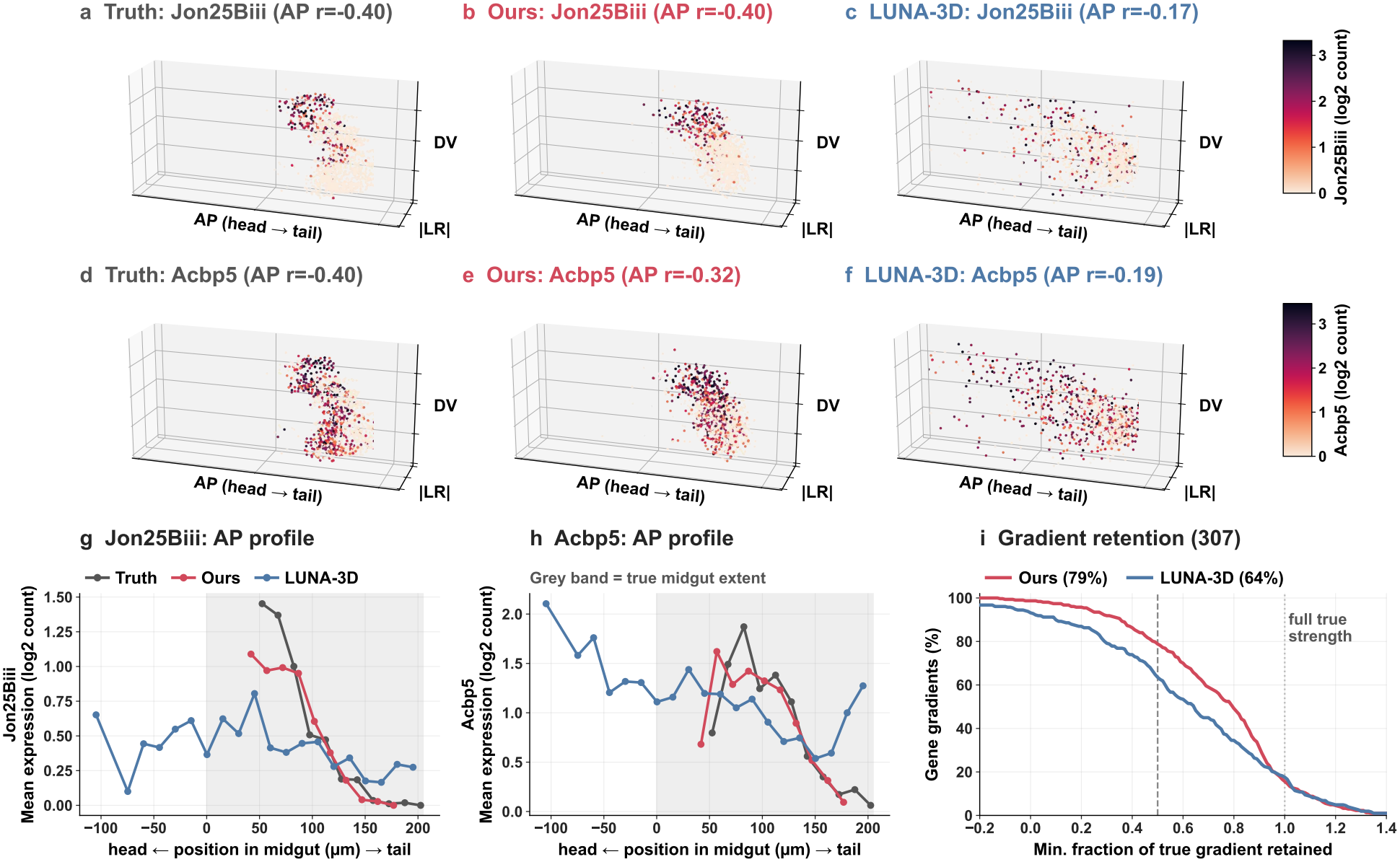
Gene-expression gradients inside organs of a held-out *Drosophila* embryo (embryo c). (a–f) The midgut in 3-D for the truth, our model and LUNA-3D, every cell colored by its measured expression of the Jonah protease *Jon25Biii* (a–c) and of the acyl-CoA binding protein *Acbp5* (d–f); *r* is the rank correlation between the gene and head-to-tail position inside the midgut (negative: highest at the head end). (g, h) Mean expression of the same genes along the midgut, head to tail; the grey band marks the extent of the true midgut, so a curve outside it means cells placed outside the true organ. (i) Every gradient inside every organ (307 gradients): the share of each true gradient’s strength that survives the reconstruction (1, full strength; 0, flattened; below 0, reversed), as the fraction of gradients above each threshold; higher is better.

#### Cells are placed in their own body segment

Along the anterior–posterior axis of bilaterian animals, Hox genes give each body segment its identity, and they are expressed in overlapping domains whose order along the body matches the order of the genes on the chromosome [70, 71]. In the *Drosophila* embryonic nervous system, of the five Hox genes we use, *Sex combs reduced* (*Scr*) is expressed most anteriorly, *Antennapedia* (*Antp*) in the thorax, and *Ultrabithorax* (*Ubx*), *abdominal-A* (*abd-A*) and *Abdominal-B* (*Abd-B*) in successively more posterior abdominal segments, domains that the Flysta3D atlas resolved in 3-D in the embryos we use [50]. Because every cell keeps its own measured expression and only its position comes from the reconstruction, these domains reappear in their true places only if each brain and nerve-cord cell is placed in the right segment; this tests positioning at the scale of a single segment, finer than the organ scale of the gut (Figure 3). We use the 4,729 brain and ventral nerve cord cells of held-out embryo c, 439 of which express at least one of five Hox genes (*Scr* 20, *Antp* 114, *Ubx* 99, *abd-A* 172 and *Abd-B* 64 cells). The gene list was fixed before any result was computed: it is the Hox genes of the nervous system, without *labial* and *Deformed*, which are absent from the measured gene panel, and *proboscipedia*, which is expressed in too few nerve-cord cells to profile. Positions are read along the anterior–posterior axis after the alignment used for the main table, with left and right folded onto one side, because no method can read a cell’s side from its expression, and with no rescaling. For each gene we sum expression in 10 *μ*m bins along the axis, smooth it over seven bins (about 70 *μ*m, or two segments, because Hox counts per cell are sparse) and mark the center of mass of each domain. We then compare the share of each gene’s expression found in every bin with the truth, summarizing the difference by the absolute difference summed over bins and averaged over genes, and we count the Hox-expressing cells placed within one segment of their true position, taking a segment as 25 *μ*m from the roughly eleven segments that span about 250 *μ*m of the late-embryonic nerve cord; this length is an estimate rather than a measurement on this embryo, and the ranking of the two methods is the same at 30 and 50 *μ*m.

Both reconstructions keep the five domains in their colinear order (Figure S6a–c), but ours places them more accurately. It puts 44% of the Hox-expressing cells within one segment of their true position, against 31% for LUNA-3D, and its domain maps deviate less from the truth (0.34 against 0.46; Figure S6d–f). The largest errors of LUNA-3D concern the thoracic and abdominal genes *Antp, Ubx* and *abd-A*, whose domains are depleted at their true positions and displaced toward the tail. The advantage extends to positional order inside organs in general: for every organ with at least 300 cells, the rank correlation between true and placed anterior–posterior position is higher for our reconstruction in all 21 organs of embryo c and in 19 of 22 organs of embryo d (mean 0.52 against 0.35 and 0.47 against 0.36; Figure S6g). The reason is the mechanism behind the order of gut cells and the gradients inside organs. Neighbouring segments share most of their expression and adjacent Hox domains overlap, so the expression of a cell is compatible with a range of positions that spans more than one segment. A single best guess per cell reports the middle of that range, which blurs the boundaries between domains and moves cells away from the segment they come from; our model instead samples one position from the range for every cell, so the cells of a domain keep their spread and the domain keeps its boundaries.

**Figure S6:**
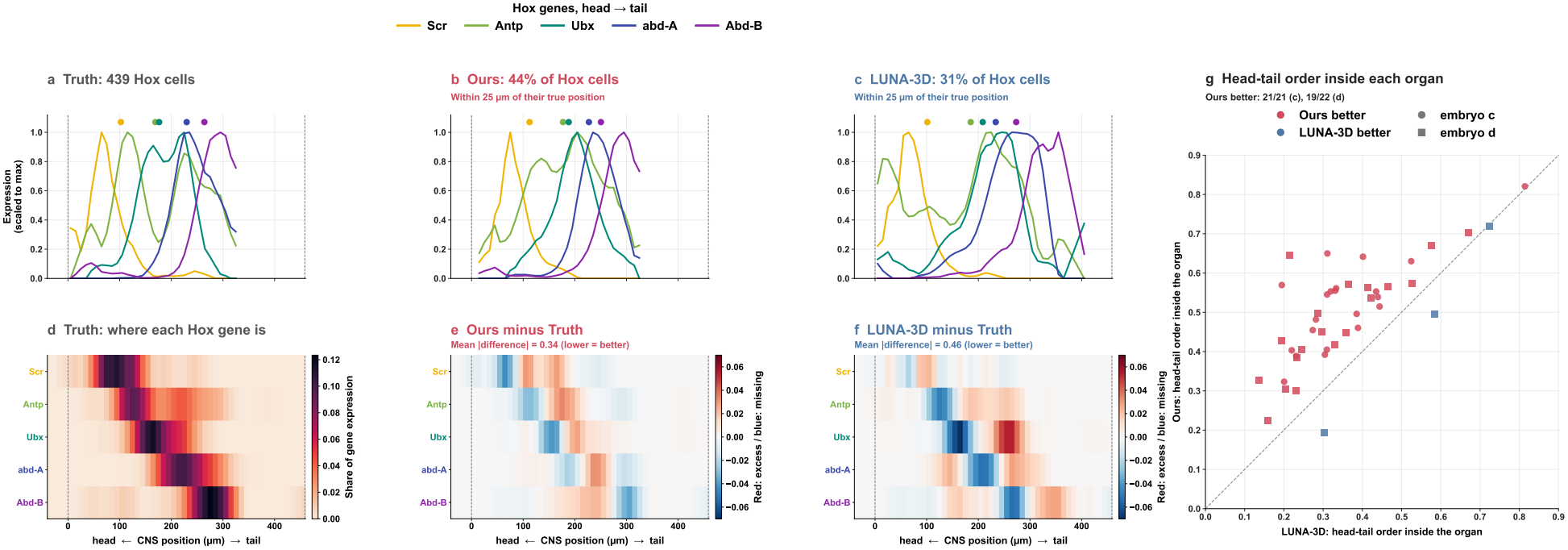
The Hox code along the central nervous system of a held-out *Drosophila* embryo (embryo c; 4,729 brain and ventral nerve cord cells, 439 of them expressing a Hox gene). (a–c) Expression of each Hox gene along the anterior–posterior axis of the nervous system for the truth, our reconstruction and LUNA-3D; every profile is smoothed over about two segments and scaled to its own maximum, dots mark each gene’s center of mass, and dashed lines the ends of the true nervous system. Titles of (b, c) give the share of Hox-expressing cells placed within one segment (25 *μ*m) of their true position. (d) Domain map of the truth: the share of each gene’s expression found at each position. (e, f) Our reconstruction and LUNA-3D minus the truth; red marks expression placed where the gene is not, blue expression missing from where it is; titles give the mean absolute difference (lower is better). (g) Every organ with at least 300 cells in both held-out embryos (circles, embryo c; squares, embryo d): rank correlation between true and placed anterior–posterior position of the organ’s cells for LUNA-3D (x) and our reconstruction (y); points above the diagonal are organs whose internal order our reconstruction keeps better.

**Figure S7:**
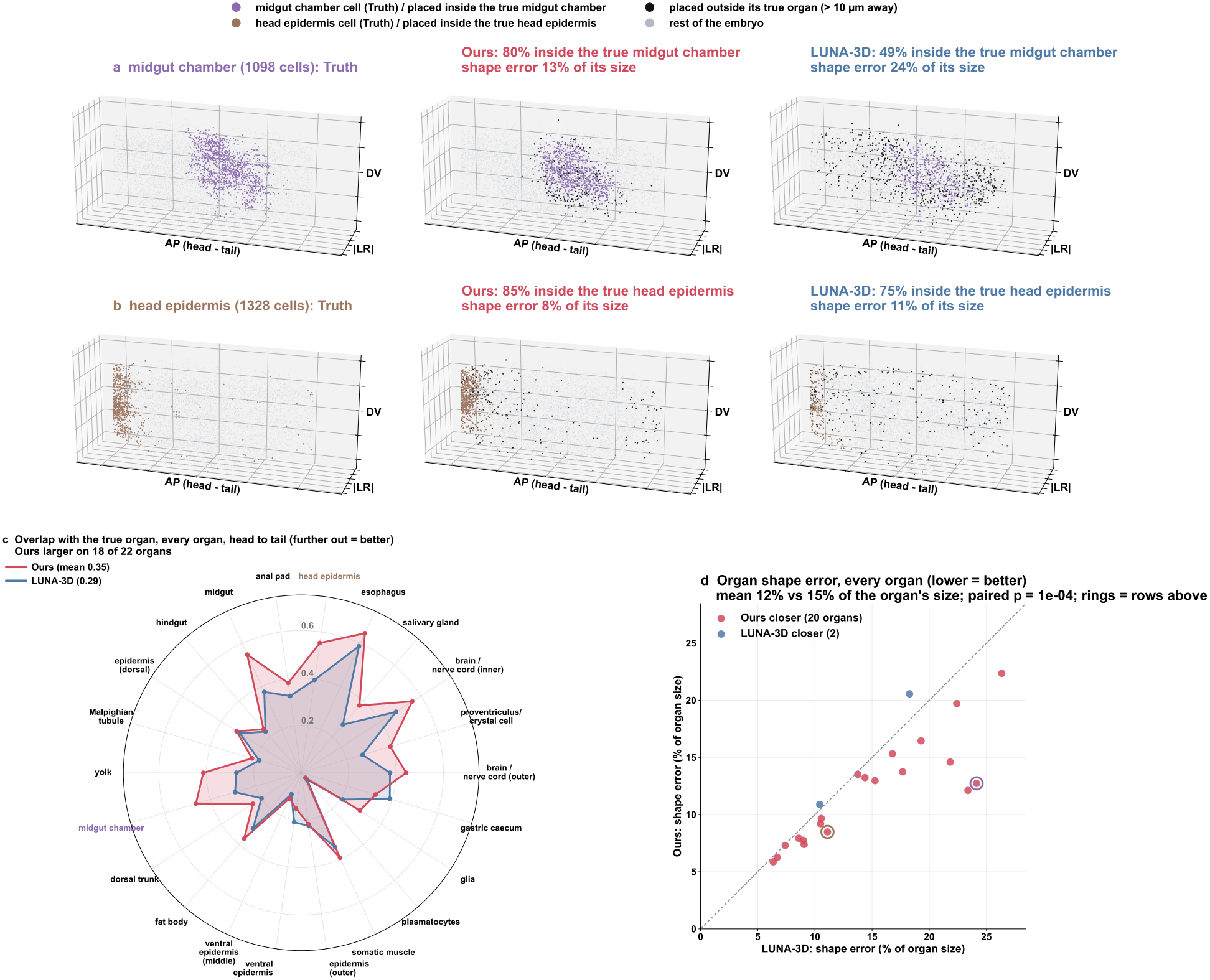
Organs of a held-out *Drosophila* embryo rebuilt in 3-D (embryo c; 22 annotated organs with at least 200 cells). (a, b) Two organs pulled out of the whole embryo, the midgut chamber (a; 1,098 cells) and the head epidermis (b; 1,328 cells), for the truth, our reconstruction and LUNA-3D. In the reconstructions, a cell is drawn in the organ’s color when it is placed inside its true organ (within 10 *μ*m of one of the organ’s true cells) and in black when it is placed outside it; grey cells are the rest of the embryo. Titles give the share of the organ’s cells placed inside it and its shape error. (c) Overlap of every organ with its true volume (Dice coefficient of the 10 *μ*m cubes each organ occupies; 1 means the same volume), organs ordered from head to tail around the circle; further out is better. (d) Shape error of every organ (mean distance between placed and true cells of the organ in both directions, as a percentage of the organ’s size) for LUNA-3D (x) and our reconstruction (y); points below the diagonal are organs our reconstruction rebuilds more closely, and rings mark the two organs of (a, b).

#### Organs keep their volume and shape

A whole organism is not a slab of tissue: its organs are solid 3-D shapes nested inside one another, from compact masses in the interior of the body to thin layers on its surface, and a reconstruction of a whole embryo is useful for anatomy only if each organ comes out with its true volume and shape [19]. We test this organ by organ on held-out embryo c, whose annotation lists 22 organs with at least 200 cells (25,677 of its 25,755 cells); both methods were trained on the same eleven embryos and are scored on the same held-out embryo, after the alignment used for the main table, with left and right folded onto one side because no method can read a cell’s side from its expression, and with positions taken as produced, without rescaling. For every organ we pull its cells out of each reconstruction and compare them with the same cells in the true embryo through three readouts. A cell is placed *inside* its organ when it lies within 10 *μ*m of one of the organ’s true cells. The *overlap* of an organ is the Dice coefficient between the sets of 10 *μ*m cubes occupied by its true and by its placed cells, so 1 means the same volume. Its *shape error* is the mean distance from each placed cell to the nearest true cell of the organ and from each true cell to the nearest placed cell, averaged over both directions and divided by the organ’s size, the root-mean-square distance of its true cells from their center; this normalization makes small and large organs comparable, since an error of 5 *μ*m is large for a small organ and negligible for the epidermis. The 10 *μ*m radius and cube size sit just above the largest gap between neighbouring true cells (4.6 *μ*m within a section and 7.4 *μ*m between sections), so that every true organ forms one connected volume at this scale.

Two organs of contrasting form illustrate the comparison (Figure S7a, b): the midgut chamber, a compartment of the gut tube deep in the interior of the embryo and the same part whose internal order our reconstruction keeps in the main text (Figure 3), and the head epidermis, a thin layer at the anterior surface. Our reconstruction places 80% of the midgut-chamber cells inside the true midgut chamber, against 49% for LUNA-3D, and 85% of the head-epidermis cells inside the true head epidermis, against 75%, with a smaller shape error for both organs (13% against 24% of the organ’s size for the midgut chamber and 8% against 11% for the head epidermis) and a larger overlap (0.46 against 0.29 and 0.55 against 0.40). Over all 22 organs, our reconstruction has the smaller shape error for 20 organs (mean 12.2% against 14.7% of the organ’s size; paired Wilcoxon signed-rank test, *p* = 9.9 × 10^*−*5^) and the larger overlap for 18 (mean 0.35 against 0.29; *p* = 5.9 × 10^*−*4^), with the gains spread over organs from the head to the tail of the embryo (Figure S7c, d). The difference again follows from sampling rather than predicting a single best guess per cell. When the expression of a cell is compatible with several positions, their average can fall outside the organ altogether, and for an organ that is a curved layer, such as the epidermis, the average of positions on the layer lies beneath it, so best guesses pull surface organs inward and off their true volume; our model instead samples one compatible position for every cell, which lies in the organ, so the organ keeps its volume, its surface and its shape.

### S11 Reconstructed brains keep their anatomy, from regions to layers and subfields

#### Cortical neurons land in their own layers across a whole brain

The neocortex is a curved sheet a few hundred micrometres thick, divided along its depth into six layers, and each subclass of its excitatory neurons occupies characteristic layers: intratelencephalic (IT) neurons span layers 2/3 to 6, extratelencephalic (ET) and near-projecting (NP) neurons lie in layer 5, corticothalamic (CT) neurons in layer 6, and layer 6b neurons at the boundary with the white matter [72, 73, 16]. Laminar position is therefore a fine-scale test of a whole-brain reconstruction. A reconstruction that is slightly too small, or that pulls cells toward the average position of their type, pushes deep-layer neurons out of the cortex and superficial neurons into deeper layers, errors that per-cell distances average away. We test mouse 1 (2.85 million cells in 147 coronal sections; both models trained on mice 2–4 and given expression only) and score the eight excitatory subclasses of the isocortex (120,891 neurons, from 463 for L6b to 47,139 for L2/3 IT). Layer labels come from the registration to the Allen Common Coordinate Framework released with the atlas [15] and are used only for scoring: the cells are split at random into an atlas half, which keeps its measured positions and labels, and a query half, and each query neuron takes the label of the atlas cell nearest its placed position. For each subclass we report the share of its cortical neurons placed inside the isocortex (*kept in cortex*) and the overlap ∑_*ℓ*_ min(*p*_*ℓ*_, *t*_*ℓ*_) between the distribution *p* of the layers where it is placed and its true distribution *t* (*layer match*). The measured positions passed through the same readout give its ceiling, 98.0% kept and 97.2% match, below 100% because a neuron at a layer boundary can have its nearest atlas cell on the other side. Our model is scored on four samples of one training, each after reseating.

Our model keeps cortical neurons in the cortex and in their own layers for every excitatory subclass (Figure S8). Averaged over the eight subclasses, it keeps 78.2% of them in the cortex against 64.5% for LUNA-3D, with a layer match of 58.8% against 44.7% (standard deviation over our four samples 0.3 and 0.1 points), and each of its samples is above LUNA-3D for every subclass on both measures. The gain is largest at the inner boundary of the cortex. LUNA-3D keeps half of the corticothalamic neurons in the cortex (L6 CT 49.5%), our model 69.3%, and the same holds for L6 IT (73.3 versus 55.7%), L5 ET (82.0 versus 69.0%) and L5 NP (84.6 versus 64.9%); for L6b, the thinnest layer, our model nearly doubles both measures (kept 39.5 versus 21.4%, ceiling 87.0%; match 36.0 versus 21.4%). Deep neurons that leave the cortex go, for both models, to the structures beneath it: the white matter, the hippocampus and the striatum. The superficial layer fails differently. L2/3 IT neurons stay in the cortex for both models (92.5 versus 91.6%), but LUNA-3D places 47% of them in layers 5 and 6a, against 27% for our model, and their layer match is 51.2 versus 35.6%. Both failures are those of a contracted cortical sheet: LUNA-3D reconstructs mouse 1 at 79, 89 and 93% of its true extent along the left–right, dorsal–ventral and anterior–posterior axes, our model at 92, 100 and 97%, and LUNA-3D squeezes 12.6% of the cell types against 2.5%. The one-call best guess of our own network, read out the same way, lies between the two (71.6% kept, 51.3% match): about half of the gain over LUNA-3D comes from the trained model and half from sampling positions rather than predicting their average (Section S7). Reseating changes our numbers by at most 1.3 points. On mouse 2 the ordering holds (kept 79.3 versus 69.8%, match 58.0 versus 50.4%); on the sagittally cut mouse 3 and on the Allen brain the two models are close (kept 86.8 versus 84.0% and 78.7 versus 80.7%, match 52.2 versus 55.1% and 55.3 versus 55.9%).

**Figure S8:**
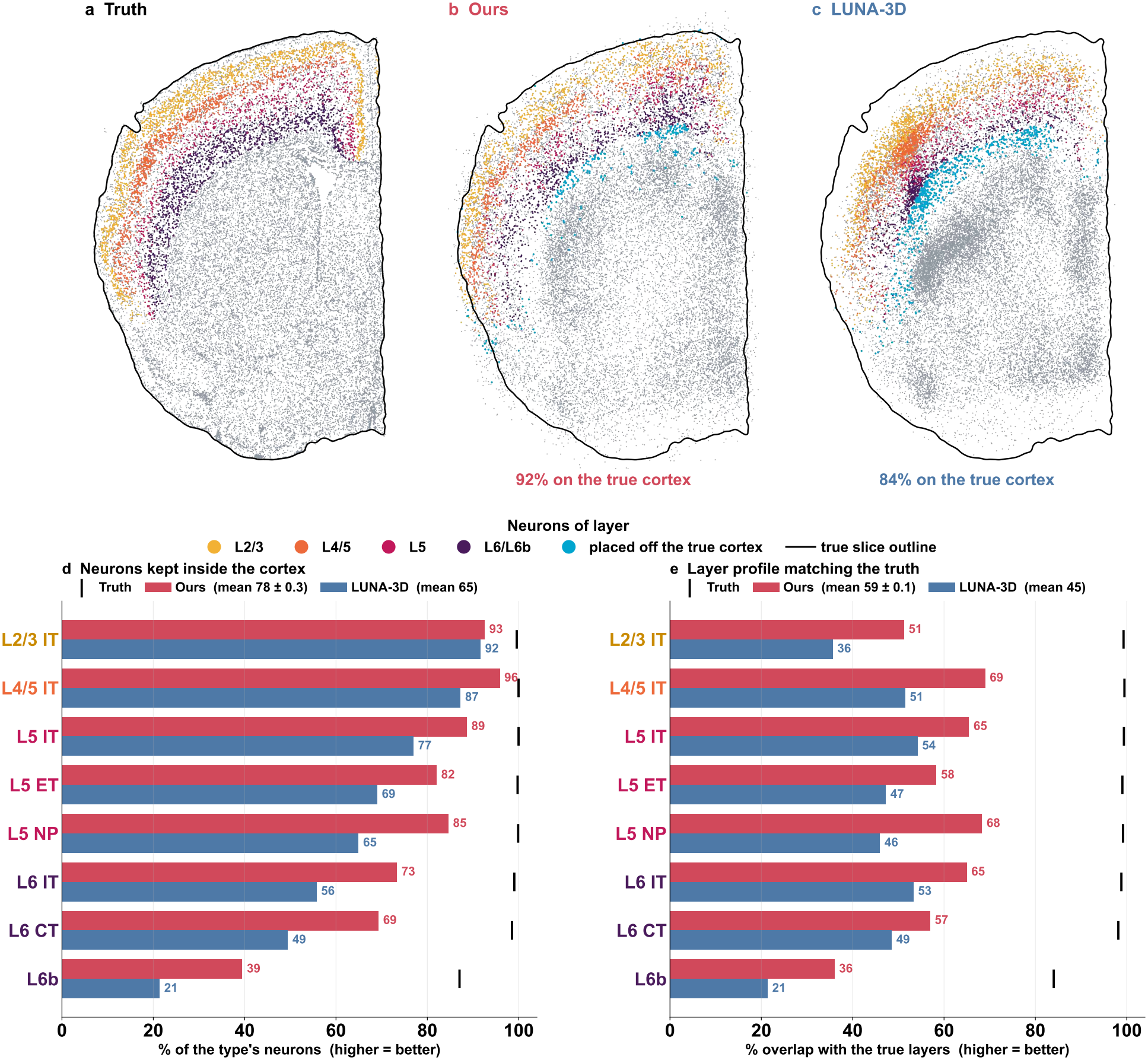
Cortical layers of a held-out whole brain (mouse 1; models trained on mice 2–4, given expression only). (a–c) Virtual coronal slab, every cell whose measured (a) or placed (b, c) position lies within ±40 *μ*m of the anterior–posterior position of section 060, for the truth, our model (one sample) and LUNA-3D; excitatory neurons colored by layer group, other cells grey. Black line, outline of the true slab, identical in every panel; each placed slab is translated as a whole (no rotation or scaling) to best fill it. Neurons drawn cyan lie off the true cortex after that translation; the percentage gives the share on the true cortex. (d) Share of each subclass’s cortical neurons placed inside the isocortex. (e) Overlap between the layers where each subclass is placed and its true layers. In (d, e), bars give our model (mean of four samples) and LUNA-3D, and the black tick the measured positions read out the same way.

#### A brain from another laboratory is reconstructed with its regions in place

Every other brain test shares its laboratory, platform and gene panel with the training brains, so a model could learn the atlas rather than the anatomy. The Allen brain [16] removes this: it was profiled by another laboratory on another platform with another gene panel, and both models, trained on mice 1–4 only, receive the 477 shared genes integrated with Harmony [66] and place all 3.74 million cells as if dissociated (Supplementary Section S2). We ask whether the major structures of the brain land where they are. For each of the 12 grey-matter divisions of the Allen Common Coordinate Framework [65] (isocortex, olfactory areas, hippocampal formation, cortical subplate, striatum, pallidum, thalamus, hypothalamus, midbrain, pons, medulla and cerebellum), labelled by the registration released with the atlas and used only for scoring, we measure the distance between the centre of its cells under placed and under measured positions, with the two hemispheres folded onto one as in every score of this brain; a layout that puts every division at the centre of the brain would be 2,706 *μ*m off on average. Our model places the division centres 399 *μ*m from where they belong on average, against 489 *μ*m for LUNA-3D (standard deviation over our four samples below 1 *μ*m), and is closer for 8 of the 12 divisions (Figure S9d). The largest gains are in the deep structures of the forebrain and in the cerebellum: the striatum (97 versus 185 *μ*m), the thalamus (109 versus 281 *μ*m), the isocortex (220 versus 410 *μ*m), the cerebellum (345 versus 692 *μ*m), the hippocampal formation (482 versus 672 *μ*m) and the pallidum (516 versus 824 *μ*m); LUNA-3D is closer for the medulla, the pons, the olfactory areas and the cortical subplate. The brain as a whole keeps its size, at 96, 93 and 96% of its true extent along the left–right, dorsal–ventral and anterior–posterior axes against 82, 87 and 105% for LUNA-3D, and its cell types keep their shape (shape error 212 versus 449 *μ*m; 1.4 versus 24.5% of the cell types squeezed). A virtual sagittal slab crosses all 53 coronal sections (Figure S9a–c): the measured brain appears as stripes because it was sectioned with gaps, whereas the reconstructions fill the volume between sections.

**Figure S9:**
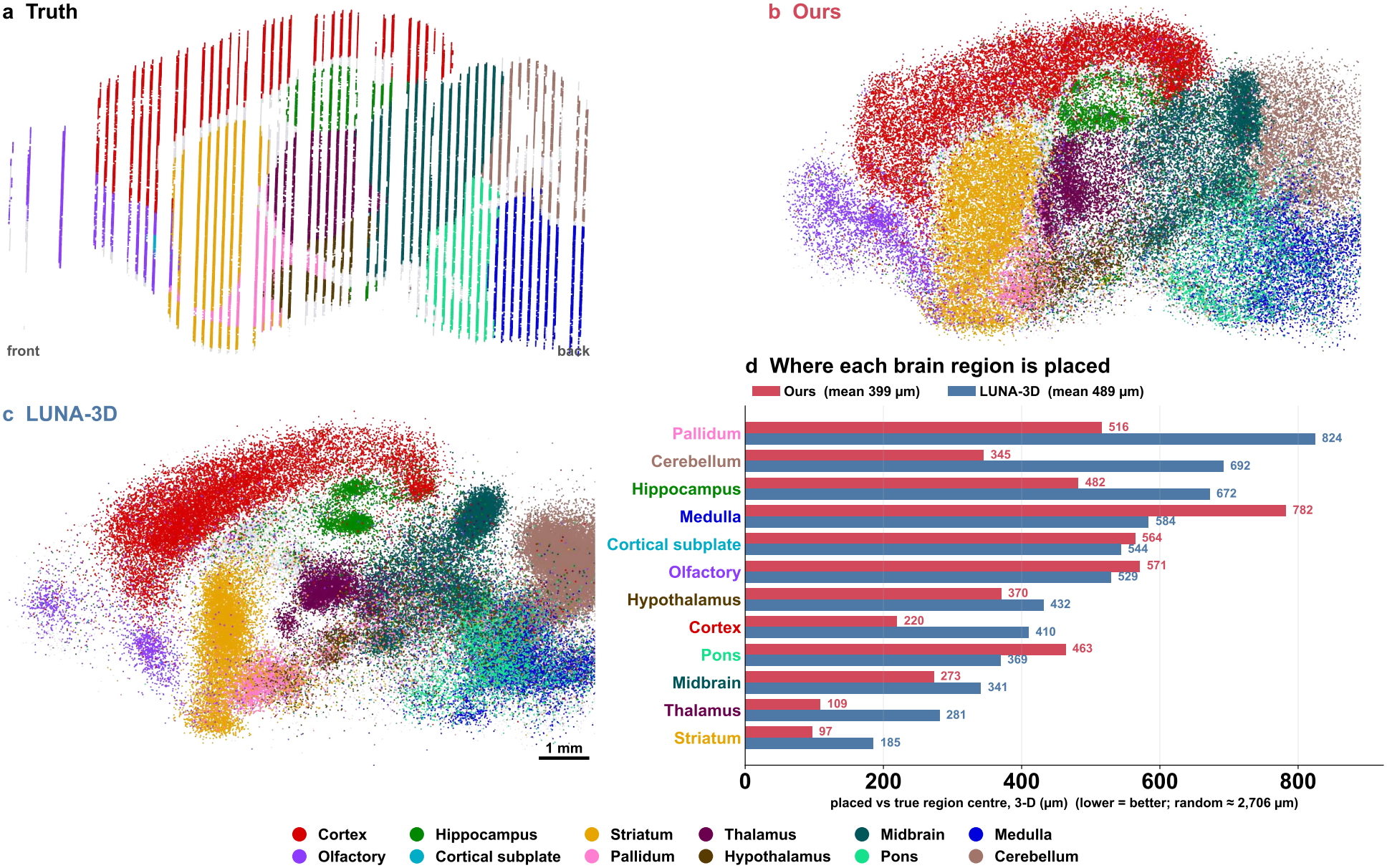
Brain regions in a brain from another laboratory (Allen MERFISH brain 638850, 3.74 million cells in 53 coronal sections; models trained on mice 1–4 and given the 477 shared genes only). (a–c) Virtual sagittal slab 1.5 mm from the midline (± 40 *μ*m) for the truth, our model (one sample) and LUNA-3D; cells colored by brain division, other cells grey; hemispheres folded onto one. (d) Distance between the centre of each division’s cells under placed and under measured positions, for our model (mean of four samples) and LUNA-3D, ordered by LUNA-3D’s distance; a layout with every division at the brain centre gives 2,706 *μ*m on average.

#### Hippocampal neurons land in their own subfields

The hippocampus is the most tightly folded structure a whole-brain reconstruction has to keep. Its principal neurons form thin, curved sheets that interlock in a C shape: the granule-cell layer of the dentate gyrus and the pyramidal layers of CA3 and CA1, which continue into the subiculum. The subfields lie a few hundred micrometres apart and are the successive stages of the main excitatory pathway of the hippocampus, from the dentate gyrus through CA3 and CA1 to the subiculum [74], so a neuron placed in the wrong subfield is placed in the wrong stage of that pathway. Where the cortical test asks for the depth of a neuron within one smooth sheet, this test asks whether a reconstruction keeps folded, adjacent sheets apart. We score the four excitatory types of the hippocampal formation in the atlas taxonomy [16], dentate gyrus granule cells, CA3 pyramidal cells, CA1–prosubiculum neurons and subiculum–prosubiculum neurons (28,152 neurons in mouse 1), with the readout of the cortical test: each placed neuron takes the structure of the atlas cell nearest its placed position, from the registration to the Allen Common Coordinate Framework [65] used only for scoring. For each type we report the share placed inside the hippocampal formation, the share placed in its own subfield (for the two types the taxonomy assigns to two structures, either counts) and the overlap between the distribution of subfields where the type is placed and its true distribution (*subfield match*); the measured positions reach a match of 99.3%.

On mouse 1, our model places 79.7% of these neurons inside the hippocampal formation against 63.3% for LUNA-3D, with a subfield match of 65.3% against 35.2% (standard deviation over our four samples 0.3 points), and puts about twice as many neurons of every type into its own subfield (Figure S10d): 61.9 against 33.0% of the dentate gyrus granule cells, 54.8 against 23.2% of the CA3 and 71.2 against 36.3% of the CA1 pyramidal cells, and 18.8 against 5.1% of the subicular neurons (measured positions 96.6, 83.9, 95.9 and 64.7%). On the brain from another laboratory, our model is ahead for every type (subfield match 82.8 against 67.8%, inside the hippocampal formation 90.4 against 77.7%; own subfield 78.1 against 66.8% for CA1, 69.6 against 47.4% for CA3, 61.5 against 56.5% for the dentate gyrus and 35.1 against 20.8% for the subiculum). On mice 2 and 3, LUNA-3D has the higher subfield match (65.7 against 60.2% and 53.4 against 52.4%; Figure S10e), and on mouse 2 it also keeps slightly more neurons inside the hippocampal formation (84.3 against 81.4%; on mouse 3, 70.1 against 73.2% for ours). The mechanism is the one behind the cortical layers. A neuron whose expression fits anywhere along a curved sheet has a distribution of plausible positions shaped like the sheet; its average falls inside the curve, in the gap between sheets, which for adjacent subfields is another subfield or tissue outside the hippocampus, whereas a sample lands on the sheet. A whole-brain contraction, such as LUNA-3D’s reconstruction of mouse 1 at 79–93% of the true extent, closes the fold further.

**Figure S10:**
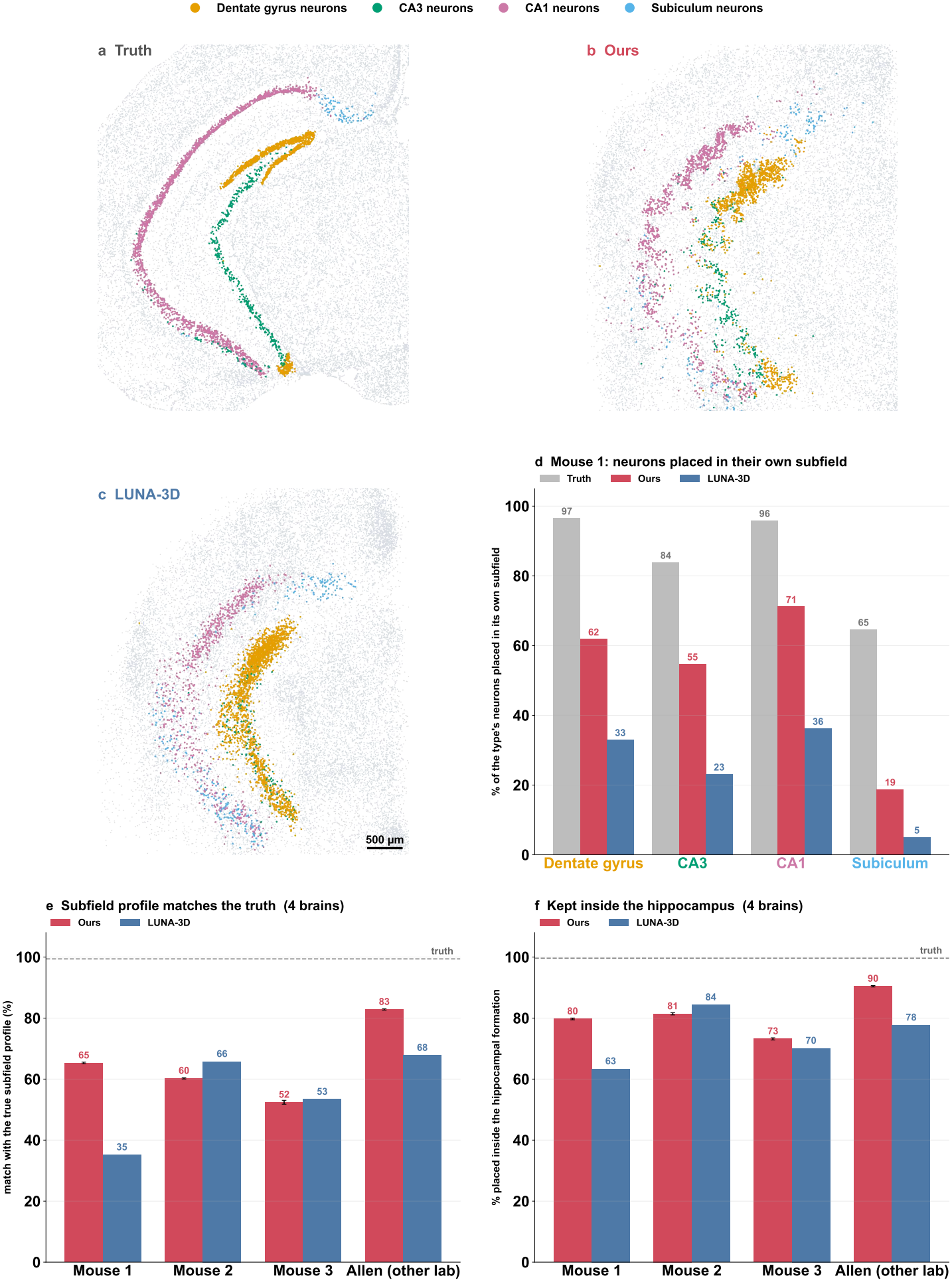
Hippocampal neurons placed into their own subfield. (a–c) Virtual coronal slice through the hippocampus of mouse 1 (± 40 *μ*m around the anterior–posterior centre of its dentate gyrus granule cells) for the truth, our model (one sample) and LUNA-3D; dentate gyrus granule cells, CA3, CA1 and subicular neurons colored, other cells grey. (d) Mouse 1: share of each type’s neurons placed in its own subfield, for the measured positions, our model (mean of four samples) and LUNA-3D. (e) Overlap between the subfields where each type is placed and its true subfields, and (f) share of the neurons placed inside the hippocampal formation, both averaged over the four types, for the four test brains.

#### Cell types keep their place in an independent, morphology-based atlas

Every anatomical readout above uses the Allen Common Coordinate Framework [65], the atlas the brains were registered to. To test whether our advantage depends on that atlas and whether it survives at a finer scale, we also score the reconstructions against CCF-ME [75], a whole-brain parcellation built from a different modality, the dendritic morphology of 101,136 reconstructed neurons from 111 brains, which divides the brain into 1,057 regions against 582 in the Allen framework; neither expression nor any of our data was used to build it, and its authors compared it with the MERFISH atlas of mouse 1 itself. Each measured cell receives its CCF-ME subregion from its official atlas coordinate (cells without one, 1–14% per brain, are left out for every method), and the readout is the one of the cortical and hippocampal tests, applied at three scales: about 25 brain divisions, about 300 Allen structures and about 1,000 CCF-ME subregions (859–1,018 scored per brain). Two scores read the same table of cell types by regions from its two sides. *Cell-type placement* asks whether each cell type lands in the right regions: for every cell type with at least 200 scored cells, the overlap between the distribution of regions where it is placed and where it truly is (100% = exact). *Regional cell mix* asks whether each region receives the right cells: for every region with at least 50 scored cells, we compare the proportions of cell types among the cells placed into it with its true proportions using the Jensen–Shannon divergence, 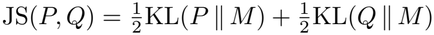 with 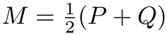 and logarithms to base 2, a symmetric distance between two mixtures that is 0 bits when the mixtures are identical and 1 bit when they share no cell type; a region that receives no cells scores 1. The measured positions read out the same way reach 96–97% placement and 0.005–0.007 bits, and a random layout 15–17% and 0.46–0.56 bits.

On mice 1 and 2 and on the Allen brain, our model places cell types into the right regions and gives each region a more faithful mix of cell types than LUNA-3D at every scale, and its margin widens as the anatomy gets finer (Figure S11e, f). At the CCF-ME scale, cell-type placement is 59.2, 58.5 and 60.5% against 52.5, 52.6 and 50.2%, and the mix error 0.137, 0.146 and 0.157 against 0.219, 0.220 and 0.286 bits (standard deviation over our four samples at most 0.16 points and 0.002 bits); on mouse 1 the margin in placement grows from 1.9 points across brain divisions to 6.0 across Allen structures and 6.7 across CCF-ME subregions. On mouse 1 the mix error of the CCF-ME subregions is lower for our model within every one of the 12 brain divisions (Figure S11d), for example 0.154 against 0.272 bits in the isocortex, 0.193 against 0.322 in the hippocampal formation and 0.094 against 0.197 in the thalamus. Panels a–c show where the errors lie in one coronal slice through the caudoputamen, which contains 145 CCF-ME subregions: every cell is drawn at its true position and colored by the mix error of its subregion, so light tissue is reconstructed with the right cells and dark tissue with the wrong ones (slice mean 0.079 against 0.103 bits). On mouse 3, cut sagittally and reconstructed from two coronal training brains, LUNA-3D places cell types better at every scale (54.2 against 51.9% at the CCF-ME scale), and the two models give each region a similar mix (0.183 against 0.181 bits at the CCF-ME scale).

**Figure S11:**
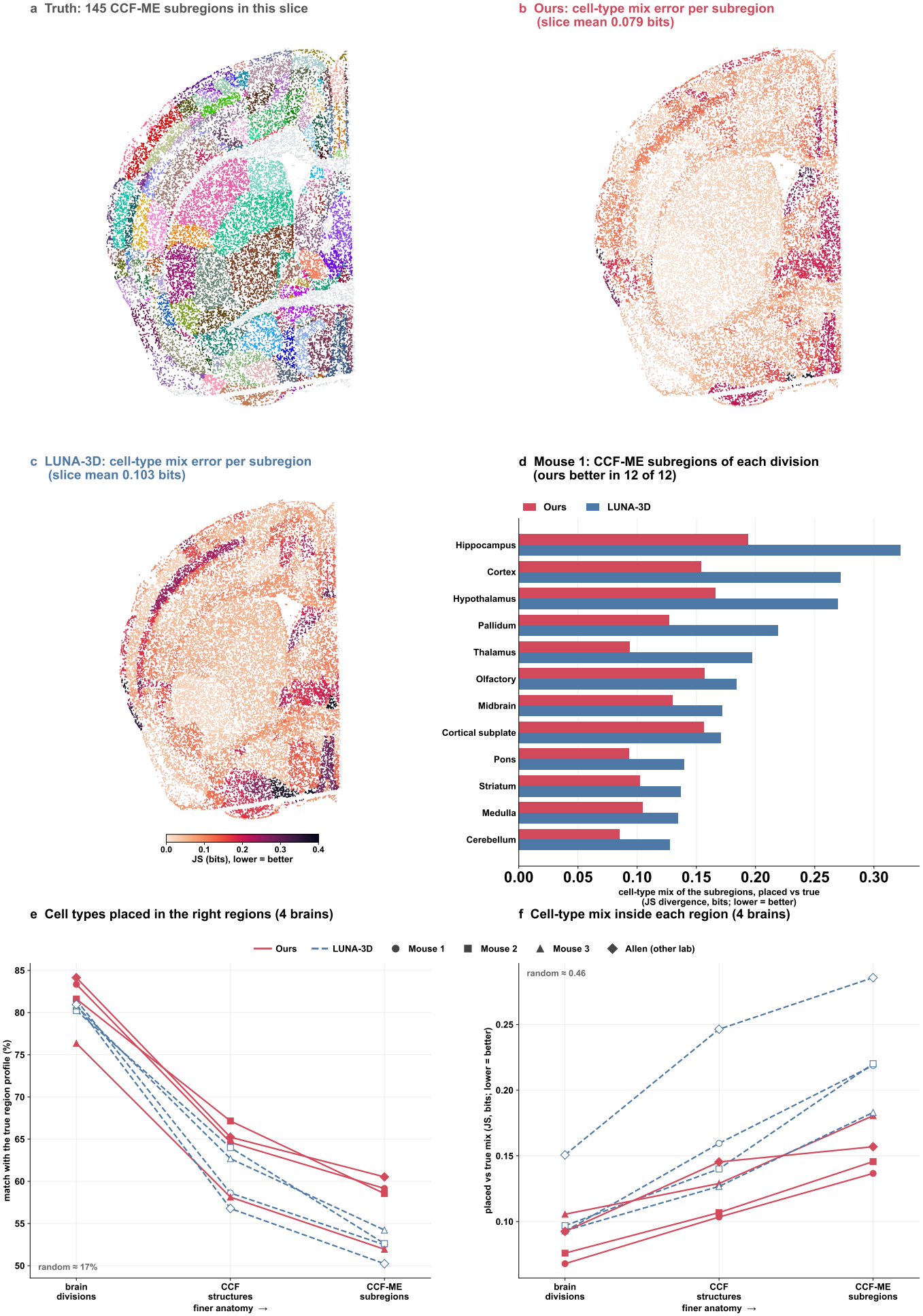
Cell types in the fine subregions of an independent, morphology-based atlas (CCF-ME). (a) A coronal slice of mouse 1 through the caudoputamen (± 40 *μ*m), cells at their true positions colored by CCF-ME subregion (colors only separate the 145 subregions). (b, c) The same cells, colored by the error of the cell-type mix that our model (b) and LUNA-3D (c) place into each subregion: Jensen–Shannon divergence between the placed and the true proportions of cell types, in bits (0 = the right mix; one color scale for both panels, light = low error, dark = high error). (d) Mouse 1: mean mix error of the CCF-ME subregions within each brain division. (e) Cell-type placement (overlap between the regions where each type is placed and where it truly is) and (f) mix error, from brain divisions through Allen structures to CCF-ME subregions, for the four test brains; solid, our model (mean of four samples); dashed, LUNA-3D.

#### Placed cells rebuild the expression maps of the measured genes

A reconstruction is read through genes: once cells have positions, every gene they carry acquires a three-dimensional map. We ask whether these maps are right for every measured gene of every test brain [15, 16], 1,122 genes in the mice and 477 in the Allen brain. For each gene we build a smoothed map of the mean expression per cell, log_2_(1 + count) of the measured MERFISH counts with a Gaussian of *σ* = 200 *μ*m on a 200 *μ*m grid, once from the cells at their measured positions and once from the same cells at their placed positions, so that the two maps differ only in where the cells are. The score is the Spearman correlation between the placed and the true map, which compares where a gene is high and where it is low, computed on the brain volume where the truth, LUNA-3D and each of our four samples all have at least five cells nearby, so that every method is scored on the same volume, and averaged over genes; a layout with the measured positions shuffled among the cells scores 0.00. Our placed cells rebuild the maps better than LUNA-3D in all four brains (Figure S12e, f): the mean correlation is 0.80, 0.74, 0.72 and 0.85 on mice 1–3 and the Allen brain, against 0.70, 0.64, 0.69 and 0.66 (our four samples agree within 0.001), and the map is better for 1,122 of 1,122, 1,122 of 1,122, 993 of 1,122 and 477 of 477 genes. The gain covers markers of every major structure [64, 16] (Figure S12d): on mouse 1, *Cux2* of the upper cortical layers 0.84 against 0.71, *Prox1* of the dentate gyrus 0.86 against 0.72, *Penk* of the striatum 0.87 against 0.74 and the inhibitory-neuron marker *Gad2* 0.84 against 0.71, with a median gain of 0.095 over all genes; on the brain from another laboratory the gap is largest, for example *Gad2* 0.86 against 0.59 and *Tcf7l2* 0.82 against 0.52. Gene domains such as a cortical layer, the dentate gyrus or the striatum keep their shape when cells keep their spread, whereas averaged positions pull cells toward the centres of their types, blur sharp domains into the space between them and shrink the brain, which is strongest for LUNA-3D on the Allen brain, reconstructed at 82, 87 and 105% of its true extent along the three axes.

**Figure S12:**
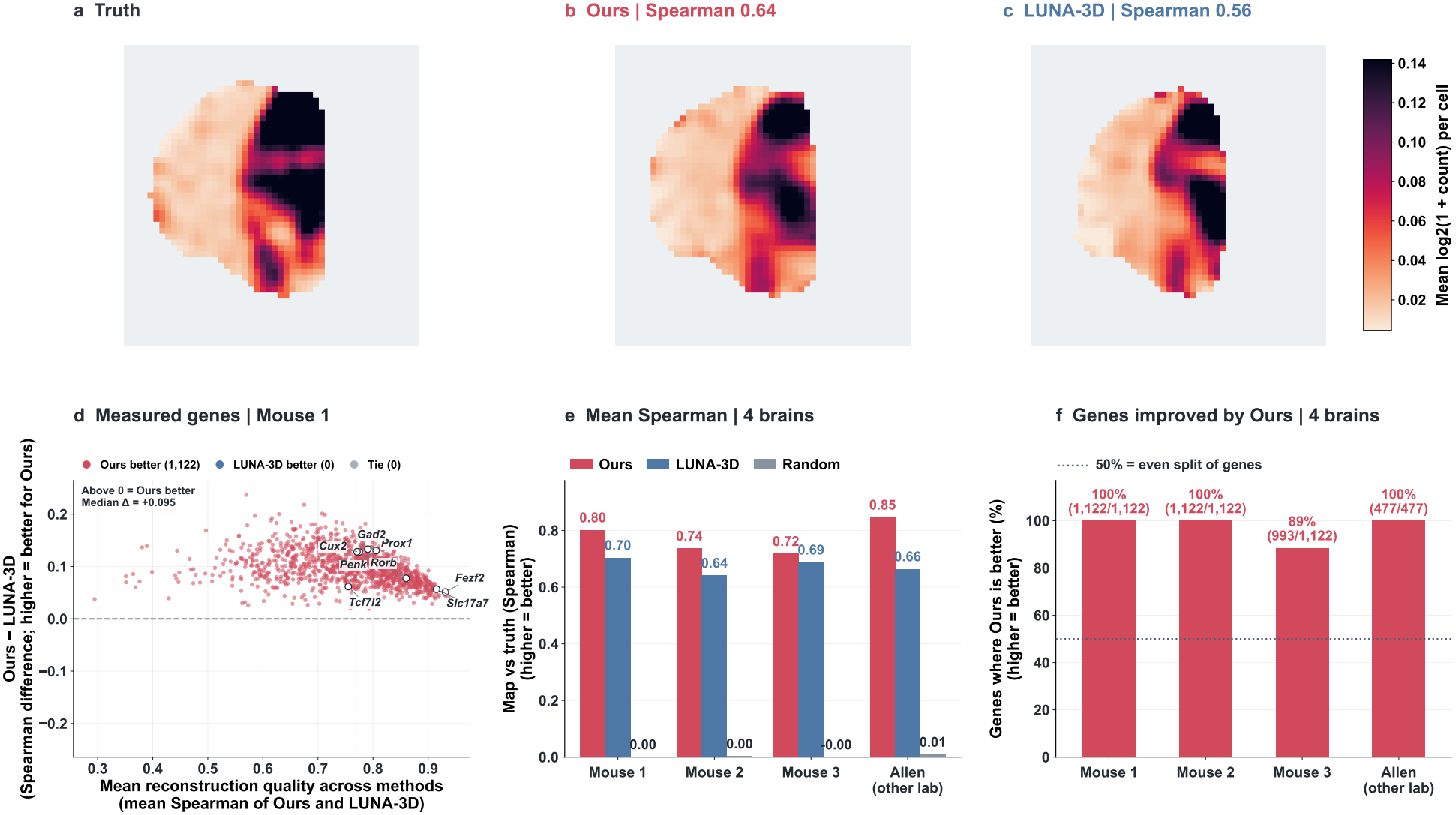
Gene-expression maps rebuilt from placed cells. (a–c) One coronal plane of the smoothed three-dimensional map of *Gata3* in mouse 1 (mean log_2_(1 + count) per cell), built from the cells at their measured positions (a), our placed positions (b; one sample) and LUNA-3D’s (c); one color scale; Spearman correlation with the true map in the titles. (d) Mouse 1, every measured gene (1,122): the difference between our and LUNA-3D’s correlation with the true map against their mean; above zero, our map is closer. (e) Mean correlation over genes for our model, LUNA-3D and random placement in the four test brains. (f) Share of genes whose map our model rebuilds better.

### S12 Dissociated cells return to their region of origin

#### Cells, models and ground truth

Every other test in this paper withholds coordinates that were measured, so that a reconstruction can be scored against them. The cells a reconstruction is built for have no coordinates to withhold: they are dissociated, profiled by a different technology from the spatial training data, with a different gene panel, capture efficiency and sampling of the brain. What such cells do carry is the region they were dissected from. The Allen whole-brain single-cell atlas [16] was collected donor by donor, each donor’s cells taken from a known anatomical region, and this gives a check on placement that needs no coordinates: a cell dissected from the thalamus should be placed in the thalamus. We draw five cohorts of 100,000 10x v3 cells at random from the atlas, without using region labels. Our model and LUNA-3D are trained on all five MERFISH brains (Zhang–Zhuang mice 1–4 and the Allen brain) and receive, for every cell, only its expression, projected into the 200-dimensional reference space of the spatial training cells by reference mapping with Symphony [63] on the 477 genes shared by the two platforms; neither model receives a region, donor, cell type or neighborhood. Our model places each cohort with three sampler seeds, and LUNA-3D is run three times, giving 15 runs per method. Both layouts are brought into the frame of the atlas by the same label-free procedure, a single rotation (with reflection) and translation, without scaling, fitted on the centers of 64 expression clusters. The dissection region and the cell-type labels were joined only after all layouts were frozen, and the practical-difference margins (2 percentage points for the hit rate, 25 *μ*m for the type geometry) were fixed before any outcome was seen.

#### Readouts

The 23 dissection regions of the atlas are each a fixed union of structures of the Allen Common Coordinate Framework. A placed cell is a hit when the plurality of its 15 nearest MERFISH atlas cells belongs to the region it was dissected from and the nearest of them lies within 500 *μ*m; the hit rate is averaged over the 23 regions, so that each region counts equally, and the 95% intervals of paired differences come from resampling whole donors. Three references give the number a scale: random placement reaches 3.6%; a measured MERFISH cell of an animal not in the atlas, scored through the same readout, reaches 76.4%, the realistic ceiling; and a reference that knows every cell’s subclass, which neither model is told, and places it at a random atlas position of that subclass reaches 27.4%. This last reference measures how much region information a placement carries beyond the identity of the cell. The type geometry is the sliced Wasserstein-2 distance between the cloud of placed 10x cells of a subclass and the cloud of MERFISH cells of the same subclass, averaged over the subclasses scored in every run. Our model reaches 37.9% against 28.5% for LUNA-3D, 10.5 points above the type-aware reference while LUNA-3D is 1.1 points above it, and shuffling our own positions among cells of the same subclass lowers our hit rate to 25.2% (that of LUNA-3D to 20.8%): our placement rests on the expression differences within a type, the transcriptional signature of where a cell of a given subclass lives, and uses them to choose among the regions its subclass occupies. Without the label-free pose, in the frame of its own training data, our model still reaches 35.6%, 7.0 points above LUNA-3D with the pose. The hit rate varies by less than 0.2 points across cohorts and seeds. Cell types are placed with the right geometry as well: the placed cloud of a subclass lies 402 *μ*m from its MERFISH counterpart for our model and 694 *μ*m for LUNA-3D, against 285 *μ*m between two independently measured MERFISH brains, and with location removed, comparing shape and spread alone, ours is still closer (281 against 428 *μ*m).

#### The placement does not depend on the scorer or on how cells are drawn

Two choices behind the hit rate could be questioned. The first is the scorer: membership of a region is decided through the nearest MERFISH atlas cells, a rule we built. We therefore also translate each placed cell into the Allen Common Coordinate Framework (the mean official coordinate of its 15 nearest atlas cells) and look it up in the official annotation volume, counting a hit when its dissection region lies within 500 *μ*m. The second is the sample: uniform draws inherit the uneven sampling of brain regions by the atlas, so we also draw five region-balanced cohorts with a fixed number of cells per dissection region, set from the MERFISH atlas before any placement was scored. Our model is ahead in all four combinations, by 8.5 to 9.9 points with every interval far from zero (Table S10, Figure S13a); it also beats the type-aware reference in every setting (37.9–39.7% against 27.3–27.9%), is higher for 17 of the 23 regions under the official annotation with either cohort design (Figure S13b), keeps the geometry of each cell type under both designs (405 against 700 *μ*m and 412 against 712 *μ*m, paired differences −291 *μ*m [−332, −252] over 185 types and −312 *μ*m [−363, −262] over 151 types; Figure S13c; these averages are taken per run, whereas the main text averages over the 185 types scored in all 15 runs, 402 against 694 *μ*m), and its hit rates change by less than one point when the distance tolerance is set to 250, 500 or 1,000 *μ*m (Figure S13d).

**Table S10:** The hit rate of the 10x placement under two scorers and two cohort designs (mean over 23 dissection regions, five cohorts and three sampler seeds; difference with 95% interval from resampling donors).

| **Scorer** | **Cohorts** | **Ours** | **LUNA-3D** | **Difference** |
|---|---|---|---|---|
| Atlas lookup (main text) | uniform | **37.9%** | 28.5% | +9.4 [8.5, 10.3] |
| Atlas lookup | region-balanced | **38.3%** | 28.4% | +9.9 [9.1, 10.8] |
| Official annotation | uniform | **38.9%** | 30.4% | +8.5 [7.6, 9.3] |
| Official annotation | region-balanced | **39.7%** | 30.3% | +9.4 [8.6, 10.2] |

**Figure S13:**
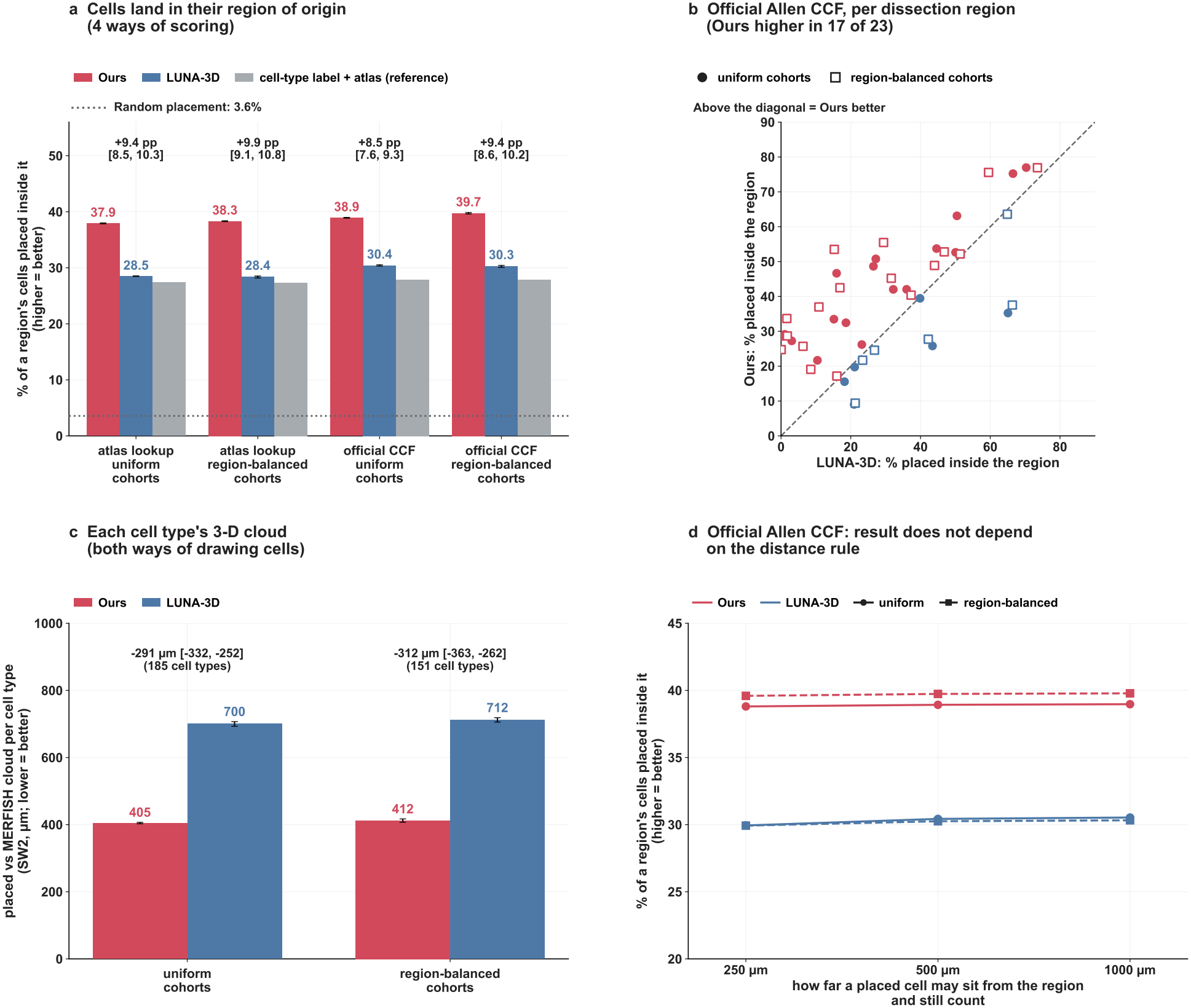
The 10x placement under other scorers and cohort designs (five cohorts × three sampler seeds per setting). (a) Share of each dissection region’s cells placed inside it, mean over 23 regions, under two scorers (nearest MERFISH atlas cells; official Allen annotation) and two cohort designs (uniform; region-balanced); error bars, standard deviation over cohorts; grey, a reference that places each cell at a random atlas position of its own cell type; dotted line, random placement; above each group, the paired difference with its 95% interval over donors. (b) Official annotation, one point per dissection region, ours against LUNA-3D; above the diagonal, ours is higher. (c) Sliced Wasserstein-2 distance between the placed 10x cells and the MERFISH cells of each cell type, mean over types, with the paired difference. (d) Official annotation with the inside-region tolerance set to 250, 500 or 1,000 *μ*m; solid, uniform cohorts; dashed, region-balanced cohorts.

**Table S11:** Maps of 645 genes outside the model input, rebuilt from placed 10x cells, under two ways of scoring the volume a layout leaves empty (Spearman correlation with the MERFISH reference map, mean over genes and 15 runs). Covered: share of the reference brain volume where a layout has at least five placed cells nearby. Shuffled: positions permuted among cells of the same subclass.

|  | **Own volume** | **Whole brain** | **Covered** |
|---|---|---|---|
| Two real brains (ceiling) | 0.68 | 0.63 | 96% |
| Ours | **0.51** | **0.31** | **58%** |
| LUNA-3D | 0.40 | 0.20 | 41% |
| Ours, shuffled within type | 0.48 | 0.30 | 58% |
| LUNA-3D, shuffled within type | 0.38 | 0.19 | 41% |
| Random placement | 0.00 | −0.00 | 74% |

### S13 Training and sampling cost

This section measures what a reconstruction costs and how the cost grows with the number of cells, which decides whether a generative model can serve atlases of millions of cells. We time our final recipes and LUNA-3D (settings in Supplementary Table S2) on the same GPU, one job at a time: an NVIDIA RTX PRO 6000 (96 GB) for the comparisons and an NVIDIA RTX A4500 (20 GB) where reseating is included, because our reseating engine runs on that GPU. Data are read into memory before each job, and data preparation and model loading are not counted. Table S13 lists the sampling of every test cell. Our sampler runs 50 steps; at each step it splits the *N* cells into windows of at most 20,000 cells and calls the network once per window, which makes 50 ⌈*N/*20,000⌉ calls. LUNA-3D samples its graphs of about 5,000 cells one at a time with 1,000 calls per graph, so its whole-test time is its mean time per graph times the number of graphs. Ours takes 1.2 seconds (embryo c) to 2.8 minutes (Allen brain) and LUNA-3D 44 seconds to 108 minutes, a ratio of 37 to 39 on every test; on the RTX A4500 the ratio is 21 to 25 (Table S15). The ratio has two factors. We take 20 times fewer steps: flow matching integrates a learned velocity field from noise to data, and half or twice our 50 steps worsen no measure by more than 4% (Table 2), while LUNA-3D’s released sampler walks all 1,000 steps of its diffusion chain. And one step costs 0.9 microseconds per cell for ours against 1.7 for LUNA-3D. Two sizes could drive the cost: the cells in the specimen and the cells in one network call. For the first, one model (mouse 1) samples *N* cells at the final settings: 0.89 seconds at 20,000 cells, 4.5 at 100,000, 46 at 1 million and 93 at 2 million (log-log slope 1.02); across the eight tests, from 25,755 to 3.7 million cells, the slope is 1.00 for both methods, so ten times the cells take ten times as long. For the second, Table S14 lists the time per call. Ours grows faster than the cells once a window is large (3.2 ms at 5,000 cells, 18.6 at 20,000, 213 at 80,000; slope 1.76 from 20,000 up), because full self-attention relates every pair of cells in the window; fixing the window at 20,000 cells is what keeps the cost of a whole specimen linear. LUNA-3D’s call grows about linearly (8.7 ms at 5,000 cells, 57.8 at 40,000; slope 0.91), so larger graphs would not lower its cost per cell. Sampling holds the features of all test cells on the GPU, so our peak memory grows with the test (1.9 to 13.3 GB, against 1.4 to 1.5 GB for LUNA-3D) and fits the 20 GB of the RTX A4500 on every test. Our full model also reseats the sample. On the RTX A4500, the sample, the one-call output that reseating reads and reseating itself (medians over the four draws of the final models) take 1.7 minutes on embryo c to 93 on the Allen brain; LUNA-3D’s sampling alone takes 2.1 to 303 minutes, 1.2 to 3.2 times longer. The reader that reseating uses is fitted once per training set (about 10 minutes for mouse 3) and is not counted.

Table S12 lists training. One update of ours holds 2 views of 10,500 to 15,700 cells on average; one of LUNA-3D holds 8 graphs of about 5,000. An epoch of ours is 246 views, 123 updates, on every dataset, so its time does not depend on the number of training cells: 11.9 to 17.7 seconds from 184,311 to 5.8 million cells (log-log slope −0.03). An epoch of LUNA-3D is one pass over all its graphs, 5 updates on the blocks and 146 on the Allen training set, so its time follows the cells: 1.1 to 22.3 seconds (slope 0.86). Time per epoch is the median over consecutive epochs after the first. A full training runs 360 epochs for ours (300 on the embryos) and 300 for LUNA-3D (400 on the blocks). Whole brains then take 1.19 to 1.77 hours with ours and 1.05 to 1.86 with LUNA-3D; on the RTX A4500, a mouse-2 epoch takes 65 seconds for ours and 63 for LUNA-3D. On the blocks and embryos ours takes far more updates (44,280 on the blocks and 36,900 on the embryos, against 2,000 and 2,400) and more time (1.33 to 1.71 against 0.12 to 0.17 hours). Per update ours is 1.2 to 1.9 times faster on every test (0.10 to 0.14 seconds against 0.15 to 0.25). To test whether LUNA-3D needs those updates, we also trained it for our number of updates with the same code and settings: 4,613 epochs on the embryos (36,904 updates) and 8,856 on the blocks (44,280). Its median per-cell error rises on all four tests (embryo d 85 to 90 *μ*m, embryo c 72 to 75, block A 878 to 922, block B 773 to 824), so the main tables report the shorter runs. At that length LUNA-3D takes 2.5 to 2.9 hours on the RTX PRO 6000, against 1.3 (embryos) and 1.7 hours (blocks) for ours at the same number of updates. Peak GPU memory in training is 4.5 to 12.7 GB for ours and 14.0 to 18.4 GB for LUNA-3D. REMAP, CeLEry, Tangram and novoSpaRc ran as one job per test, training and mapping together; Table S16 gives the wall-clock of the runs behind the main table, on servers shared with other jobs.

**Table S12:** Training cost on an NVIDIA RTX PRO 6000. Updates: optimizer steps per epoch. Training: seconds per epoch × epochs, ours 360 (embryos 300), LUNA-3D 300 (blocks 400). GPU: peak memory in GB. One training serves both embryos. Lower in bold.

|  |  | Updates / epoch |  | s / epoch |  | s / update |  | Training (h) |  | GPU (GB) |  |
|---|---|---|---|---|---|---|---|---|---|---|---|
| Test | Cells | Ours | LUNA-3D | Ours | LUNA-3D | Ours | LUNA-3D | Ours | LUNA-3D | Ours | LUNA-3D |
| Mouse 3 | 4,074,316 | 123 | 102 | 17.1 | 17.1 | **0.139** | 0.168 | 1.71 | **1.43** | **4.7** | 14.5 |
| Mouse 2 | 4,595,268 | 123 | 115 | **17.6** | 19.5 | **0.143** | 0.170 | 1.76 | **1.63** | **4.7** | 14.5 |
| Mouse 1 | 2,975,768 | 123 | 75 | 17.7 | **12.7** | **0.144** | 0.169 | 1.77 | **1.05** | **4.8** | 14.8 |
| Allen | 5,822,676 | 123 | 146 | **11.9** | 22.3 | **0.097** | 0.153 | **1.19** | 1.86 | **4.5** | 14.0 |
| Block A | 198,675 | 123 | 5 | 17.1 | **1.2** | **0.139** | 0.237 | 1.71 | **0.13** | **5.4** | 14.4 |
| Block B | 184,311 | 123 | 5 | 17.0 | **1.1** | **0.139** | 0.220 | 1.70 | **0.12** | **5.5** | 14.4 |
| Embryo c | 290,699 | 123 | 8 | 16.0 | **2.0** | **0.130** | 0.249 | 1.33 | **0.17** | **12.7** | 18.4 |
| Embryo d | 290,699 | 123 | 8 | 16.0 | **2.0** | **0.130** | 0.249 | 1.33 | **0.17** | **12.7** | 18.4 |

**Table S13:** Sampling every test cell on an NVIDIA RTX PRO 6000: 50 steps for ours, 1,000 for LUNA-3D. Calls: network evaluations. Time per call: ms, for a window of up to 20,000 cells (ours) or a graph of about 5,000 (LUNA-3D). Ratio: LUNA-3D over ours. GPU: peak memory in GB. Lower in bold.

|  |  | Calls |  | ms / call |  | Minutes |  |  | GPU (GB) |  |
|---|---|---|---|---|---|---|---|---|---|---|
| Test | Cells | Ours | LUNA-3D | Ours | LUNA-3D | Ours | LUNA-3D | Ratio | Ours | LUNA-3D |
| Mouse 3 | 1,585,782 | 4,000 | 317,000 | 18.1 | 8.8 | **1.2** | 46.3 | 38.4 | 7.9 | **1.5** |
| Mouse 2 | 1,227,408 | 3,100 | 245,000 | 17.9 | 8.7 | **0.93** | 35.6 | 38.4 | 6.4 | **1.5** |
| Mouse 1 | 2,846,908 | 7,150 | 569,000 | 18.4 | 8.7 | **2.2** | 82.9 | 37.9 | 13.3 | **1.5** |
| Allen | 3,739,961 | 9,350 | 748,000 | 18.2 | 8.6 | **2.8** | 107.5 | 37.9 | 4.1 | **1.4** |
| Block A | 184,311 | 500 | 37,000 | 16.8 | 8.7 | **0.14** | 5.4 | 38.5 | 1.9 | **1.4** |
| Block B | 198,675 | 500 | 40,000 | 18.6 | 8.7 | **0.15** | 5.8 | 37.6 | 2.0 | **1.4** |
| Embryo c | 25,755 | 100 | 5,000 | 12.0 | 8.9 | **0.02** | 0.74 | 37.1 | 2.6 | **1.5** |
| Embryo d | 40,198 | 150 | 8,000 | 12.6 | 8.9 | **0.03** | 1.2 | 37.7 | 3.0 | **1.5** |

**Table S14:** Time per network call against the cells in the call, on an NVIDIA RTX PRO 6000 (ms). Ours: the mouse-1 model. LUNA-3D: the mouse-2 model.

| Cells per call | 1,250 | 2,500 | 5,000 | 10,000 | 20,000 | 40,000 | 80,000 |
|---|---|---|---|---|---|---|---|
| Ours | 2.3 | 2.3 | 3.2 | 6.7 | 18.6 | 60.9 | 212.8 |
| LUNA-3D | 4.6 | 5.6 | 8.7 | 15.6 | 27.0 | 57.8 | – |

**Table S15:** Sampling with reseating on an NVIDIA RTX A4500, in minutes. Ours: the 50-step sample, the one-call output that reseating reads, and reseating; their total. LUNA-3D: its sampler. Ratios: LUNA-3D over our sample, and over our total. Lower in bold.

| Test | Sample | One call | Reseating | Ours, total | LUNA-3D | Ratio, sample | Ratio, total |
|---|---|---|---|---|---|---|---|
| Mouse 3 | 6.2 | 0.62 | 38.5 | **45.3** | 128.9 | 20.9 | 2.8 |
| Mouse 2 | 4.8 | 0.60 | 32.5 | **37.9** | 99.5 | 20.9 | 2.6 |
| Mouse 1 | 11.1 | 1.0 | 63.5 | **75.6** | 232.4 | 20.9 | 3.1 |
| Allen | 14.5 | 2.4 | 76.5 | **93.4** | 302.8 | 20.8 | 3.2 |
| Block A | 0.69 | 0.13 | 8.4 | **9.2** | 14.9 | 21.7 | 1.6 |
| Block B | 0.78 | 0.12 | 10.5 | **11.4** | 16.2 | 20.9 | 1.4 |
| Embryo c | 0.09 | 0.03 | 1.6 | **1.7** | 2.1 | 24.3 | 1.2 |
| Embryo d | 0.14 | 0.07 | 2.0 | **2.2** | 3.3 | 24.5 | 1.5 |

**Table S16:**
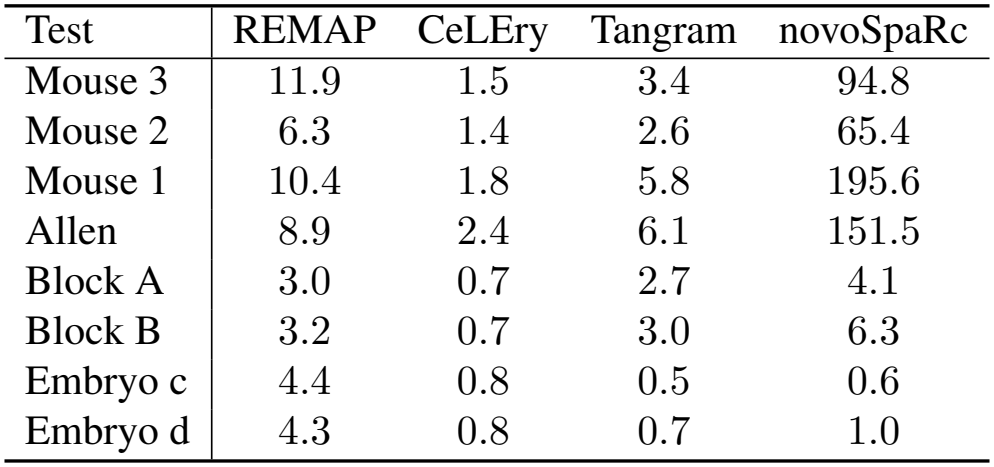
Other baselines: wall-clock of the runs behind the main table, one job per test, training and mapping together, in hours (novoSpaRc: CPU process-hours, summed over its parallel parts; settings in Supplementary Table S2).

